# Morphological profiling by Cell Painting in human neural progenitor cells classifies hit compounds in a pilot drug screen for Alzheimer’s disease

**DOI:** 10.1101/2023.01.16.523559

**Authors:** Amina H. McDiarmid, Katerina O. Gospodinova, Richard J.R. Elliott, John C. Dawson, Rebecca E. Hughes, Susan M. Anderson, Sophie C. Glen, Simon Glerup, Neil O. Carragher, Kathryn L. Evans

## Abstract

Alzheimer’s disease (AD) accounts for 60-70% of dementia cases. Current treatments are inadequate and there is a need to develop new approaches to AD drug discovery. We chose to develop a cell phenotype-based drug screen centred on the AD-risk gene, *SORL1*, which encodes the protein SORLA. Increased AD risk has been repeatedly linked to variants in *SORL1*, particularly those that confer loss of, or decreased, SORLA. This is consistent with the lower *SORL1* levels observed in post-mortem brain samples from individuals with AD. Consistent with its role in the endolysosomal pathway, deletion of *SORL1* is associated with enlarged endosomes in neural progenitor cells (NPCs) and neurons. We, therefore, hypothesised that multiparametric, image-based phenotyping would identify features characteristic of *SORL1* deletion. An automated morphological profiling assay (known as Cell Painting) was adapted to wild-type and *SORL1^-/-^* NPCs. This methodology was used to determine the phenotypic response of *SORL1^-/-^* NPCs to treatment with compounds from a small FDA/internationally-approved drug library (TargetMol, 330 compounds). We detected distinct phenotypic signatures for *SORL1^-/-^* NPCs compared to isogenic wild-type controls. Furthermore, we identified 16 approved drugs that reversed the mutant morphological signatures in NPCs derived from 3 *SORL1^-/-^* subclonal iPSC lines. Network pharmacology analysis revealed the 16 compounds belonged to five mechanistic groups: 20S proteasome, aldehyde dehydrogenase, topoisomerase I and II, and DNA synthesis inhibitors. Enrichment analysis confirmed targeting to gene sets associated with these annotated targets, and to pathways/biological processes associated with DNA synthesis/damage/repair, Proteases/proteasome and metabolism._Prediction of novel targets for some compounds revealed enrichment in pathways associated with neural cell function and AD. The findings suggest that image-based phenotyping by morphological profiling distinguishes *SORL1^-/-^* NPCs from isogenic wild-type lines, and predicts treatment responses that rescue *SORL1^-/-^*-associated cellular signatures that are relevant to both SORLA function and AD. Overall, this work suggests that i) a quantitative phenotypic metric can distinguish iPSC-derived SORL1^-/-^ NPCs from isogenic wild-type control and ii) phenotypic screening combined with multiparametric high-content image analysis is a viable option for drug repurposing and discovery in this human neural cell model of Alzheimer’s disease.

## 2 Introduction

Alzheimer’s disease (AD) is the most common form of dementia. AD is a neurodegenerative condition associated with a progressive decline in cognition, which is more pronounced than that observed in normal aging, ultimately leading to incapacitation and death (The World Health Organisation, 2022). The characteristic hallmark of AD is an accumulation of toxic amyloid-beta (Aβ) proteins which are produced by amyloidogenic processing of the amyloid precursor protein (APP). Aβ_40_ and Aβ_42_ form insoluble, extracellular plaques in the brain (Hampel et al., 2021). Aβ-related pathology, intracellular neurofibrillary tangles of hyperphosphorylated Tau protein (Serrano-Pozo et al., 2011) and neuroinflammation (Leng and Edison, 2021) are thought to lead to disruption of neuronal function and neurodegeneration in brain regions important for cognition (for example, the hippocampus). Neurodegeneration in these areas is associated with the overall impact on memory, thinking, language and learning capacity observed in AD.

Currently, there is no treatment which can cure AD. Modest, temporary relief from the cognitive impairment is achieved in a proportion of AD cases using acetylcholine esterase inhibitors (e.g. rivastigmine) (McGleenon et al., 1999; Polinsky, 1998) or the N-methyl-D-aspartate receptors antagonist memantine (Folch et al., 2018). No new drugs have been licensed for AD treatment since memantine in 2002. Only two new potential treatments for AD, the immunotherapeutics aducanumab and lecanumab, have been discovered in the last two decades (Ferrero et al., 2016; Herring et al., 2021; Lalli et al., 2021; van Dyck et al., 2022). Whilst aducanumab reduced plasma levels of Aβ_40_ and Aβ_42_ in a dose-dependent manner (Ferrero et al., 2016), it was not associated with significant improvements to cognition or function in participants (Lalli et al., 2021). Results from the trial of lecanumab suggest treatment has modest beneficial effects on disease progression over 18 months, but adverse outcomes such as reaction to infusion (26.4% of participants) and effusions or edema (12.6% of participants) (van Dyck et al., 2022) were also observed. This has raised questions around the efficacy and safety of these immunotherapies, which target Aβ aggregates in the brain. Thus, there is the opportunity to explore alternative targets/pathways and treatments, via drug discovery and repositioning, rather than solely focussing on clearance of Aβ plaques.

Genetic, clinical and functional analyses strongly support the involvement of the sortilin-related receptor 1 (*SORL1*, which encodes SORLA) in AD. In genome-wide association analyses, the associated variants in *SORL1* are largely non-coding and are expected to exert their effects via altering expression levels (Campion et al., 2019). Haploinsufficiency of *SORL1* is highly penetrant in AD, with 2% of early-onset AD cases being attributed to a rare *SORL1* variants (Holstege et al., 2017; Scheltens et al., 2021). *SORL1* haploinsufficiency in mini-pigs induced a cerebrospinal fluid biomarker profile identical to AD cases (Andersen et al., 2022). *SORL1* genetic risk variants have also been shown to predict endophenotypes of AD (decreased white matter fractional anisotropy and increased amyloid pathology in post-mortem brain) in individuals who were dementia-free at the time of the study (Felsky et al., 2014). It can be concluded, therefore, that reduced SORLA expression increases AD risk in both late/early-onset, sporadic and familial cases.

SORLA is a member of the domain VPS10p-domain receptor gene family that comprises five multifunctional neuronal proteins (Sortilin, SORLA, SORCS1-3). It acts as a receptor and a trafficking molecule, shuttling cargo between the plasma membrane, endosomes and the Golgi apparatus (Andersen et al., 2016). SORLA influences cellular response to growth factors, for example it potentiates the cellular response to brain-derived neurotrophic factor (BDNF) (Rohe et al., 2013). SORLA impacts amyloidogenic processes and thus Aβ levels (Andersen et al., 2016, 2005; Eggert et al., 2018; Schmidt et al., 2017). Decreased expression of SORLA is found in the brains of AD cases *post-mortem* (Dodson et al., 2006; Scherzer et al., 2004). Studies that looked prior to symptom onset suggest that this difference occurs prior to the development of clinical disease. In addition, *SORL1* brain expression was reduced in cognitively intact individuals with significant AD-like neuropathology, as well as in individuals with AD, in comparison to controls (Grear et al., 2009). Decreased *Sorl1* expression in mice accelerates Aβ production and senile plaque deposition (Andersen et al., 2005; Dodson et al., 2006), while overexpression of a human *SORL1* cDNA significantly reduced the amount of murine Aβ in wild-type mice and human Aβ in a mouse model carrying an AD-associated mutation in the human APP gene (Caglayan et al., 2014).

SORLA is implicated in endolysosomal trafficking of a number of molecules that are important to AD pathology, including APP (Hung et al., 2021; Knupp et al., 2020; Mishra et al., 2022; Young et al., 2015). It shuttles internalised APP from endosomes to the trans-Golgi network, decreasing the production of Aβ, and sorts Aβ peptides to lysosomes, where they are degraded. There is increasing evidence that endolysosomal pathways are important in the pathogenesis of neurodegenerative conditions, from both rare and common variant perspectives (Hu et al., 2015; Limone et al., 2022; Mishra et al., 2022 Szabo et al., 2022)). This is in keeping with recent studies demonstrating that loss of SORLA leads to morphological and functional abnormalities in organelles from this pathway (Hung et al., 2021; Knupp et al., 2020). Knupp et al 2020 showed that depletion of *SORL1* led to enlargement of early endosomes (independent of amyloidogenic APP processing) in SORLA depleted human iPSC-derived neural progenitor cells (NPCs) and neurons, but not microglia. Similarly, Hung et al., 2021 found that loss of *SORL1* in iPSC-derived neurons resulted in endosome, lysosome, and autophagy defects. In a minipig model of *SORL1* haploinsufficiency, endosomal enlargement was observed in neurons (Andersen et al., 2022). These findings suggest firstly that morphological phenotypes observed and quantified using fluorescence microscopy distinguish between SORLA depleted and wild-type cells, and secondly that this will be observed in NPCs, as well as neurons.

The finding of a phenotypic impact of SORLA loss in NPCs is not surprising given previous literature. For example, a developmental basis for AD has been hypothesised and some proteins implicated in neurodegeneration and AD-related pathology have important developmental functions (Arendt et al., 2017; Schor and Bianchi, 2021). In keeping with neurodevelopmental hypotheses of AD, atypical neurodevelopmental trajectories have been described in carriers of *SORL1* risk variants (Felsky et al., 2014) and there is evidence for increased proliferation of NPCs in mice lacking Sorla (Rohe et al., 2008). In addition, to being key to neurodevelopment, recent studies have shown that NPCs and hippocampal neurogenesis persist into at least the ninth decade of life in AD cases (Moreno-Jiménez et al., 2019; Tobin et al., 2019). Hippocampal neurogenesis was found to decline with age and, while levels were variable, it was decreased to a greater degree in patients with mild cognitive impairment and AD, with the extent of decline being correlated with disease severity. The persistence of NPCs into old age, and the correlation between decline and disease, suggest an important role for NPCs in the aging brain and in AD. In addition, NPCs are an ideal starting point for developing high-throughput drug screening as they have AD- and SORLA-relevant phenotypes that can be observed using image analysis and they can be grown in large quantities in a time and cost-effective manner.

Given the above considerations and the fact that previous studies showed morphological changes induced by loss of SORLA are similar in NPCs and neurons, we developed a phenotypic drug screen using *SORL1^-/-^* NPCs. In terms of the screening assay, we used a morphological profiling assay, Cell Painting, which facilitates hypothesis-free and relatively unbiased interrogation of phenotypic features, quantified from fluorescence microscopy images. It involves simultaneous application of multiple fluorescent dyes to label cellular organelles and the cytoskeleton. Following image acquisition, a CellProfiler image analysis pipeline is used to quantify >1000 cellular and subcellular morphological features based on stain intensity, texture, shape, area, pattern, granularity, adjacency and colocalization (Gustafsdottir et al., 2013). Dimensionality reduction, multivariate statistical analysis (including distance metrics and classification by machine learning) are then used to quantify (even subtle) phenotypic differences between mutant and wild-type cells. Subsequently, machine learning data analysis methods can be applied to identify compounds whose application causes reversion of the quantitative phenotypic signature (through any mechanism) of mutant cells towards that of wild-type cells. Such image-based multiparametric, computational approach may overcome some of the limitations of traditional single-readout end-point drug discovery assays (Chandrasekaran et al., 2021).

Here we show that morphological profiling by Cell Painting robustly classifies wild-type NPCs from those lacking SORLA. In addition, we performed a pilot drug screen with a small library comprising FDA/internationally-approved, biologically annotated small molecules. The screen yielded hits that reversed the mutant phenotype, demonstrating the potential of this assay to profile treatment response and identify compounds relevant to SORLA-related pathology and potentially AD.

## 3 Materials & Methods

### 3.1 Human induced pluripotent stem cell (hiPSC) line

Human induced pluripotent stem cell (hiPSC) line WTSIi004-B (also known as QOLG-1) was obtained from the European Bank for Induced Pluripotent Stem Cells (ebisc.org) with a material transfer agreement and access use agreement in place. The WTSIi004-B line was derived from fibroblasts obtained from a healthy male donor aged 35-39 at the time of primary cell collection. To revive hiPSCs from cryopreservation cells were thawed by incubation for 2 minutes at 37°C and retrieved by centrifugation (200 x *g* for 3 minutes) prior to resuspension in Essential 8 (E8, (A1517001, Life Technologies) medium with 1’RevitaCell. Colonies were maintained on vitronectin-coated plastic culture vessels, fed daily with E8 and incubated at 37°C (95% humidity with 5% CO_2_). Colonies were passaged at least twice after revival and prior to genome editing. All hiPSC cultures were confirmed as negative for mycoplasma on a monthly basis by submitting media supernatant for PCR testing by Institute of Genetics and Cancer Technical Services using a Lonza MycoAlert^®^ kit (LT07-218).

### 3.2 Generation of isogenic homozygous SORLA-depleted iPSCs by CRISPR-Cas9/Genome Editing

CRISPR-Cas9 guide RNAs (gRNAs) targeting exon 31 of the human *SORL1* gene (*SORL1*ex31) were designed using two open-source, computational tools; the Zhang Lab CRISPR Design website (https://crispr.mit.edu) and CHOPCHOP (https://chopchop.cbu.uib.no/). The final sequence was selected based on high specificity to the target site (*SORL1* exon 31) and low predicted off-target activity. Primer sequences targeting *SORL1* exon 31 in iPSCs were: CACCGTCGGTACCCGTCGCACACCC (top) and AAACGGGTGTGCGACGGGTACCGAC (bottom). The oligos were phosphorylated and subsequently cloned into the px458 vector, co-expressing the Cas9 endonuclease and GFP (RRID: Addgene_48138). Colonies of QOLG-1 wild-type parent hiPSCs were fed E8 2 hours prior to transfection. Cells were dissociated using Accutase (A6964, Merck), counted and 1×10^6^ cells were transfected using the Nucleofector II, Human Stem Cell kit (VPH-5012, Lonza) and the B-016 programme on the Amaxa Nucleofector II B device (Amaxa Biosystems). Pre-warmed E8 media (A1517001, Life Technologies) with 1 x revitacell (A2644501, Life Technologies) was added following nucleofection. The cells were then incubated at 37 °C for 5 min and gently added to a vitronectin-coated 6-well plate containing 2 ml of E8 containing 1 x revitacell. 2 μg of the Cas9 plasmid containing the gRNA were used in each transfection.

Forty-eight hours following transfection, colonies were dissociated with Accutase and retrieved from suspension by centrifugation (200g for 3 minutes). The cell pellet was resuspended in 700 μL Essential 8 Flex (E8 Flex, A2858501, Life Technologies) plus 10% CloneR™ supplement (05888, Stemcell Technologies). Single GFP-positive cells were sorted into individual wells of vitronectin-coated 96-well plates using a FACSJazz cell sorter (BD Biosciences). The single cells were allowed to expand into single-cell derived colonies by culturing for 48 hours (incubated at 37°C in 95% humidity with 5% CO_2_) in 100 μL E8 Flex plus 10% CloneR. A full medium change was then performed with E8 Flex plus 10% CloneR. After 24 hours, plates were topped up with 25 μL E 8 Flex plus 10% CloneR. After 24 hours a full medium change with El 8 Flex was repeated. Plates were then fed every 72 hours and monitored daily for the appearance of healthy colonies. After 10-14 days, colonies were passaged using 0.5 mM EDTA into a new 96-well. Once confluent, each clonal colony (subclone) was passaged again into two 96-wells to generate duplicate plates where one plate was used to generate cellular material for genomic DNA extraction and genotyping to confirm mutation of the *SORL1*ex31 target locus. The other plate was used to maintain/expand the respective subclones for selection based on the genotyping result.

### 3.3 *SORL1*ex31 Genotyping Assay and sequencing

QuickExtract^™^ DNA Extraction Solution (QE09050, Lucigen) was used to extract gDNA from the subclones for PCR. The gDNA samples were then amplified by polymerase chain reaction (PCR) using ReadyMix(TM) Taq PCR Reaction Mix (P4600,Sigma) containing 12.5μL Reaction Mix (P4600, Merck, 12.5uL), 2 μL 20 μM Primers 8.5 μL molecular grade and 2μL Quick Extract gDNA sample. Primers flanking the target site at *SORL1* exon 31: forward 5’ CTGCTCAGAGCTGTGCCAGT 3’ and reverse 5’ AGCCTTCCCTGGAGGTACAC 3’. A thermal cycler was used under the following programme: initial denaturation at 95°C for 1 min then 10 cycles of 95°C for 20 secs at 60°C-67°C (decreased by 1°C each cycle for 30 secs - 1min at 72°C for 1min with a further 25-30 cycles of 95°C for 20 secs at 50°C-57°C (decreased by 1°C each cycle) for 30 secs - 1min at 72°C for 1 min and a final extension at 72°C for 10 mins. Temperature was dependent on optimal annealing temperature of primers.

PCR products were visualised on a 1.5 % agarose gel (in 1X TBE Buffer) by loading 5 μL product with 10x loading dye alongside 10 μL of 1kb plus DNA Ladder. Products of the expected size were subject to sequence analysis. For Sanger sequencing the PCR product was cleaned (to remove excess primers and nucleotides) using ExoSAP-IT^™^ in the following reaction: 1 μL PCR Product, 1 μL ExoSAP-IT^™^ and 3 μL dH2O. Mutation of the *SORL1*ex31 locus was confirmed by sequencing of the target locus. For Sanger sequencing the PCR product was purified to remove excess primers and nucleotides using ExoSAP-IT^™^ using the following reaction: 1 μL PCR product, 1 μL ExoSAP-IT^™^ and 3 μL dH2O (60 mins at 37°C followed by 20 mins at 80°C. Sequencing was performed on the ExoSAP-IT^™^ treated product using the following reaction mix: 5 μL of ExoSAP-IT^™^ treated product, 1 μL of Big Dye v3.1, 1 μL of Big Dye sequencing buffer, 1 μL Primer (stock 3.2 μM) and 2μL dH_2_O using the following program: Initial denaturation at 96°C for 1 mins and then 30 cycles of 10 secs at 96°C at 5 secs at 50°C at 4 mins at 60°C. The purified products were precipitated with EDTA and ethanol for analysis on a 3130 or 3730 Genetic Analyser (Applied Biosystems). The same sequencing method was used to explore off-target effects of CRISPR-Cas9 editing in subclones with confirmed homozygous mutation at the *SORL1*ex31 locus were selected (n=3) along with crWT (n=3) subclones that were shown to carry no changes to the *SORL1*ex31 locus. Primers sequences flanking the off-target sites were: S1EX31Tch2 forwards 5’ TATGGGCTTCAAAGGGGAGG 3’ and reverse 5’ TTATGCTGCATCTCCCCAGG 3’, S1EX31Tch11 forwards 5’ GAGAAGAGTGCTGGGACTGT 3’ and reverse 5’ TGGATCCCTACTGTATGGCC 3’, S1EX31Tch1 forwards 5’ GAGAGAAAATGCAGCCAGGC 3’ and reverse 5’ TGTGTTTCTCCCTTCCCCAC 3’, S1EX31Tch22 forwards 5’ CACAAAATGCCCACCCACAG 3’ and reverse 5’ TAGTAGAGAGGGGCTTTCGC 3’, S1EX31Tch11_2 forwards 5’ GGGTCTGGTGCCTGGAAG 3’ and reverse 5’ CAAGATTGCGCCACTGTACT 3’, S1EX31Tch19 forwards 5’ GAGTCCCAGAGCCACGATC 3’ and reverse 5’ TGCAGCATTAACAGAGCAGG 3’, S1EX31Tch10 forwards 5’ AATTGCCTACCTCCTCCACC 3’ and reverse 5’ GCCACGTTCTTCTGTCTGTC 3’, S1EX31Tch11_3 forwards 5’ GCAGAGTGGTGACGGACA 3’ and reverse 5’ CCCCTGAGAATGGAGGACC 3’, S1EX31Tch1_2 forwards 5’ TCTCAGCCCGGATAAGTAGG 3’ and reverse 5’ GTGTGAGTGGCCGAGAGTT 3’.

### 3.4 Karyotyping

After CRISPR-Cas9 editing, each *SORL1*ex31 (n=3) knock-out subclone, crWT (n=2) and the QOLG-1 wild-type parent line (n=1) were karyotyped using a commercial service that provides information on chromosomal changes >1Mb resolution (KaryoStat Assay, ThermoFisher). Briefly, confluent hiPSCs colonies were detached from one well of a 6-well vitronectin-coated culture vessel using 0.5uM EDTA and retrieved by centrifugation (200g for 3 mins) to generate a cell pellet for karyotypic analysis.

### 3.5 Immunoblotting

Cells were lysed in ice cold 1% Triton lysis buffer [20 mM Tris–HCl pH 8.0, 10 mM EDTA, 1% Triton X-100 and 1x protease inhibitor cocktail (5892970001, Roche)] and protein concentration was measured using Bio-Rad BSA protein assay (5000116, Bio-Rad). Protein lysates were loaded on NuPAGE Tris–acetate 3–8% precast gels (EA03752BOX, Life Technologies) and subject to electrophoresis at 150 V for 1.5 h. Gels were transferred onto PVDF membranes at 30 V for 1.5 h. Membranes were blocked in 5% milk in 0.2% Tween-20 in TBS for 1 h at room temperature and probed with primary antibodies against SORLA (1:4000; 611680, BD Transduction Labs) and GAPDH (1:10,000; MAB374, Merck) diluted in blocking solution overnight at 4 °C. After washes (3 × 10 min) in 0.2% Tween-20 in TBS, membranes were incubated with secondary HRP-conjugated antibodies diluted 1:10,000 in blocking solution for 1 h at room temperature. After another three washes with TBS-0.2% Tween-20, blots were visualised using the Pierce ECL Plus Western Blotting Substrate (11527271, Thermo Scientific) and exposed using autoradiography film. Protein lysate obtained from HEK293 cells transfected with a plasmid overexpressing a human *SORL1* cDNA was used as a positive control.

### 3.6 Derivation of Neural Precursor Cells (NPCs)

Colonies of hiPSCs from *SORL1*ex31 sublones, crWT and parent wild-type were cultured until 70-80% confluent with no visible differentiation before accutase dissociation and retrieval from suspension by centrifugation at 200g for 3 mins. Neural induction was achieved using the StemDiff Neural Induction Kit (IM, 08581, Stem Cell Technologies) and following the embryoid-body (EB) protocol as per manufacturer’s instructions. Briefly, an aggrewell-800 plate (34811,Stem Cell Technologies) was prepared by washing with anti-adherence solution (07010,Stem Cell Technologies). The aggrewell-800 plate is a 24-well culture plate each well contains 300 microwells used for the creation of EBs from hiPSCs. 3×10^6^ cells from each hiPSC line (n=3 *SORL1*ex31 knock-out subclones, n=2 crWT and n=1 parent wild-type), were resuspended in 1mL IM and added to individual wells of the Aggrewell-800 plate. The plate was then subjected to centrifugation (100g for 3 mins) to deposit the cells in the microwells (~10,000 cells/microwells, 3×10^6^ total cells per well) and fed daily with IM (half media change) for 5 days. EBs were replated on matrigel-coated 6-well plates, and fed daily (full media change) with IM and monitored for emergence of neural rosettes. After 7 days, EBs were treated for 1 hour with neural rosette selection reagent (05832,Stem Cell Technologies) prior to collection and replating into matrigel-coated 12-well plastic culture plates to establish passage 0 (p0) NPCs. On day 14, NPCs were passaged for the first time (p1) into Neural Progenitor Media (NPM) and allowed to expand to 100% confluence. At this point, NPCs were expanded to p2 and p3 prior to banking and cryopreservation in liquid nitrogen in vials of 2×10^6^ cells in NPM with 10% (v/v) DMSO. The derivation was performed in technical duplicate. NPC cultures were confirmed as negative for mycoplasma by routinely submitting media supernatant for testing.

### 3.7 Immunocytochemistry

NPCs were fixed using 4% PFA for 20 mins after 48h in culture in an optical 384-well plate. NPCs were washed twice with 1x phosphate buffered saline (PBS) for 5 mins per wash. Permeabilisation was achieved by incubation for 1h in freshly prepared blocking solution (PBS with 5% (v/v) normal donkey serum (D9663, Sigma) and 0.3% (v/v) Triton X-100) followed by incubation overnight at 4°C with primary antibodies (10μL solution per well). Primary antibodies were diluted in blocking solution. Anti-Sox2 (1:500 AB5603, Chemicon) primary antibody was used as a stem cell marker and Anti-Nestin (1:500, MAB5326, Chemicon) as a marker for neural stem/progenitor/precursor cells. NPCs were washed twice in 1xPBS before incubation in blocking solution (50μL per well) for 30 minutes. Fluorescent-tagged secondary antibodies were diluted to 1:500 in blocking solution. NPCs were incubated with Alexa-488 Donkey Anti-Rabbit,(ab150077, Abcam and Alexa-594 Donkey Anti-Mouse (ab150116, Abcam) for 2 hours protected from light to preserve fluorescence. After three washes in 1xPBS, the DNA counterstain DAPI (50μL per well: 300μM) was applied for 10 minutes and then NPCs were washed three further times in 1xPBS before finally adding 1xPBS (50μL per well) and a foil plate seal applied prior to imaging using florescence detection microscopy. All steps carried out at room temperature unless stated otherwise.

### 3.8 Drug screening

*SORL1^-/-^* subclones (n=3) and parental WT (n=1) NPCs were seeded (5000 cells per well) in 384-well optical-bottom microplates coated with matrigel in NPM. A total of 9 plates were seeded, 3 per *SORL1*ex31 knock-out subclone. In 32 wells per plate, parental WT NPCs were seeded to serve as reference samples for phenotypic rescue and as within-plate positive controls. Plates were incubated for 24h in a tissue culture incubator at 37°C, 95% humidity and 5% CO_2_ to allow time for setting and recovery before the addition of compound treatments. The TargetMol Annotated Anti-Cancer library (L2110, 330 compounds, Supplementary Information Table 1) was supplied in assay-ready plates and thawed fresh on the morning of use. Serial dilution of the stock library was achieved using Biomek FX liquid handling system to generate compound plates at working concentrations suitable for in-well dilution in the NPC screening plates at three concentrations: 100nM, 300nM and 1μM. Thus, each subclone was screened across three concentrations in three plates (totalling 9 plates). All NPCs were cultured for 24h after compound treatment before fixation.

### 3.9 Cell Painting assay

NPCs were fixed in 4% PFA for 20 mins at room temperature. NPCs were washed twice with 1x PBS for 5 mins per wash. Fluorescent dyes were diluted in 1% (v/v) BSA solution in 1xPBS with 0.1%(v/v) Triton X-100 to prepare the Cell Painting staining solution for labelling the nucleus (DAPI, 1μg/mL; ThermoScientific^™^ cat no. 62248), nucleolus (SYTO14, 3μM; Invitrogen^™^ cat no. S7576), Golgi apparatus/plasma membrane (Wheat Germ Agglutinin, 2μg/mL, Invitrogen^™^ cat no. W11262), cytoskeleton (Phalloidin, 1:500, Abcam cat no. ab176757) and mitochondria (MitoTracker Deep Red, 600nM; Invitrogen^™^ cat no. M22626) (Table 1). The Cell Painting Staining solution was applied to each well of the 384-well optical-bottom microplates (20μL per well) and incubated for 1h at room temperature protected from light to preserve fluorescence. Cells were washed twice with 1xPBS and then 1xPBS added to each well (50μL per well) and a foil plate seal applied prior to imaging using florescence detection microscopy.

**Table 1.**
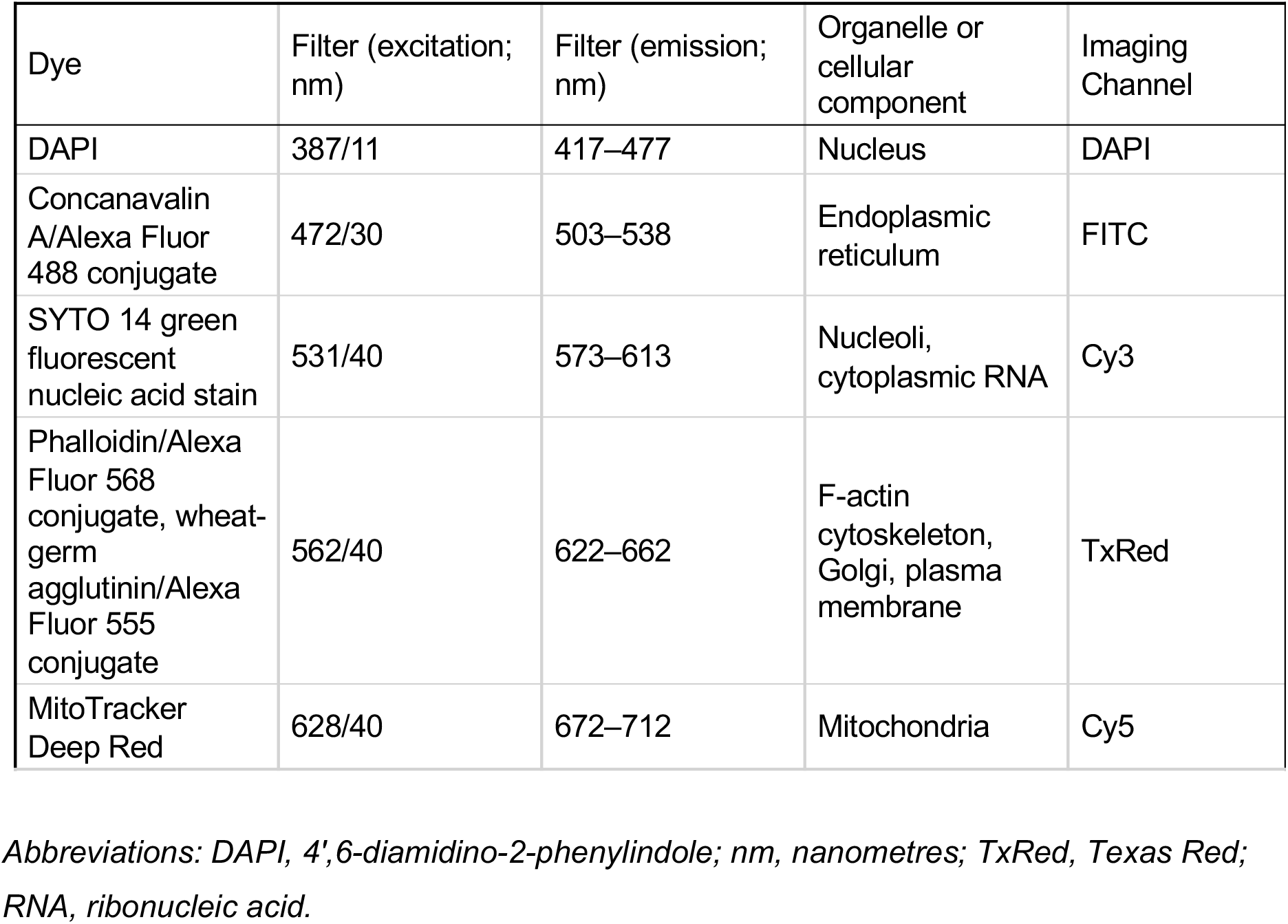
Cell Painting assay reagents with conjugated fluorophore emission/excitation spectra filters, cellular components label and ImageXpress^™^ XL channel used for image acquisition correeponding to each dye.

| Dye | Filter (excitation; nm) | Filter (emission; nm) | Organelle or cellular component | Imaging Channel |
| --- | --- | --- | --- | --- |
| DAPI | 387/11 | 417–477 | Nucleus | DAPI |
| Concanavalin A/Alexa Fluor 488 conjugate | 472/30 | 503–538 | Endoplasmic reticulum | FITC |
| SYTO 14 green fluorescent nucleic acid stain | 531/40 | 573–613 | Nucleoli, cytoplasmic RNA | Cy3 |
| Phalloidin/Alexa Fluor 568 conjugate, wheat-germ agglutinin/Alexa Fluor 555 conjugate | 562/40 | 622–662 | F-actin cytoskeleton, Golgi, plasma membrane | TxRed |
| MitoTracker Deep Red | 628/40 | 672–712 | Mitochondria | Cy5 |

*Abbreviations: DAPI, 4',6-diamidino-2-phenylindole; nm, nanometres; TxRed, Texas Red; RNA, ribonucleic acid.*

### 3.10 Automated fluorescence microscopy

Fluorescence imaging was performed using ImageXpress Confocal (Molecular Devices, USA) and accompanying MetaXpress Software (Molecular Devices, USA) with a robotic plate handling arm (Harmony, PAA: Peak Analysis and Automation, UK). User-defined parameters (such as exposure time and laser off-set) were optimised based on a sample of 10 wells randomly distributed across each of the 384-well microplates (781091, Greiner) and such parameters kept constant between plates thereafter as to avoid variation in intensity values across screening experiment. For immunocytochemistry, images were acquired for 3 fluorescent channels (filters: DAPI, FITC and Texas Red) at 20x magnification. For Cell Painting, images were acquired for 5 fluorescent channels (filters: DAPI, FITC, Cy3, Texas Red and Cy5, Table 1) at 20x magnification. For each well, 4 fields of view were captured. Illumination correction was achieved using a within-instrument tool to adjust for small variations in sample illumination for each field of view. Illumination correction was applied at time of image acquisition to correct intensity values according to an illumination correction function calculated using MetaXpress Software.

### 3.11 Image-analysis

To quantify the efficiency of neural induction of hiPSCs to NPCs, the proportion of cells expressing or co-expressing Sox2/Nestin as NPC marker(s) was quantified by CellProfiler (v4.1.2) using a pipeline applied to images acquired from immunocytochemistry. The analysis pipeline targeted the region of interest for each cell (defined by segmenting DAPI counterstained nuclei). The proportion of cells expressing or co-expressing markers was counted by optimising fluorescence intensity threshold parameters to select and validate Sox2+ and Nestin+ objects. From this, the percentage change in marker expression relative to the experimental controls was calculated to determine comparative changes in cell populations on different genetic backgrounds/experimental conditions. Exposure time, imaging plane and reference levels were calibrated within each experiment and kept constant with minor adjustments to the threshold set to eliminate background signal (adjustments made with respect to negative (primary antibody-free) controls.

Cell Painting image analysis was performed using CellProfiler (cellprofiler.org, v4.1.2). The region of interest, i.e. a cell, was targeted by segmenting nuclei from images acquired in the DAPI channel to generate a nuclear mask. Using the nuclear mask as a reference point, segmentation of the cell body using the plasma membrane and cytoskeleton stain (TxRed channel) was used to create a mask for the cytoplasm. Those masks were then used to isolate cellular objects within each image that were subjected to multiparametric analysis to quantify morphological features per object per image from the Cell Painting assay using a bespoke pipeline (CellProfiler, v4.2.1) which included, but was not limited to, measures of stain colocalization, object adjacency, size, shape, area, texture, radial distribution, granularity and intensity.

### 3.12 Data analysis and statistics

CellProfiler analysis was submitted as an array job using scripts data staging analysis and data destaging GridEngine scripts generated by cptools2 (https://github.com/CarragherLab/cptools2), on “Eddie” the high-performance computing cluster at University of Edinburgh. Analysis was performed in 100-image chunks, and the resulting datasets concatenated using Python. The quantitative multivariate object-resolution datasets derived from image analysis (by MetaXpress or CellProfiler) were exported as .csv files and stored on University of Edinburgh datastore prior to upload to the HC analysis module included with StratoMineR (Core Life Analytics). During data pre-processing, metadata was defined within the dataset and quality control carried to remove images with poor focus, illumination artefacts and debris. For image quality control, image quality metrics were analysed using MeasureImageQuality.py (CellProfiler, v4.2.1). Object size and shape measures can also be used as indicators of object segmentation performance and in-well debris/other imaging artifacts. Poor quality images were removed due to sparsity/high confluency, clumped cells leading to mis-segmentation during image-analysis, poor focus, in-well debris or illumination artefacts which could not be corrected for since these issues impact the precision of object-level quantification and the aggregation of the datasets to image-level medians. Positive and negative control image sets were visually inspected since these image sets form the test and training classes used for classification by machine learning. Redundant variables with Pearson correlation co-efficient >0.99 (*p* < 0.05) were removed. Variables (known as features) were selected, transformed to ensure normality of distribution across all features and normalised according to the sample median to control for inter-plate variation. Feature scaling was achieved by calculating robust Z-scores on a per-plate basis.

Principal component analysis was used for dimensionality reduction and factor analysis applied to extract components important for explaining variation with reference to the samples and controls. Variables causing singularity were eliminated. Factor analysis was applied with respect to with the DMSO vehicle-treated *SORL1^-/-^* and DMSO vehicle-treated WT NPCs. This was performed using oblique (oblimin) factor rotation method and factor scores calculated by ten Berge method. Using Kaiser’s criterion and examination of a Scree plot, a set of 50 Principal Components (PCs) were selected for analysis. Euclidean distance and Bray-Curtis Dissimilarity metrics were used to test for phenotypic separation of *SORL1^-/-^* and wild-type controls on the 50 PCs. Selected PCs were also used as quantitative signatures in a machine learning classification model for hit selection. A Random Forest (RF), neural network (NN) and supported vector machine (SVM) classifier algorithms were trained on the vehicle (DMSO) treated positive and negative control images (20% test versus 80% training set) and then each model used to classify sample (compound treatment) images as either *SORL1*ex31 knock-out or wild-type. NN resulted in greatest phenotypic separation of morphological signatures associated with genotypes and compound treatment, so was therefore selected as the method for hit selection in this study. All algorithms were applied to detect objects with morphological profiles similar to the positive control. Briefly, the positive control used phenotypic profiles from the DMSO vehicle-treated isogenic parent wild-type NPCs and this was used as the focus class in the classifier in order to determine which compound-treated *SORL1*ex31 knock-out NPCs (n = 3 subclones) classified as similar (>50.5% likelihood) to the wild-type positive control following treatment. Compounds that induced phenotypic morphological profiles in the *SORL1*ex31 knock-out that lead to increased likelihood of classification as an isogenic wild-type parental control were selected as hits (provided the result replicated in all subclones). Graphical visualisations were produced using Plotly in R (v4.0.2, www.r-project.org).

### 3.13 Transcriptomic profiling

An RNAseq expression dataset which profiled expression of 3 successive passages of wild-type NPCs (p2, p4 and p5) from hiPSC line WTSIi004-B (also known as QOLG-1) was generated by Marie-Thérèse El-Daher and Yanick Crow (Medical Research Council Human Genetics Unit, Institute of Genetics & Cancer, University of Edinburgh). The dataset was kindly shared with us for filtering irrelevant genes from the network analysis and background correction during enrichment analysis. Marie-Thérèse El-Daher derived NPCs using the StemDiff Neural Induction Kit (IM, 08581, Stem Cell Technologies) following the embryoid-body (EB) protocol as per manufacturer’s instructions as described above. Integrity of total RNA was analysed by Fragment Analyser Automated Capillary Electrophoresis (Agilent Technologies Inc, 5300) using the Standard Sensitivity RNA Analysis kit (DNF-471-0500) with DNA contamination assessed by Qubit dsDNA HS Assay kit (Q32854). RNA concentration was quantified by Qubit RNA HS Assay kit (Q32855) with 2.0 Fluorometer (Thermo Fisher Scientific Inc, Q32866). Sequencing libraries were generated from 50-100ng total RNA starting material using QuantSeq 3’mRNA-Seq Library Prep kit (FWD) for Illumina (Lexogen Inc, #015). After first and second strand synthesis, cDNA libraries were amplified for 18 cycles with adaptor and index sequences incorporated for parallel sequencing and bead-purification used to remove contaminating primers or adapter-dimers. Libraries were quality controlled using the Agilent Bioanalyser with the DNA High Sensitivity kit, and cDNA concentrations quantified by Qubit dsDNA High Sensitivity assay. Single read (1×75bp) sequencing was performed on the NextSeq 550 platform (Illumina Inc, SY-415-1002) using the NextSeq 500/550 High-Output v2.5 kit (20024906). Raw data was converted to FASTQ format using Bcl2fastq2 (Illumina, v2.17.1.14). QuantSeq was used to map and quantify mRNA expression level using the Bluebee genomics analysis platform (www.bluebee.com/quantseq).

### 3.14 Network analysis

Validated and predicted target lists for each compound hit were generated using database searches. Experimentally validated compound-protein interactions were generated by searching the compound names (grouped by mechanism of action according to published functional annotations) in the STITCH database (STITCH: https://stitch.embl.de). The top 10 physical interactors were used for visualisation of the networks targeted by each class of compounds. For enrichment analysis, the top 20 interactors (physical subnetwork only) were exported to the STRING database (STRING; https://string.embl.de). The networks were then filtered to include only nodes (proteins) and validated edges (interactions between proteins) with experimental and database evidence to validate them. Network lists were also filtered to eliminate any proteins that did not show expression at transcript level in QOLG-1 wild-type NPCs. A predicted target list for each compound hit was generated using a similarity ensemble approach (SEA) search (sea.bkslab.org, (Keiser et al., 2007)). The SEA search uses the SMILES string for each compound to query possible compound-protein interactions not limited to experimental annotations. By exploring the pharmacological space corresponding to a particular chemical structure it is possible to computationally predict possibly novel protein targets. Those putative interactions predicted by SEA were ranked according to significance, which is a measure of the probability of the predicted interaction, as well as Tanimoto coefficient (MaxTC), a measure of structure-based similarity. Predicted molecular targets were removed if they did not relate to a human protein, and those with a Z-score <1.96, MaxTC ≥0.4 and significance *p* ≤ 10^-15^ selected for further analysis. Protein-protein interaction (PPI) enrichment analysis was used to determine whether proteins in each network was greater than expected due to chance (*p* < 0.05).

### 3.15 Enrichment analysis

Enrichment analysis was conducted using the enrichment tool included in the STRING platform. This was used to determine which predicted and validated network components (i.e. proteins/genes) were overrepresented in biological pathways (KEGG database, https://www.genome.jp/kegg/), biological processes and molecular functions (Gene Ontology; http://geneontology.org/). The STRING database was used to highlight nodes representing genes associated with KEGG/GO terms within networks.

## 4 Results

### 4.1 Generating SORLA-depleted hiPSCs for derivation of neural progenitor cells

We used CRISPR/Cas9 genome editing to target exon 31 of the *SORL1* gene in a human iPSC line (Figure 1A). Five independent subclones carrying homozygous mutations predicted to lead to the introduction of a premature stop codon were generated. Loss of SORLA expression in *SORL1^-/-^* hiPSCs was shown by western blot (Figure 1B). Three *SORL1^-/-^* subclones were selected for use in this study along with one parental wild-type (WT) and two isogenic CRISPR wild-type (crWT) lines. NPCs were derived from the above lines and SORLA depletion confirmed by western blotting (Supplementary Figure 1). Nine off-target genomic loci predicted for the sgRNA were subjected to sequence analysis, which revealed no mutations at those sites in the edited hiPSCs. Karyotypic normality was also confirmed before and after gene-editing. The proportion of Nestin+/Sox2+ cells was quantified using fluorescence microscopy in both WT and edited lines. Neural induction was highly efficient and comparable in the two lines (WT median proportion Nestin+ cells = 100.00%; *SORL1^-/-^* median proportion Nestin+ cells = 100.00%, Mann-Whitney *U* = 3791, *p* = .7122, Figures 1C and D). However, the proportion of Sox2+ nuclei was significantly lower in *SORL1^-/-^* NPCs (WT median proportion Sox2+ nuclei = 79.69%; *SORL1^-/-^* median proportion Sox2+ nuclei = 69.82%, Mann-Whitney *U* = 1432, *p* < .0001, Figure 1E) suggesting there may be differences in pluripotency of NPCs derived from SORLA-depleted hiPSCs.

**Figure 1.**
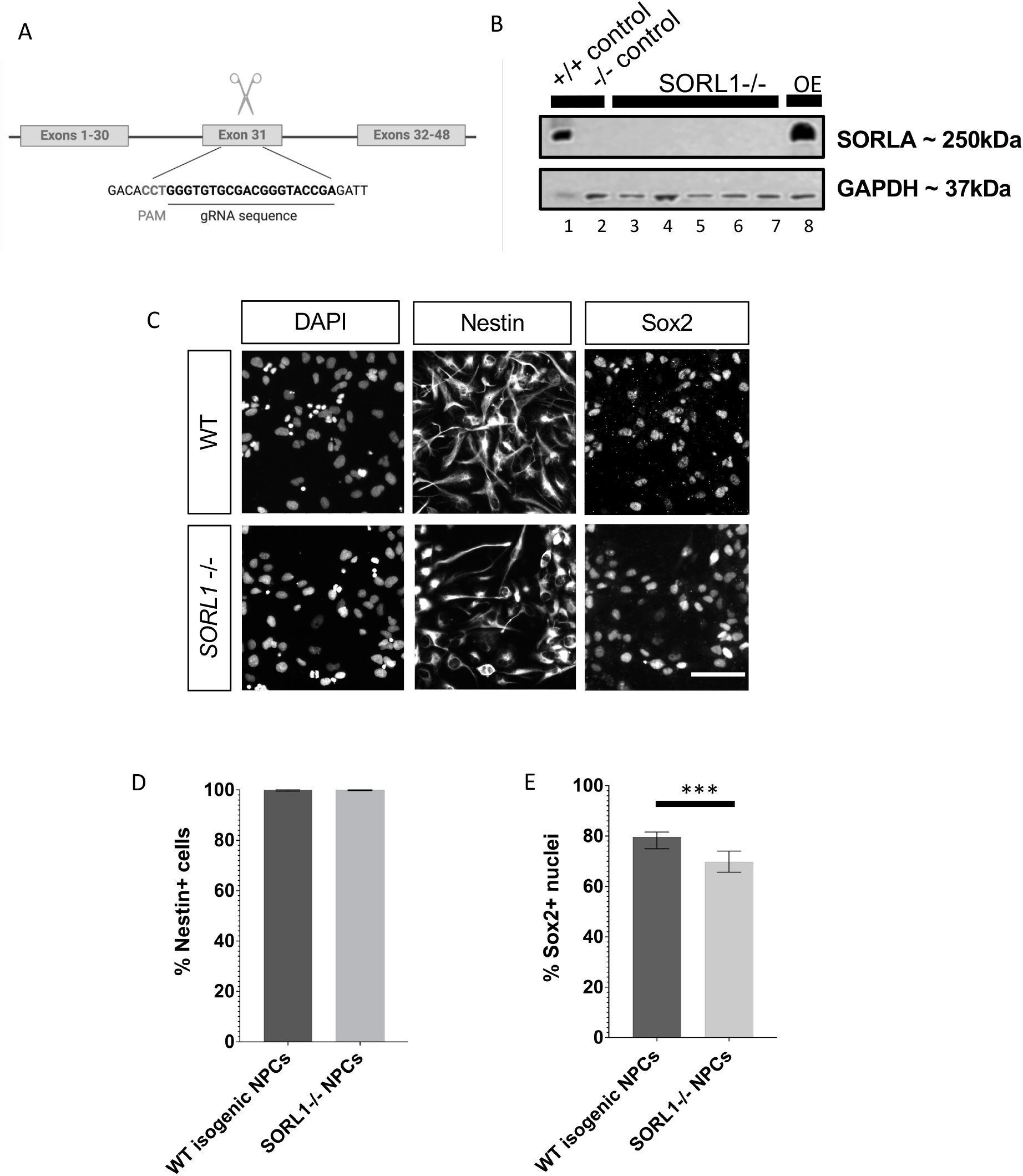
Depletion of SORLA by targeting of exon 31 in SORL1. (A) A single-guide RNA was used to target CRISPR-Cas9 cleavage at a site within exon 31 of the SORL1 gene to induce a homozygous mutation by non-homologous end-joining. (B) Mutations introduced using this method resulted in depletion of SORLA expression beyond levels detectable by immunoblotting by SORLA primary antibody with GAPDH loading control (lane 1 = unedited wild-type positive control, lane 2 = negative control, lanes 3-7 = multiple SORLA depleted subclones with lanes 4-6 representing SORL1-/- subclones utilized in the current study and lane 8 = SORLA overexpression in HEK cells as a positive control). (C) Representative grey-scale images of NPCs from wild-type and SORL1 -/- with immunocytochemistry used to detect Sox2 (stem cell marker) and Nestin (neural progenitor cell marker). (D) Quantification showed no change in the relative proportion of Nestin+ showed when comparing wild-type with SORL1-/- NPCs (Mann-Witney U test). (E) A significant decrease in the proportion of Sox2+ cells was found in SORL1 -/- NPCs compared to wild-type (Mann-Witney U, p<0.0001). Scale bar represents 50μm. Non-parametric testing was applied due to unequal variance and non-Gaussian distribution of cell proportion quantification. Graphed data was grouped according to genotype. The bar representing WT isogenic NPCs shows quantification of the median cell marker proportions from a pool of 88 images acquired from 2 wild-type subclones from the CRISPR-Cas9 editing to mutate SORL1 exon 31 and 1 parent wild-type line. The bar representing SORL1-/- NPCs shows median cell marker proportions from a pool of 88 images acquired from the 3 SORL1-/- subclones from the CRISPR-Cas9 editing to mutate SORL1 exon 31. Error bars show range. ***p < 0.0001.

### 4.2 Cell Painting assay labels cellular and subcellular components in wild-type and *SORL1^-/-^* NPCs in a pilot drug screen

To screen for compounds that rescue morphological profiles in SORLA-depleted NPCs, *SORL1^-/-^* NPCs were cultured for 24 hours with one of 330 compounds selected from the TargetMol Annotated Compound Set (L2110, targetmol.com; Supplementary Information Table 1) at one of three concentrations (100nM, 300nM or 1000nM) alongside vehicle-treated, wild-type and *SORL1^-/-^* NPCs. The Cell Painting assay was used to label cellular and subcellular compartments (Figure 2, Table 1). After pre-processing to remove redundant variables, 756 quantitative measurements of cellular features were selected for further analysis (Supplementary Information Table 2).

**Figure 2.**
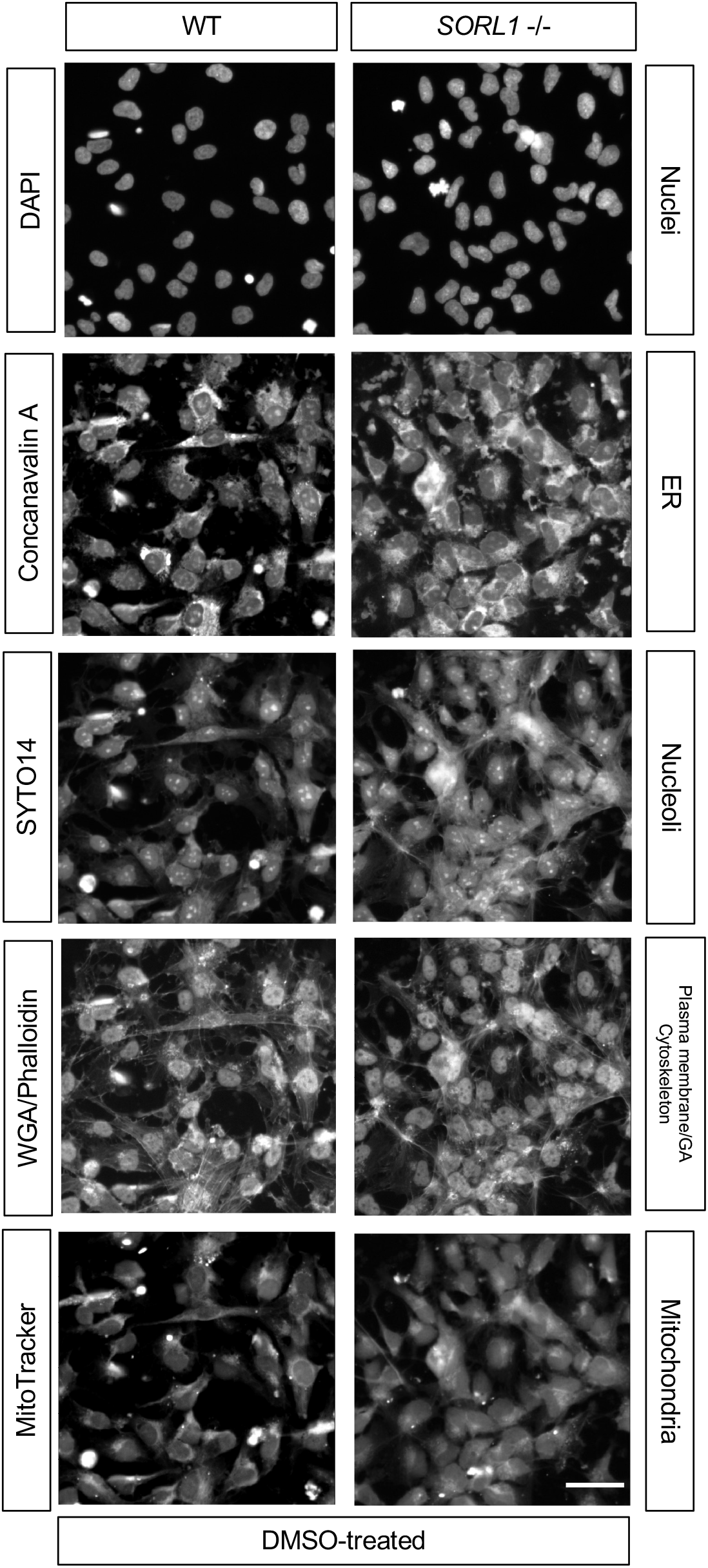
Cell Painting using conjugated cell compartment-specific dyes in wild-type and SORL1-/- neural progenitor cells. Monolayer, adherent NPCs were fixed (4% PFA) and a multiplexed reaction mix containing fluorescent conjugated dyes specific to various cellular and subcellular compartments applied to cells with subsequent image capture using the high-throughput, automated, confocal fluorescence microscopy platform ImageXpress Micro Confocal (20x). These grey-scale images demonstrate fluorescent labelling of nuclei by DAPI, endoplasmic reticulum by Concanavalin A, nucleoli/cytoplasmic RNA by SYTO14, plasma membrane and Golgi apparatus by Wheat Germ Agglutinin, cytoskeleton by Phalloidin and mitochondria by MitoTracker in DMSO wild-type and SORL1-/- NPCs and as such represent vehicle-treated positive and negative controls in the drug screening assay. Scale bar represents 50μm. Abbreviations: WT, wild-type; DMSO, dimethyl sulfoxide; RNA, ribonucleic acid;, WGA, Wheat Germ Agglutinin); NPC, neural progenitor cell; DAPI, 4’,6-diamidino-2-phenylindole.

### 4.3 *SORL1^-/-^* NPCs have distinct morphological profiles from wild-type control NPCs

Principal component analysis (PCA) was used to visualise the 756 features and phenotypic separation for the vehicle (DMSO)-treated wild-type and *SORL1^-/-^* NPCs in three dimensions (Figure 3A). PCA was again applied to reduce dimensionality of the 756 features to 50 non-redundant PCs that represent full array of features measured using the Cell Painting assay. These 50 PCs were used to define morphological signatures for the vehicle-treated wild-type, and vehicle-treated and untreated *SORL1^-/-^* NPCs, as well as sample treatment wells. Hierarchical clustering of phenotypic signatures for untreated and vehicle-treated *SORL1^-/-^* NPCs showed similar morphology for these two groups, which was distinct from that observed in the vehicle-treated wild-type control (Figure 3B). Euclidean distance measures did not perform well in distinguishing between the vehicle-treated positive (wild-type) and negative (*SORL1^-/-^*) control classes, so a non-Euclidean Bray-Curtis Dissimilarity metric was applied to test for significance of phenotypic separation when comparing phenotypic profiles from these two groups (Figure 3C).

**Figure 3.**
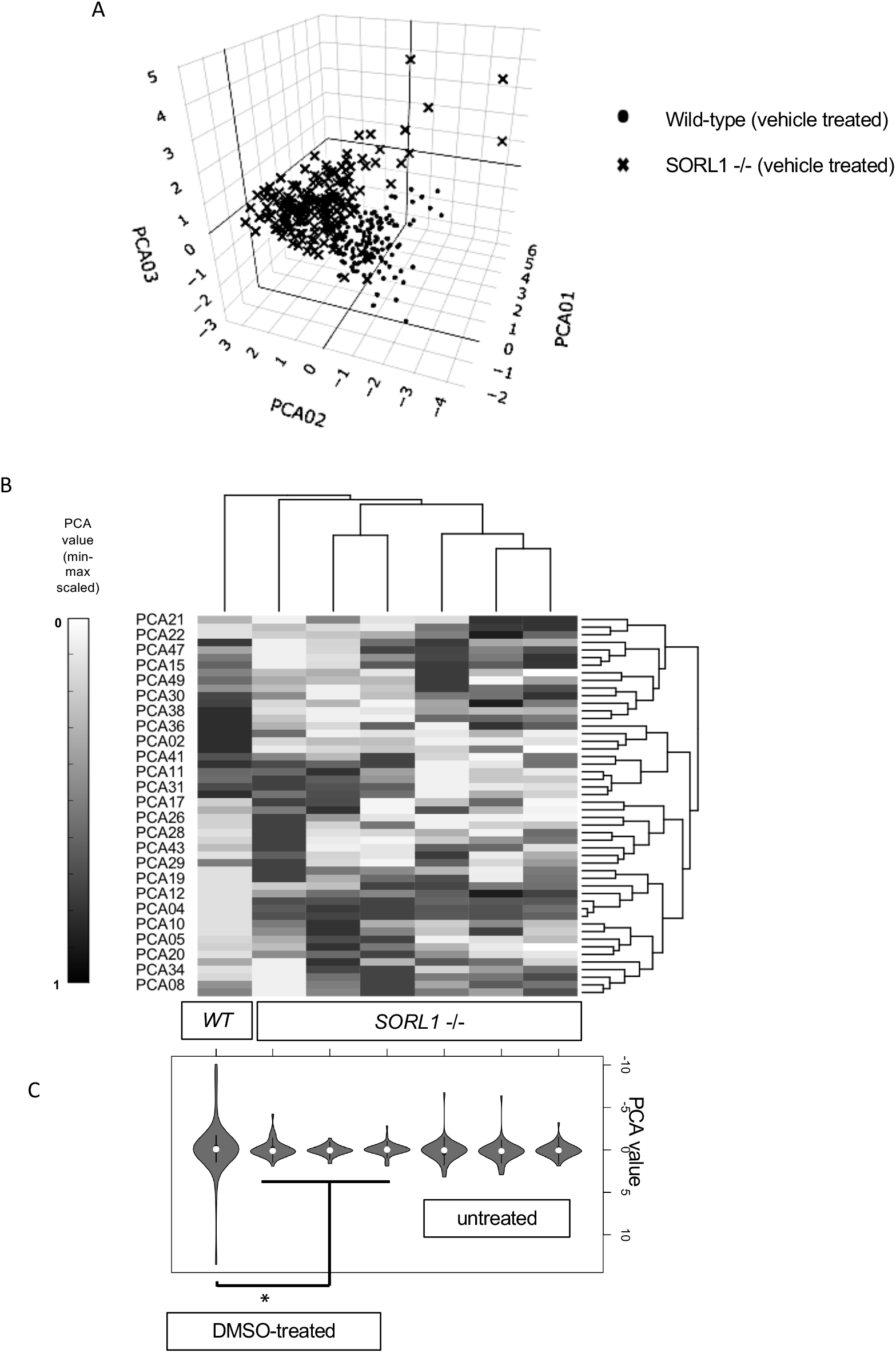
Morphology of SORL1-/- NPCs is distinct from isogenic wild-type controls. (A) PCA was used to visualize phenotypic separation of vehicle-treated SORL1-/- NPCs and isogenic wild-type controls in 3D. PCA was then used to reduce dimensionality from 756 quantitative measures of cellular features from the Cell Painting assay to 50 phenotypic PCs. (B) Hierarchical clustering and heatmap visualization of 50 PCs (after min-max scaling) showed separation of the untreated and vehicle-treated NPCs SORL1-/- NPCs, as well as separation of vehicle-treated SORL1-/- NPCs from vehicle treated wild-type NPCs. (C) Violin plots depict phenotypic signature of untreated and vehicle-treated NPCs SORL1-/- NPCs and vehicle-treated wild-type NPCs based on unscaled PC values for PCs 1-50 (not all PC labels displayed on axis), and significant separation was observed in the screening control classes vehicle-treated, i.e. morphological signature of DMSO-treated SORL1-/- NPCs was significantly different to that of the DMSO-treated wild-type NPCs. Abbreviations: 3D, three-dimensions; DMSO, dimethyl sulfoxide; NPC, neural progenitor cell; PC, principal component; PCA, principal component analysis); WT, wild-type.

### 4.4 Classification by machine learning predicts 14 drugs that reverse *SORL1^-/-^* NPC phenotypic profiles to that of wild-type NPCs

Our aim was to assay for drug induced phenotypic reversion using the quantitative phenotypic signatures described above. Neural network (NN) classification with 2-fold cross-validation was used to identify compounds that induced a morphological profile similar to that of the wild-type controls in the *SORL1^-/-^* NPCs. NN classification performance/accuracy was assessed using a confusion matrix with measures of sensitivity, specificity and detection rate (Figure 4A, Supplementary Information Table 3). Images from the positive control class were >99.68% likely to classify as a wild-type NPCs, while those from the negative control class were >99.64% likely to classify as *SORL1^-/-^* NPCs (Supplementary Information Table 4).

**Figure 4.**
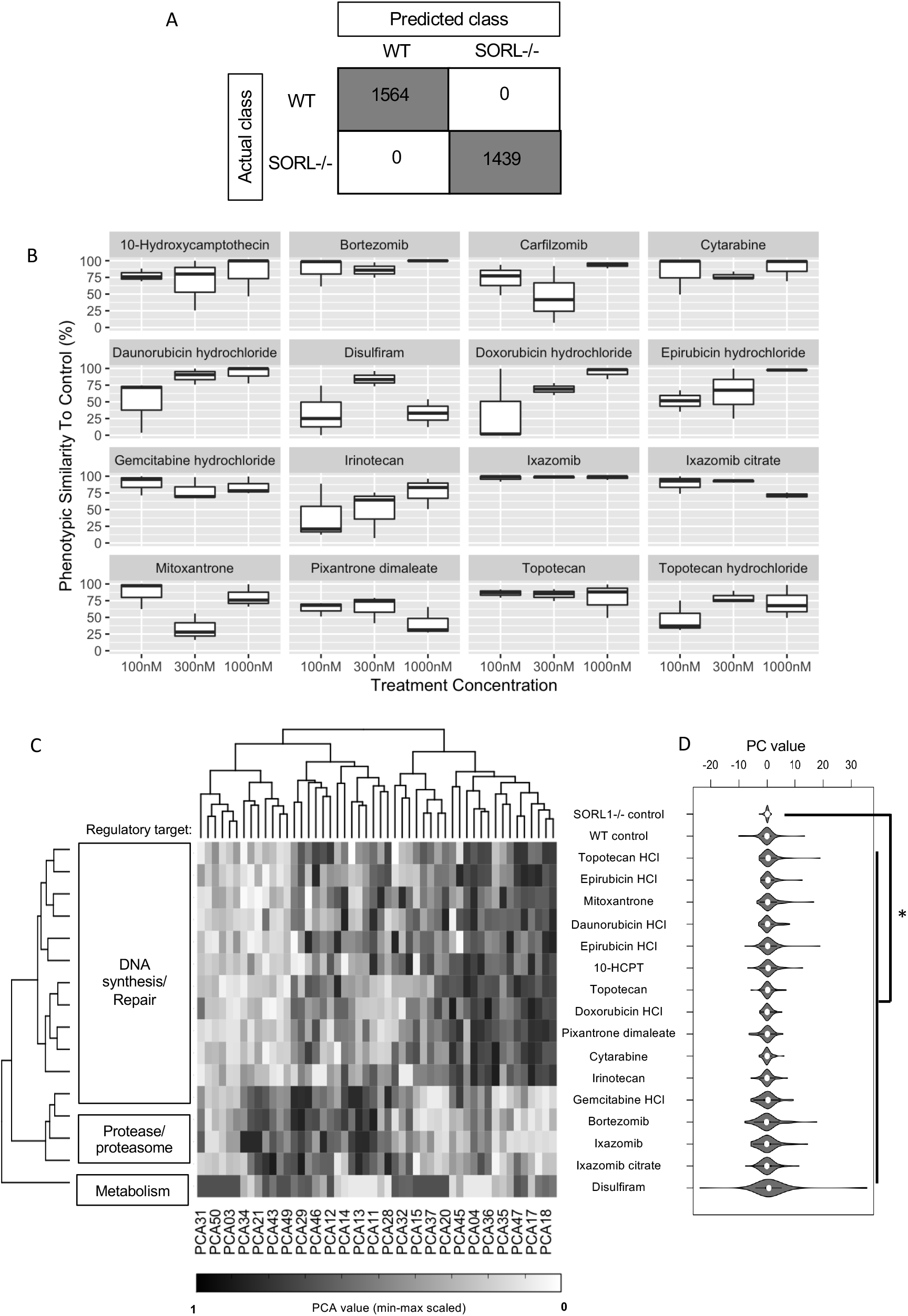
Neural network classification predicts 16 hit compounds that induce significant changes to SORL1-/- NPC morphology which increases similarity to isogenic wild-type controls. Sixteen inhibitor compounds regulating three biological pathways/targets from five mechanistic classes from the 330-compound library were identified from the pilot drug screen. (A) Confusion matrix demonstrating the accuracy of classification by Neural Network for actual and predicted WT and SORL1-/- control images. (B) Percentage likelihood of classification as a WT was used as a measure of phenotypic similarity to control (%, x-axis) and was measured for each compound treatment (330 compounds tested at 3 concentrations; 100nm, 300nm and 1000nM; y-axis) with compound hits selected (all 16 displayed) as those with >50.5% phenotypic similarity to control after 24h treatment. (C) Hierarchical clustering and heatmap visualization of 50 PCs (after min-max scaling, not all PC labels displayed on axis) showed clustering of morphological profiles according to annotated target class where similar morphology was induced by compounds targeted to DNA synthesis/repair pathways, and also similar morphological signatures were associated with treatment with compounds targeted to protease/proteasome and metabolism. (D) Violin plots depict phenotypic signature of treatment of SORL1-/- NPCs with the 16 hit compounds with reference to vehicle-treated SORL1-/- and wild-type NPCs based on unscaled PC values for PCs 1-50, and significant separation (Bray-Curtis dissimilarity test, p<0.05) was observed between DMSO vehicle-treated control SORL1-/- NPCs and the morphological signature induce by compound treatment for the 16 hits which showed >50.5% similarity to the wild-type control. No significant difference was observed between compound-treated SORL1-/- NPCs and the DMSO vehicle-treated wild-type controls. Abbreviations: 10-HCPT, 10-hydroxycamptothecin; NPC, neural progenitor cell; PC, principal component; PCA, principal component analysis); WT, wild-type.

Binary classification of compound-treated wells with reference to these control classes was performed.The NN classification model was used to output a similarity score between 0.0 and 1.0 for every compound-treated sample for each of the three *SORL1^-/-^* NPC subclones at each concentration (100nM, 300nM and 1000nM), where a score of 1.0 denotes 100% phenotypic similarity to the vehicle-treated wild-type controls with 0% similarity to the vehicle-treated *SORL1^-/-^* NPCs (negative controls) (Supplementary Figure 2)‥ Based on this, hits were defined as those compound-treated samples with a score of >0.505 after classification, which denotes a >50.5% similarity to the wild-type controls. From the hit selection by NN classification, 16 compounds reversed or partially reversed the phenotype of *SORL1^-/-^* NPC in all three subclones at one or more of the concentrations tested, i.e. there were 16 compounds where *SORL1^-/-^* NPC samples showed >50.5%similarity to the wild-type after treatment (Figure 4B).

These 16 compounds corresponded to 14 unique drug treatments since two of the 16 hits (topotecan and ixazomib) were represented with two formulations (ixazomib/ixazomib citrate and topotecan/topotecan HCl). According to library annotation, those 16 compounds (representing 14 drugs) were grouped into three classes: metabolism, protease/proteasome and DNA synthesis/repair inhibitors (Supplementary Information Table 1, Figure 4C). Phenotypic signatures induced by compounds from the same regulatory target annotation were hierarchically clustered, suggesting an overlapping morphological profile for compounds targeted to each regulatory process (Figure 4D), and phenotypic profiles similar to the wild-type controls were observed in *SORL1^-/-^* NPCs upon treatment with hit compounds (Figure 4E). All 16 compounds which reversed the mutant phenotype identified using NN classification induced morphological profiles in *SORL1^-/-^* NPC that were significantly different to negative control class (*p < 0.05*, Bray-Curtis dissimilarity test, Figure 4E).

PCs with the greatest predictive power in each of the classification models were ranked by relative importance; PC12, PC07 and PC02 were top-ranked for predictive power in the NN classification model (Supplementary Information Table 5). These components represented measures of plasma membrane, Golgi apparatus, mitochondrial, endoplasmic reticulum and cytoskeletal texture and radial distribution and nuclear eccentricity, shape, area, radial distribution and compactness.

### 4.5 Exploring mechanisms associated with SORLA depletion in NPCs

To understand the possible biological and molecular mechanisms that underpin the phenotypic changes associated with *SORL1^-/-^* NPCs, a first step was to use the STRING database to generate a protein-protein interaction network for physical interactors with SORLA, as this network will be disrupted by SORLA depletion. In order to annotate the network to indicate genes expressed in wild-type NPC lines, we generated a transcriptomic profile using RNA-sequencing data of wild-type NPCs (QOLG-1 iPSC donor line (Figure 5A). Having narrowed down the genes in the network to those expressed in NPCs we performed enrichment analysis. Significant enrichment of the NPC-expressed SORLA interactors (Figure 5A) was observed in multiple AD- and SORLA-related Gene Ontology Biological Processes (GOBP) terms (Figure 5B), e.g. vesicle-mediated transport to the plasma membrane (GOBP:0016192, FDR < 0.05), retrograde transport endosome to Golgi (GOBP:0042147, FDR < 0.05), regulation of Aβ formation (GOBP:1902003, FDR < 0.05) and neurofibrillary tangle assembly (GOBP:1902996, FDR < 0.05). Additionally, enrichment was observed in cellular components linked to the image-based phenotypic features relevant to SORLA function (e.g. the most significant enrichment was in early endosome GOCC:0005769 with FDR *p* = 7.29 × 10^-15^) and also those cellular components included in PC02, PC07 and PC12, which ranked as most important for distinguishing *SORL1^-/--^* NPCs from wild-type controls (Supplementary Information Table 5).

**Figure 5.**
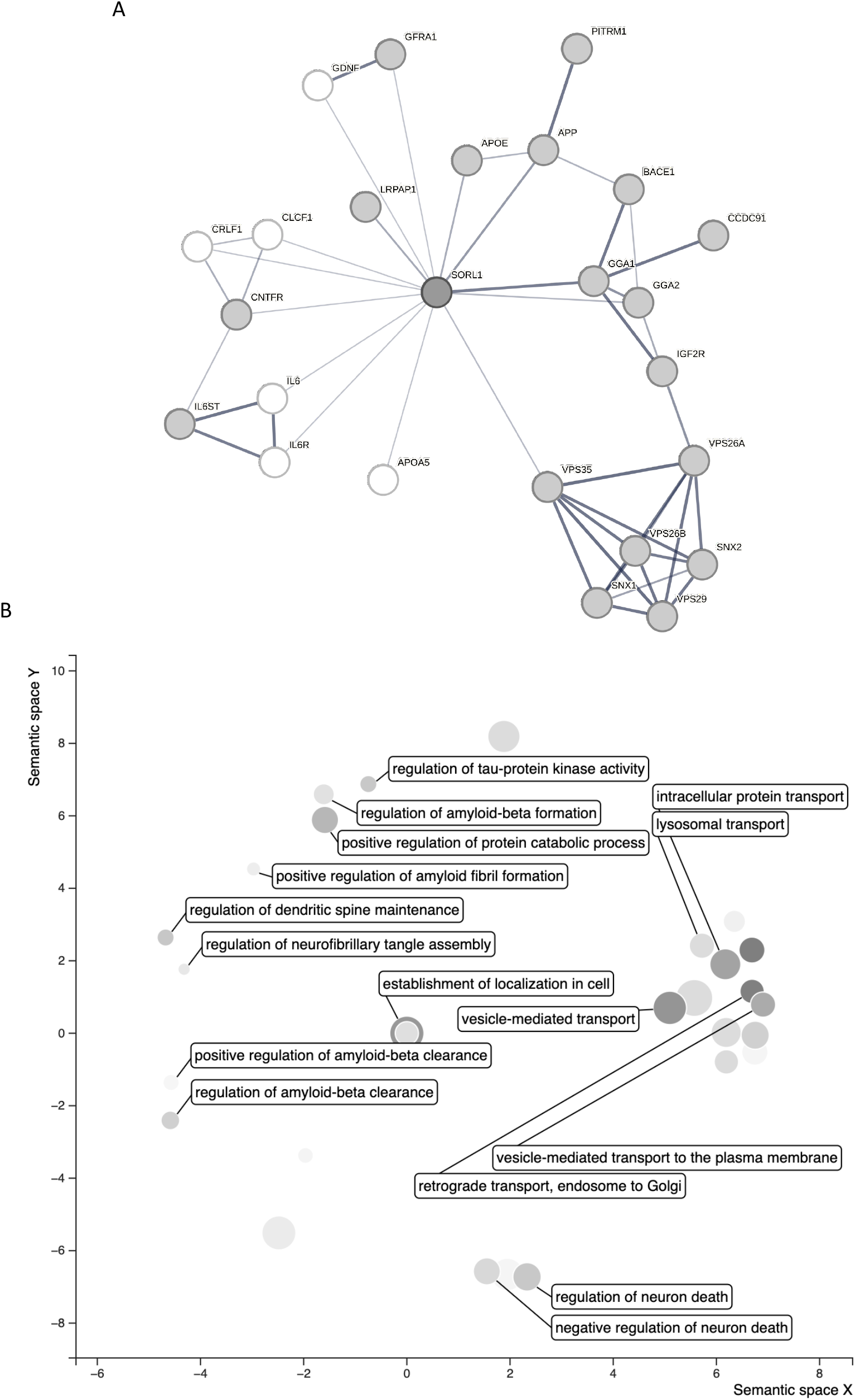
The molecular mechanisms associated with SORLA-depletion are enriched in Alzheimer’s disease relevant biological processes. (A) Multiple proteins known to interact with SORLA based on experimental evidence from the STRING database were expressed at transcriptomic-level in our NPCs (expressed genes shown in dark grey). (B) Enrichment of the network of NPC-expressed SORLA interactors (dark grey nodes from the network diagram in A) revealed significant overlap between genes interacting with SORLA, and those in GOBP terms associated with endolysosomal sorting and amyloid processing (visualization by semantic similarity produced using ReViGO).

### 4.6 Experimentally validated compound-protein interaction networks are enriched in pathways associated with DNA repair, metabolism and protease regulation

To understand the molecular targets and pathways associated with the phenotypic response of *SORL1^-/-^* NPCs to the hit compounds, we performed a STITCH-STRING database search for experimentally validated compound-target interactions for the 14 distinct drug molecule hits grouped by regulatory class. This confirmed that bortezomib, carfilzomib and Ixazomib target proteasome-related genes (Figure 6A), disulfiram targets aldehyde dehydrogenase 2 (Figure 6B) and the remaining compounds targeted to topoisomerase I (Figure 6C), topoisomerase II (Figure 6D) interact with TOP1 and TOP2A/2B respectively. Compounds inhibiting DNA synthesis (gemcitabine HCl and cytarabine) were targeted to multiple target genes expressed in our NPCs (Figure 6E). There were common targets for the compounds within each mechanistic class (Supplementary Figure 3). This data agrees with the functional/mechanistic annotation for the list of predicted hits provided with the compound library (Table 2).

**Figure 6.**
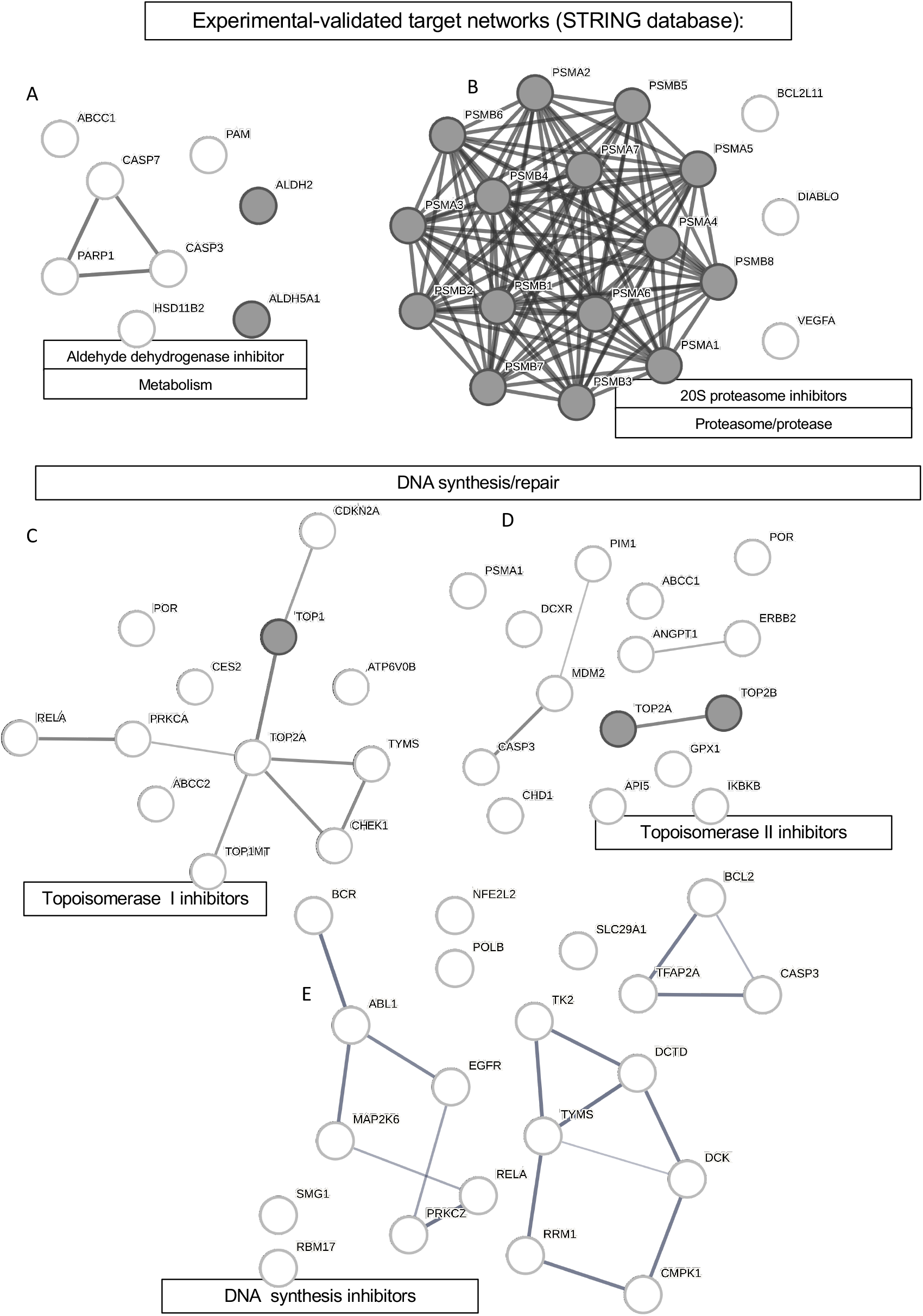
The 16 hit compounds target networks associated with annotated mechanism of action. The 16 compounds represent 14 FDA/internationally-approved drugs from 5 mechanistic classes targeed to 3 biological pathways. Network analysis using the STITCH-STRING database was used to explore and confirm annotated mechanism by interrogating compound-protein interactions for each mechanistic class. (A) The aldehyde dehydrogenase inhibitor disulfiram was has experimentally validated tarrgets ALDH2 and ALDH5A1. (B) Bortezomib and carfilzomib is targeted to multiple subinits of the 20S proteasome. (C) Topoisomerase I inhibitors topotecan, irinotecan and 10-hydroxycamptothecin were targeted to TOP1. (D) Topoisomerase II inhibitors were targeted to TOP2A and TOP2B. (E) DNA synthesis inhibitors gemcitabine HCl and cytarabine has no targets expressed in our NPCs. Nodes in grey indicate approved target annotation. All nodes were expressed in an independent line of wild-type human NPCs derived from the same parent line, QOLG-1 as determined by RNAseq analysis of transcriptome. STRING (protein only) networks shown in this figure, for STITCH-STRING network including compounds see Supplementary Figure 3).

**Table 2.** Table summarising the 14 drugs according to regulatory target and annotated molecular mechanism.

| FDA-approved annotation (target/class): |  |  |
| --- | --- | --- |
| Regulatory Target | Molecular mechanism | Compound Name |
| Metabolism | Aldehyde dehydrogenase inhibitor | Disulfiram |
| Proteasome/protease | 20S proteasome inhibitor | Bortezomib |
|  |  | Ixazomib/Ixazomib citrate |
|  |  | Carfilzomib |
| DNA synthesis/repair | Topoisomerase I inhibitor | Topotecan/topotecan HCl |
|  |  | Irinotecan |
|  |  | 10-hydroxycamptothecin |
|  | Topoisomerase II inhibitor | Pixantrone dimaleate |
|  |  | Mitoxantrone |
|  |  | Epirubicin HCl |
|  |  | Daunorubicin HCl |
|  |  | Doxorubicin HCl |
|  | DNA synthesis inhibitor | Gemcitabine HCl |
|  |  | Cytarabine |

Enrichment analysis was performed on the protein-protein interaction network associated with each group of compounds within the STRING database. The top eight targets expressed in NPCs for the aldehyde dehydrogenase inhibitor disulfiram included two genes from the family encoding aldehyde dehydrogenases (ALDH2, ALDH5A1). The most significant GO biological process and molecular function terms for the disufiram network were response to corticosteroid (GO:0031960, FDR *p* = 0.046) and cysteine-type endopeptidase activity involved in the execution phase of apoptosis (GO:0097200, PDR *p* = 0.0047) respectively (Figure 6A).

As expected, the proteasome inhibitors bortezomib, carfilzomib and Ixazomib target a network of 18 proteins expressed in our NPCs including 15 genes encoding subunits of the 20S proteasome (PSMA1, PSMA2, PSMA3, PSMA4, PSMA5, PSMA6, PSMA7, PSMB1, PSMB2, PSMB3, PSMB4, PSMB5, PSMB6, PSMB7 and PSMB8). The most significant GO biological process and molecular function terms for the bortezomib, carfilzomib and ixazomib network were proteasomal ubiquitin-independent protein catabolic process (GO:0010499, FDR *p* = 5.58 × 10^-36^) and threonine-type endopeptidase activity (GO:0004298, PDR *p* = 8.06 × 10^-37^) respectively (Figure 6B).

There were mechanistic classes included in the compounds regulating DNA synthesis/repair pathways. The topoisomerase I inhibitors topotecan, irinotecan and 10-hydroxycamptothecin target a network of 12 proteins expressed in our NPCs including topoisomerase I (TOP1). The most significant GO biological process and molecular function terms for this network were DNA topological change (GO:0006265, FDR *p* = 0.005) and DNA topoisomerase activity (GO:0003916, FDR *p* = 6.82 × 10^-5^) respectively (Figure 6C). The topoisomerase II inhibitors epirubicin, doxorubicin, daunorubicin, pixantrone and mitoxantrone target a network of 15 proteins expressed in our NPCs including topoisomerase II A and B (TOP2A and TOP2B). The most significant GO biological process and molecular function terms for this network were negative regulation of apoptotic process (GO:0043066, FDR *p* = 0.0012) and DNA topoisomerase type II (GO:0003918, FDR *p* = 0.0124) respectively (Figure 6D). Cytarabine and gemcitabine were associated with network of 20 proteins including multiple proteins associated with DNA biosynthetic processes (TK2, DCTD, TYMS and DCK) expressed in our NPCs (Figure 6E). However, the annotated mechanism of action for these compounds is not via protein-compound interaction but rather intercalation/incorporation into DNA during replication in S-phase of the cell cycle. In keeping with this, the network associated with cytarabine and gemcitabine was most enriched in deoxyribonucleoside monophosphate biosynthetic process (GO:0009157, FDR *p* =3.76 × 10^-6^) and transferase activity (transferring phosphorus-containing groups, GO:0016772, FDR *p* = 1.34 × 10^-5^).

### 4.7 Structural similarity and computational ligand-based prediction of novel biological targets and mechanistic pathways

Since compounds may have unexpected off-target effects not previously reported in drug library annotations or databases, we also explored mechanism of action for each of the 14 drugs using established cheminformatics methods such as structural similarity ligand-based target prediction. We used the Similarity Ensemble Approach (SEA) search tool to query compound-target databases with the smiles notation of the chemical structure of our 14 drug hits. We then used network and enrichment analysis to expand the compound target search space to derive a set of novel computationally predicted (rather than experimentally validated) compound-protein interactions for each drug.

With a total of 207, carfilzomib had the greatest number of predicted targets, followed by 66 predicted for bortezomib and between 1-33 predicted for the remaining compounds (Z-score <1.96; TC ≥0.4; *p* ≤ 10’^15^). To refine the predicted networks to include only those protein-protein interactions likely to occur in our cellular system, the target lists from the SEA search were filtered to include only targets expressed in our wild-type NPC line. Predicted protein-protein interaction networks were visualised for carfilzomib, bortezomib, disulfiram, mitoxantrone, irinotecan, cytarabine and daunorubicin/doxorubicin/epirubicin (which shared an identical predicted interaction network). As there were too few interactions predicted for genes expressed in NPCs for ixazomib, gemcitabine, topotecan and 10-hydroxycamptothecin, network analysis could not be performed.

Enrichment analysis was performed to determine which of the pathways had previously been associated with AD. Target networks for disulfiram, mitoxantrone, irinotecan, cytarabine and daunorubicin/doxorubicin/epirubicin were not enriched in KEGG pathway for Alzheimer’s disease. But target networks for the proteasome inhibitors carfilzomib (Figure 7A) and bortezomib (Figure 7B) were strongly associated with the KEGG pathway for Alzheimer’s disease (hsa05010; carfilzomib: FDR *p* = 7.56 × 10^-38^ and bortezomib: FDR *p* = 1.04 × 10^-43^). Nodes within the KEGG pathway annotation associated with AD are indicated in dark grey (Figure 7A and B). These included *BACE1* and *CAPN1/2* within the networks for both carfilzomib and bortezomib, as well as *PSEN1/2* in the network for bortezomib only.

**Figure 7.**
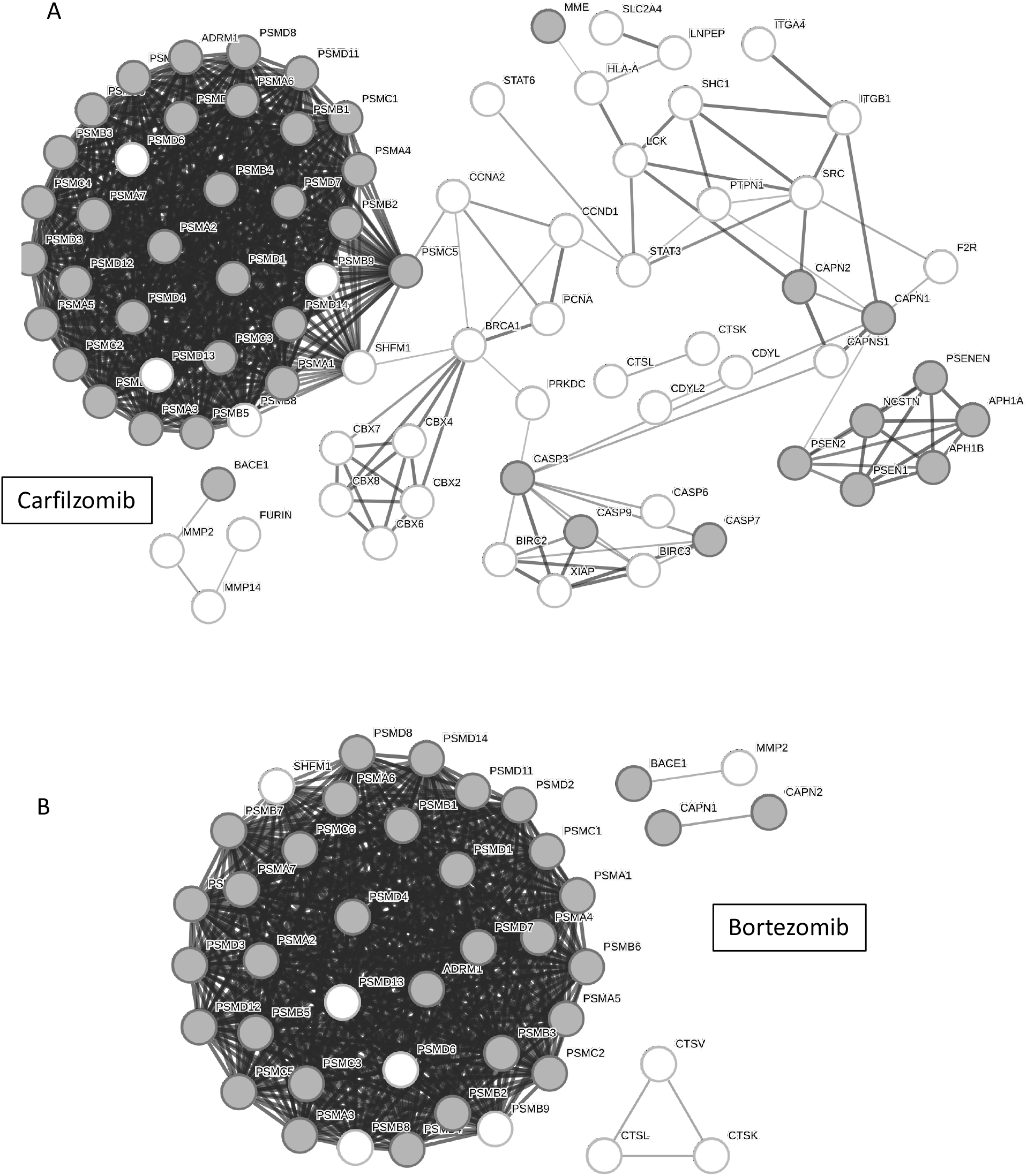
Predicted compound-protein interaction for proteasome inhibitors are enriched in Alzheimer’s disease-associated genes. The proteasome inhibitors (A) carfilzomib and (B) bortezomib were predicted to target multiple subunits of the 20S proteasome based on structural similarity. Enrichment analysis showed multiple nodes in the target network for each compound was enriched in the KEGG pathway for Alzheimer’s disease (hsa05010). Nodes in grey depict those molecular targets which are both part of the predicted compound target and feature in the gene list for KEGG pathway hsa05010.

## 5 Discussion

Drug development for dementia has had an extremely high rate of attrition in clinical trials over many decades (Cummings, 2018; Devenish, 2020). While recent progress, with the immunotherapy drugs appears promising (Dhillon, 2021; Ferrero et al., 2016; Lalli et al., 2021), efficacy and minimising off-target and adverse reactions requires further development (Herring et al., 2021; Song et al., 2022) and here is a need for alternative therapeutic approaches. One possible reason for the previous high failure rate of AD drugs is their development in animal models which do not faithfully recapitulate human disease. One possible reason for the previous high failure rate of AD drugs is their development in animal models. Here, we used human iPSC-derived NPCs, a strategy that may increase the likelihood of success, as candidate compounds are tested on human cells during discovery and development, facilitating early efficacy testing in the context of human physiology, and allowing early recognition of human-specific toxicity. We used loss of SORLA as a cellular screening model for drug discovery because of the strong genetic and functional evidence supporting *SORL1*/SORLA’s involvement in AD. The fact that previous studies had identified image-based endosomal phenotypes in *SORL1^-/-^* NPCs provided the rationale for using Cell Painting, a technique which allows for relatively unbiased interrogation of cell morphology and has been used previously used in cancer drug screening studies (for example, Gustafsdottir et al., 2013; Hughes et al., 2020; Warchal et al., 2020). Cell Painting is a type of high-content image analysis that generates quantitative morphological profiles. Here we demonstrate that it can be applied to distinguish wild-type hiPSC-derived NPCs from those lacking SORLA. The principal component with the greatest predictive power in the classification models represented measures pertaining to the ER, Golgi apparatus and plasma membrane, which is consistent with SORLA’s role in intracellular trafficking. We further showed that the Cell Painting assay permitted classification of hits from a pilot screen of 330 FDA approved compounds and identified 16 compounds, representing 14 drug molecules, where there was evidence for reversion of the mutant phenotype at one or more of the concentrations tested.

Given SORLA interacts directly and indirectly with a large number of other proteins, a drug screening assay that is agnostic to the cellular impact of SORLA loss, and which utilises high dimensionality morphological datasets to capture a broad range of phenotypes may be more appropriate than an assay focused on a single phenotype. This approach also seems to be well suited to a disease such as AD, where pathology is likely to arise via dysregulation of multiple cellular functions. In keeping with this, the 16 hits from the drug screen induced changes in multiple cellular/subcellular compartments, including the nucleus, mitochondria, endoplasmic reticulum, Golgi apparatus and cell membrane, suggesting that the high-content approach identifies compounds with diverse target proteins/pathways and mechanism of action.

The hits included four inhibitors of proteasome/proteases, eight topoisomerase inhibitors, three DNA synthesis inhibitors and an aldehyde dehydrogenase inhibitor, representing three biological pathways: DNA damage/repair, metabolism and protein degradation. Recent genetic studies have linked mutations in aldehyde dehydrogenase with cognitive impairment (Jin et al., 2022). Aldehyde dehydrogenase 2 activity is important for regulating neuroinflammation and Aβ-levels *in vivo*, and variants in the *ALDH2* gene have been associated with AD (Joshi et al., 2019). Modification of this pathway, using chemical compounds related to disulfiram may, therefore, be useful in the regulation of this aspect of AD neuropathology.

While SORLA has not previously been associated with DNA damage/repair, deletion of another VPS10-domain family member, *SORCS2*, led to increased DNA double strand breaks (DSBs) in the dentate gyrus in mice and to higher numbers of topoisomerase 2β-dependent DSBs in human dopaminergic neurons (Gospodinova et al., 2021). In addition, there is mounting evidence for a relationship between DNA damage and neurodegeneration (Pessina et al., 2021; Ross and Truant, 2017). Some drugs which target DNA damage/repair pathway are central nervous system penetrant, for example topotecan rapidly passes through the blood-brain barrier (Friedman et al., 1994; Rapisarda et al., 2004) making them appropriate for therapies targeted to the brain.

As inhibition or impairment of the proteasome has been associated with AD status (Thibaudeau et al., 2018; Upadhya and Hegde, 2007), it was unexpected that drugs inhibiting the proteasome rescued *SORL1^-/-^* associated morphology. The phenotypic rescue we observed during the drug screen may be indicative of a difference in the impact of proteasome inhibition in NPCs and neurons, or it may point to a previously uncharacterised mechanism of action of the appropriate drug(s). This analysis revealed a number of non-proteasome linked, AD-associated predicted targets of our drug hits. For example, production of Aβ depends on proteolysis of APP by β- and g-secretases (Nunan and Small, 2000; Zhang et al., 2011). Interestingly, β-secretase (which is expressed in NPCs) is a computationally predicted target of both bortezomib and carfilzomib, which are both annotated as proteasome inhibitors. Incorporation of bortezomib into amyloidosis treatment regimens has been reported to significantly improve patient outcomes in clinical trials (Huang et al., 2016; Kastritis et al., 2019). Such instances illustrate the point that pharmacological confirmation of ligand-target interaction is beneficial to translating a possible lead compound towards drug candidate nomination.

In terms of limitations, while a library of compounds enriched for drugs targeted towards cancer is not necessarily an obvious choice for an AD study, the fact that NPCs are pluripotent, stem-like cells means that profiling the morphological response to compounds from a library typically active in undifferentiated cell types is arguably a prudent initial choice. This is particularly important given that morphological features associated with SORLA depletion are similar in both NPCs and neuronal cell types (Hung et al., 2021; Knupp et al., 2020).

As compounds that cause reversion of the mutant phenotype do not act by altering SORLA directly the precise mechanisms by which they elicit their response remains to be characterised. The results of our network analysis support future deconvolution of molecular targets, which is an essential next step, as translation of drug treatments to clinical testing requires a detailed understanding of mechanism. Additionally, compound safety requires a clear understanding of possible off-target effects. We conducted an *in silico* target-ligand interaction search using a database of experimentally validated interactions (STRING) and computationally-predicted interactions based on structural similarity (SEA). The *in silico* networks identified suggest possible targets lines could direct future target-ligand interaction and binding affinity assays, whilst insights from the pathway/enrichment analysis will support exploration of the downstream mechanisms responsible for the phenotypic reversion observed in this study.

AD pre-clinical research is largely focussed on mature neural cells, such as neurons and/or glia. This is logical given that these cells are relevant to the disease mechanisms as they are currently understood (Scheltens et al., 2021). But recent evidence suggests NPCs are also important in AD. Proliferative, pluripotent NPCs persist in the ageing brain (Tobin et al., 2019) and there is a greater decrease in adult hippocampal neurogenesis in AD cases than in healthy controls (Moreno-Jiménez et al., 2019). Our findings in *SORL1^-/-^* NPCs are in keeping with this: there were fewer nuclei positive for the multipotent neural stem cell marker Sox2 than in wild-type cultures. This suggests that SORLA loss in NPCs may lead to alterations in the levels of pluripotent NPCs (which may be relevant during development and/or adulthood).

In summary, our goal was to apply morphological profiling via Cell Painting to differentiate wild-type neural progenitor cells from those lacking SORLA and thus develop a primary drug screening assay to discover compounds for pre-clinical testing and future translation for AD. Using Cell Painting, we discovered a morphological signature that distinguishes neural progenitor cells lacking *SORL1* from wild-type isogenic controls and demonstrated that this assay has the potential for use in small-molecule and drug library screening. A set of putative hits was identified, but follow-up studies to confirm their effect and potency via dose-response profiling, as well as target/mechanism of action, is needed to determine their translational potential. Our findings indicate that cell-types such a NPCs and drug libraries purposed for other diseases represent a chance to identify compounds/classes of compounds that may not be identified using more traditional, hypothesis-based screening approaches. Future experiments that adopt this methodology could screen larger drug libraries with greater diversity in terms of mechanistic action, targets and pathways, as well as libraries of small molecules with unknown pathways/mechanisms for novel drug discovery opportunities in AD.

## Supporting information

Supplementary Figures and Supplementary Information inc. Tables

