## Supplementary Figures and Supplementary Information inc. Tables for "Morphological profiling by Cell Painting in human neural progenitor cells classifies hit compounds in a pilot drug screen for Alzheimer’s disease"

*Supplementary Information Table 2 | Table of the 756 non-redundant image-based quantitative features (and mean value) for morphological profiling in the drug screen.* .. 17

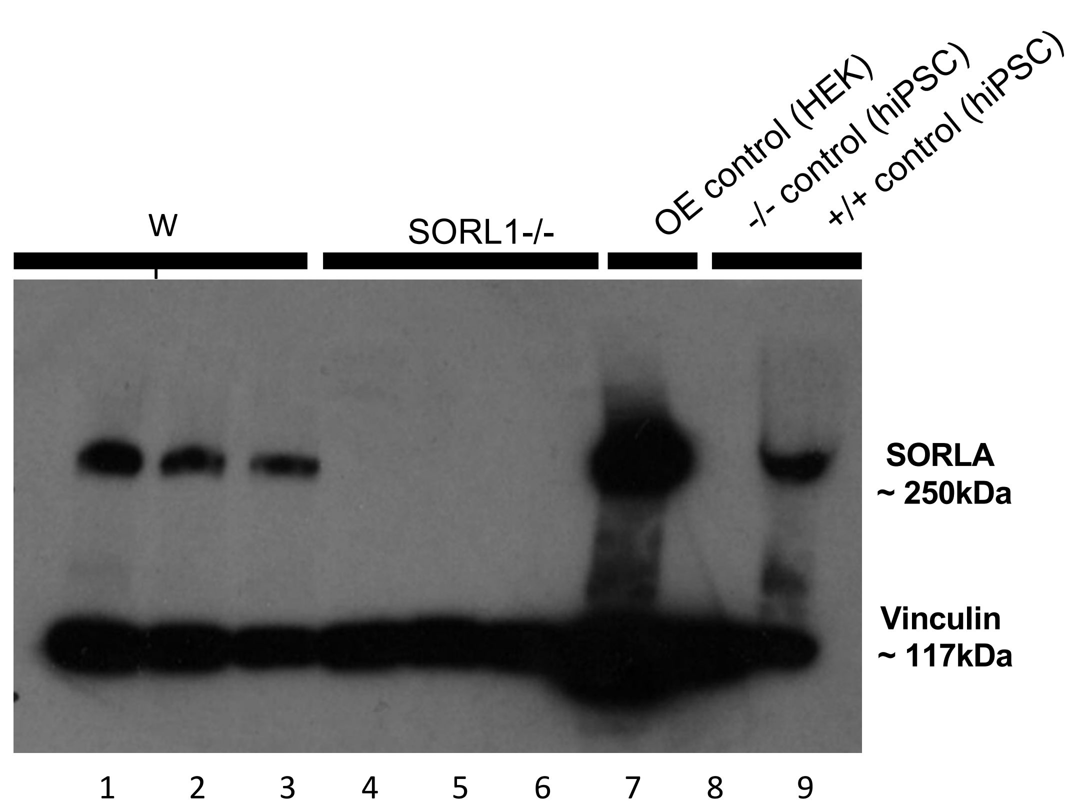

**Supplementary Figure 1 | Immunoblot for SORLA in neural progenitors derived from SORL1-/- and wild-type hiPSCs.**

Mutations were introduced in iPSCs by non-homologous end-joining using a CRISPR/Cas9 guide RNA targeted to exon 31 of the SORL1 gene. Neural induction of iPSCs to NPCs was performed using a neural induction kit with SMAD inhibition (embryoid body method, Stem Cell Technologies). This method resulted Sox+/Nestin+ NPCs in which depletion of SORLA expression beyond levels detectable by immunoblotting was observed. For this immunoblot, a SORLA primary antibody with Vinculin loading control (lane 1-3 = unedited wild-type control, lane 2 and 3 = wild-type CRISPR controls, lanes 4-6 = multiple SORLA depleted subclones, lane 7 = SORLA overexpression in HEK cells as a positive control, lane 8 = a negative control from SORL1-/- hiPSCs and lane 9 = wild-type hiPSCs as a positive control).

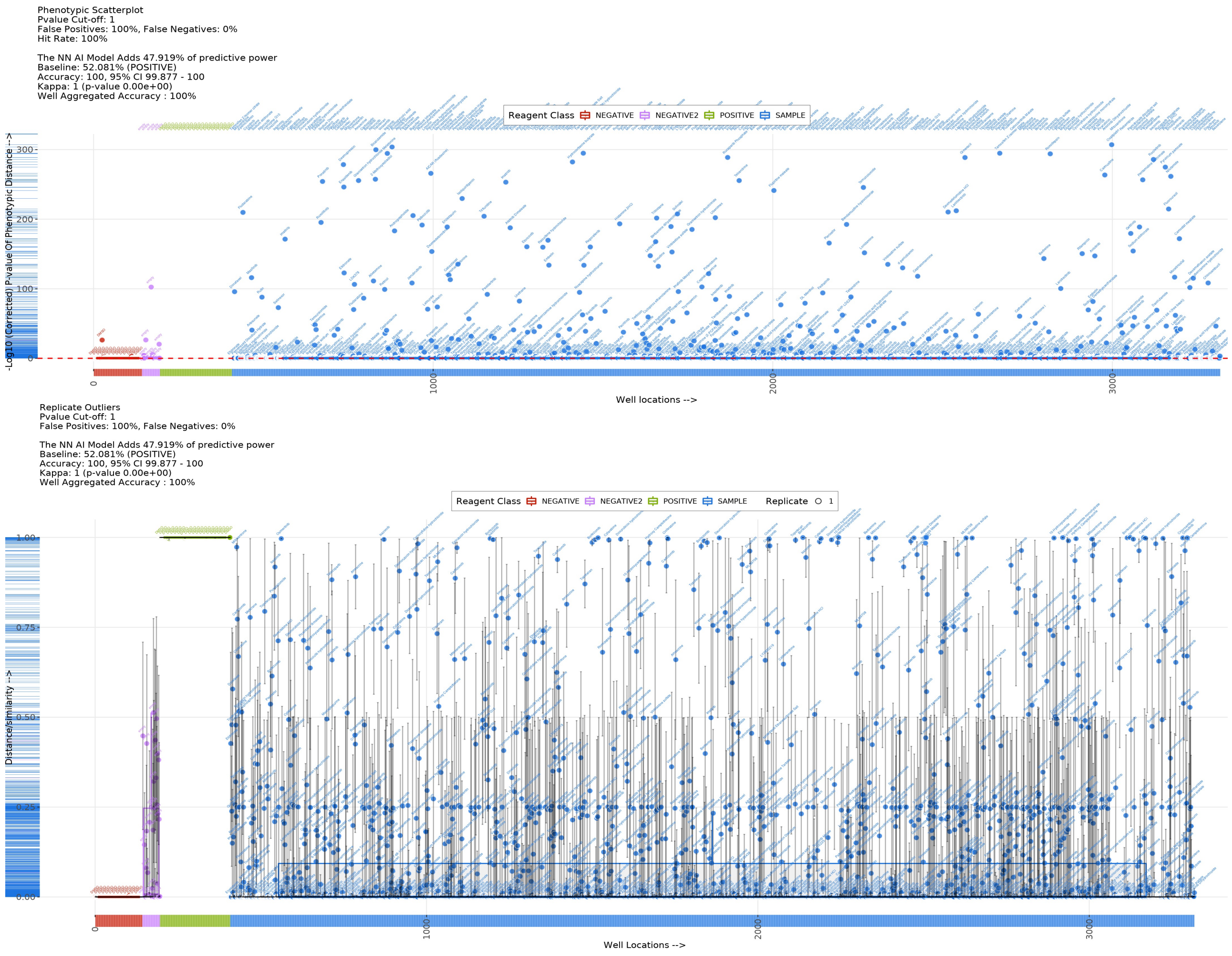

**Supplementary Figure 2 | Significance of the phenotypic distance for control and compound-treated SORL1-/- NPCs (top) and phenotypic similarity scores (bottom) with reference to wild-type controls based on neural network classification summarized by box plot.**

Box plots showing the significance of the phenotypic distance (-log10 corrected p-value), top panel, y-axis) for each treatment well (well location plotted on x-axis). The phenotypic similarity was determined from binary classification results from the neural network analysis (bottom panel, y-axis) such that a score 0.0 – 1.0 was calculated for each treatment well (well location plotted on x-axis). Phenotypic similarity >0.505 represents >50.5% likelihood of classification as a wild-type after treatment in SORL1-/- NPCs. Each data point represents the aggregated well-median (based on image-level median values from object-level measurements in CellProfiler). Red data represent DMSO vehicle treated SORL1-/- NPCs (NEGATIVE control class, reference class for phenotypic similarity/distance measures and training class for neural network algorithm). Purple data points represent untreated SORL1-/- NPCs (NEGATIVE2). For SORL1-/- NPCs, images were captured from generated from 3 CRISPR-derived subclones treated with 100nM, 300nM or 1000nM concentrations. Green data points represent DMSO vehicle-treated, wild-type NPCs derived from the parental hiPSC line (POSITIVE control class, training class for neural network classification). The supervised classes were the positive and negative control classes. Model output was 100% accurate, with no missclassification of positive and negative control images in test set (training 80%: test 20%). Compound hits were selected if they induced phenotypic similarity >50.5% in NPCs derived from all 3 SORL1-/- CRISPR-derived hiPSC subclonal lines.

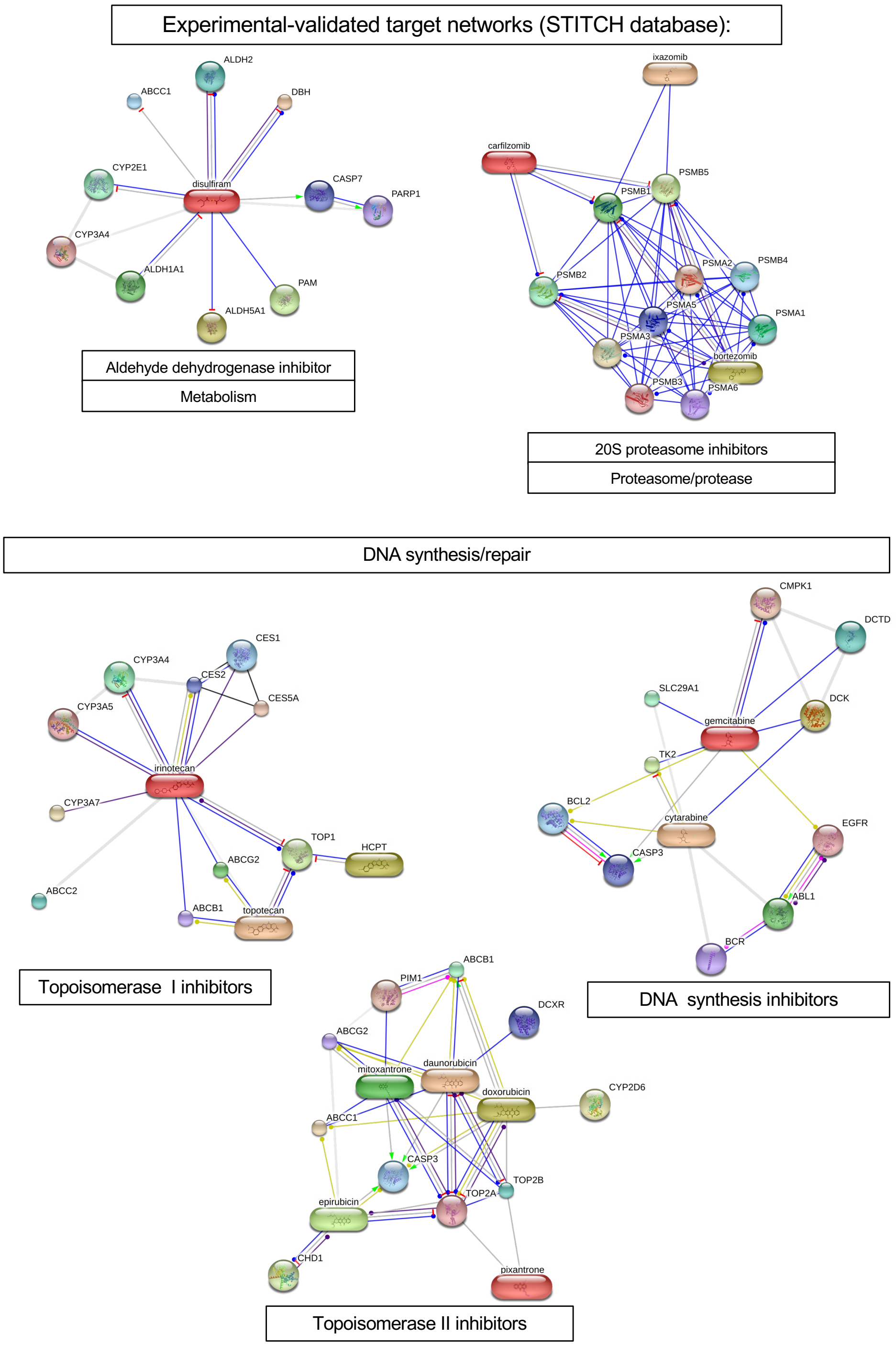

**Supplementary Figure 3 | Network visualization of compound hits grouped by mechanistic class using STITCH-STRING database confirms targeting to annotated proteins.**

Sixteen inhibitor compounds regulating three biological pathways/targets from five mechanistic classes from the 330-compound library were identified from the pilot drug screen. Those 16 compounds represent 14 FDA/internationally-approved drugs, were grouped by mechanistic class to define experimentally-validate target networks from the STITCH/STRING database. Compound-protein interactions for each mechanistic class observed with compound (oval nodes) and protein target (circular nodes**).**

**Supplementary Information Table 1 | Screening of 330 compounds from the TargetMol L2110 library of approved anti-cancer compounds (hits identified in screen are highlighted yellow).**

| Name | Pathways | Target |
|---|---|---|
| zanubrutinib | Angiogenesis | BTK |
| Pyrvinium pamoate | Cytoskeletal Signaling;Stem Cells | Wnt/beta-catenin |
| Sivelestat | Proteases/Proteasome | Serine Protease inhibitor |
| Olmutinib | Angiogenesis; JAK/STAT signaling; Tyrosine Kinase/Adaptors | EGFR inhibitor |
| Darolutamide | Endocrinology/Hormones | Androgen Receptor antagonist |
| Larotrectinib sulfate | Tyrosine Kinase/Adaptors | Trk receptor inhibitor |
| Ixabepilone | Cytoskeletal Signaling | Microtubule Associated inhibitor |
| Eltrombopag Olamine | Others | Thrombin agonist |
| Tazarotene | Metabolism | Retinoid Receptor inhibitor |
| Tamibarotene | Metabolism | Retinoid Receptor agonist |
| Raltitrexed | DNA Damage/DNA Repair | DNA/RNA Synthesis inhibitor |
| Pirarubicin | DNA Damage/DNA Repair | Topoisomerase inhibitor |
| Nelarabine | DNA Damage/DNA Repair | DNA/RNA Synthesis inhibitor |
| Mitoxantrone | DNA Damage/DNA Repair | Topoisomerase inhibitor |
| Histamine 2HCl | GPCR/G Protein; Immunology/Inflammation; Neuroscience | Histamine Receptor agonist |
| Flupirtine maleate | Membrane transporter/Ion channel; Neuroscience | NMDAR antagonist; Potassium Channel antagonist |
| Fludarabine Phosphate | DNA Damage/DNA Repair | DNA/RNA Synthesis inhibitor |
| Encorafenib | MAPK Signaling | Raf inhibitor |
| Embelin | Apoptosis; GPCR/G Protein; Metabolism | IAP inhibitor; Lipoxygenase inhibitor; Prostaglandin Receptor inhibitor |
| Dexmedetomidine HCl | GPCR/G Protein | Adrenergic Receptor agonist |
| Calcium Levofolinate | Others | Others |
| Bexarotene | Metabolism | Retinoid Receptor inhibitor |
| (S)-crizotinib | DNA Damage/DNA Repair | MTH1 inhibitor |
| Copanlisib | PI3K/Akt/mTOR signaling | PI3K inhibitor |

| | | |
|---|---|---|
| Calcitriol | Metabolism | Vitamin inhibitor |
| Lonafarnib | MAPK | Raf inhibitor; Ras inhibitor |
| Tosedostat | Metabolism | Aminopeptidase |
| Enzastaurin | Cytoskeletal Signaling | PKC inhibitor |
| Fosbretabulin Disodium | Cytoskeletal Signaling | Microtubule Associated inhibitor |
| Talazoparib | Chromatin/Epigenetic; DNA Damage/DNA Repair | PARP |
| Palbociclib Isethionate | Cell Cycle/Checkpoint | CDK |
| Palbociclib hydrochloride | Cell Cycle/Checkpoint | CDK |
| Lapatinib Ditosylate | Angiogenesis; MAPK; JAK/STAT signaling; Tyrosine Kinase/Adaptors | EGFR |
| Entinostat | Chromatin/Epigenetic; DNA Damage/DNA Repair | HDAC |
| Imatinib | Angiogenesis; Tyrosine Kinase/Adaptors; Cytoskeletal Signaling | Bcr-Abl; c-Kit; PDGFR |
| Irinotecan | DNA Damage/DNA Repair | Topoisomerase inhibitor |
| Pemetrexed Disodium Hydrate | DNA Damage/DNA Repair; Metabolism | DHFR inhibitor; DNA/RNA Synthesis inhibitor |
| Selumetinib | MAPK | MEK; ERK |
| Abiraterone | Metabolism | P450 inhibitor |
| Vinorelbine Tartrate | Cytoskeletal Signaling | Microtubule Associated inhibitor |
| Ledipasvir | Microbiology&Virology; Proteases/Proteasome | HCV Protease inhibitor |
| Ribociclib | Cell Cycle/Checkpoint | CDK inhibitor |
| Icotinib | Angiogenesis; JAK/STAT signaling; Tyrosine Kinase/Adaptors | EGFR inhibitor |
| Rucaparib Phosphate | Chromatin/Epigenetic; DNA Damage/DNA Repair | PARP |
| Selinexor | Membrane Transporter/Ion Channel | CRM1 inhibitor |
| Entospletinib | Angiogenesis | Syk |
| Gemcitabine HCl | DNA Damage/DNA Repair | DNA/RNA Synthesis |
| Telotristat Etiprate | Metabolism | Hydroxylase activator |
| Pacritinib | Angiogenesis; Tyrosine Kinase/Adaptors; Chromatin/Epigenetic; Tyrosine Kinase/Adaptors; JAK/STAT signaling; Stem Cells | FLT inhibitor; JAK inhibitor; Tyrosine Kinases inhibitor |
| Volasertib | Cell Cycle/Checkpoint | PLK |
| Zosuquidar 3HCl | Membrane transporter/Ion channel; Neuroscience | P-gp modulator |

| | | |
|---|---|---|
| Romidepsin | Chromatin/Epigenetic; DNA Damage/DNA Repair; NF-Kb | HDAC inhibitor |
| Enzalutamide | Endocrinology/Hormones | Androgen Receptor |
| Larotrectinib | Tyrosine Kinase/Adaptors | Trk receptor |
| 3-Bromopyruvic acid | Metabolic Enzyme/Protease | Hexokinase |
| Topotecan | DNA Damage/DNA Repair | topoisomerase |
| Cabozantinib hydrochloride (849217-68-1(free base)) | Angiogenesis; Tyrosine Kinase/Adaptors | c-Kit; c-Met/HGFR; TAM Receptor; VEGFR |
| Desmopressin | GPCR/G Protein | Vasopressin Receptor |
| Nimustine Hydrochloride | Others | Others |
| Nintedanib Ethanesulfonate Salt | Tyrosine Kinase/Adaptors | FGFR;PDGFR; VEGFR |
| Berberine | Microbiology&virology | Antibacterial |
| Neticonazole Hydrochloride | Microbiology&virology | Antifungal |
| Pipobroman | Others | Others |
| Estramustine phosphate sodium | Cytoskeletal Signaling | Microtubule Associated |
| Gilteritinib | Tyrosine Kinase/Adaptors | FLT;TAM Receptor |
| Miriplatin | DNA Damage/DNA Repair | DNA Alkylation |
| Goserelin acetate | GPCR/G Protein | GNRH Receptor inhibitor |
| Anlotinib Dihydrochloride | Tyrosine Kinase/Adaptors | VEGFR inhibitor |
| Pentostatin | Neuroscience | AChR inhibitor |
| Berbamine | Neuroscience | CaMKII inhibitor |
| Jaceosidin | Metabolism | UGT inhibitor |
| Erdafitinib | Angiogenesis; Tyrosine Kinase/Adaptors | FGFR inhibitor |
| Entrectinib | Angiogenesis; Immunology/Inflammation; Tyrosine Kinase/Adaptors | ALK inhibitor; ROS inhibitor; Trk receptor inhibitor |
| Osimertinib mesylate | Angiogenesis; JAK/STAT signaling; Tyrosine Kinase/Adaptors | EGFR |
| Relugolix | GPCR/G Protein | GNRH Receptor inhibitor |
| Acalabrutinib | Angiogenesis | BTK inhibitor |
| Cobimetinib | MAPK | MEK inhibitor |
| Brigatinib | Angiogenesis; Immunology/Inflammation; Tyrosine Kinase/Adaptors; JAK/STAT signaling | ALK inhibitor; EGFR inhibitor; FLT inhibitor; IGF-1R inhibitor; ROS inhibitor |
| Ivosidenib | Metabolism | Dehydrogenase inhibitor |
| Bestatin hydrochloride | Cytoskeletal Signaling; Metabolism | Aminopeptidase inhibitor; Integrin inhibitor |
| Vesnarinone | Metabolism | PDE |

| | | |
|---|---|---|
| Anlotinib | Angiogenesis; JAK/STAT signaling; Tyrosine Kinase/Adaptors | EGFR inhibitor |
| Homoharringtonine | JAK/STAT signaling; Stem Cells | STAT inhibitor |
| Cephalotaxine | Immunology/Inflammation | Antiviral inhibitor |
| Niraparib | Chromatin/Epigenetic; Others; DNA Damage/DNA Repair | Others inhibitor; PARP inhibitor |
| Troglitazone | Metabolism | PPAR agonist |
| Carbendazim | Microbiology&virology | Antifungal |
| LY2835219 | Cell Cycle/Checkpoint | CDK inhibitor |
| Carmustine | DNA Damage/DNA Repair | DNA Alkylator/Crosslinker |
| Thioguanine | Chromatin/Epigenetic | DNA Methyltransferase inhibitor |
| Lorlatinib | Angiogenesis; Immunology/Inflammation; Tyrosine Kinase/Adaptors; Tyrosine Kinase/Adaptors | ALK inhibitor; ROS inhibitor; Tyrosine Kinases inhibitor |
| Olaparib | Chromatin/Epigenetic; DNA Damage/DNA Repair | PARP |
| Batyl alcohol | Immunology/Inflammation | Others |
| Tetrandrine | Membrane transporter/Ion channel | Calcium Channel inhibitor |
| DL-Menthol | Others | Others |
| Arctigenin | Autophagy; Proteases/Proteasome | Autophagy; MMP |
| Berbamine dihydrochloride | Angiogenesis | Bcr-Abl |
| Tanshinone I | Metabolism | Phospholipase inhibitor |
| Andrographolide | NF-Kb | NF-κB inhibitor |
| S-Isocorydine(+) | Others | Others |
| Coenzyme Q10 | Metabolic Enzyme/Protease | Endogenous Metabolite |
| Catharanthine | Neuroscience | AChR antagonist |
| 10-Hydroxycamptothecin | DNA Damage/DNA Repair | Topoisomerase inhibitor |
| Glycyrrhizin | Metabolism | Dehydrogenase inhibitor |
| Sodium Demethylcantharidate | Others | Others |
| Limonin | Microbiology&Virology; Proteases/Proteasome | HIV Protease inhibitor |
| Crenolanib | Angiogenesis; Tyrosine Kinase/Adaptors | PDGFR; FLT |
| Fruquintinib | Angiogenesis; Tyrosine Kinase/Adaptors | VEGFR inhibitor |
| Masitinib | Angiogenesis; Tyrosine Kinase/Adaptors; Cytoskeletal Signaling | Bcr-Abl inhibitor; c-Fms inhibitor; c-Kit inhibitor; Hck inhibitor; PDGFR inhibitor; Src inhibitor |
| Thymopentin | Endocrinology/Hormones | Estrogen/progestogen Receptor agonist |

| | | |
|---|---|---|
| Vismodegib | GPCR/G Protein; Stem Cells | Hedgehog/Smoothened |
| Cabozantinib | Angiogenesis; Tyrosine Kinase/Adaptors | c-Kit; c-Met/HGFR; TAM Receptor; VEGFR |
| Apatinib | Angiogenesis; Apoptosis; Tyrosine Kinase/Adaptors | c-Kit inhibitor; c-RET inhibitor; Src inhibitor; VEGFR inhibitor |
| Chlorotrianisene | Endocrinology/Hormones | Estrogen/progestogen Receptor agonist |
| Hesperetin | GPCR/G Protein; Stem Cells; Neuroscience | 5-HT Receptor inhibitor; TGF-beta/Smad inhibitor |
| Cabazitaxel | Cytoskeletal Signaling | Microtubule Associated inhibitor |
| Ulipristal acetate | Endocrinology/Hormones | Estrogen/progestogen Receptor |
| Dexmedetomidine | GPCR/G Protein | Adrenergic Receptor agonist |
| Mocetinostat | Chromatin/Epigenetic; DNA Damage/DNA Repair | HDAC |
| Tozasertib | Cell Cycle/Checkpoint; Chromatin/Epigenetic | Aurora Kinase |
| Binimetinib | MAPK | MEK inhibitor |
| Cediranib | Angiogenesis; Tyrosine Kinase/Adaptors | c-Kit inhibitor; PDGFR inhibitor; VEGFR inhibitor |
| Osimertinib | Angiogenesis; JAK/STAT signaling; Tyrosine Kinase/Adaptors | EGFR inhibitor |
| Baricitinib | Angiogenesis; Cell Cycle/Checkpoint; Tyrosine Kinase/Adaptors; Chromatin/Epigenetic; JAK/STAT signaling; Stem Cells | Chk inhibitor; JAK inhibitor; Tyrosine Kinases inhibitor |
| Dacomitinib | Angiogenesis; JAK/STAT signaling; Tyrosine Kinase/Adaptors | EGFR inhibitor |
| Tivozanib | Angiogenesis; Tyrosine Kinase/Adaptors | Ephrin Receptor inhibitor; PDGFR inhibitor; VEGFR inhibitor |
| Nedaplatin | DNA Damage/DNA Repair | DNA/RNA Synthesis inhibitor |
| Bortezomib | Proteases/Proteasome; Ubiquitination | Proteasome |
| Pixantrone dimaleate | DNA Damage | Topoisomerase |
| Camostat mesilate | Membrane transporter/Ion channel | Sodium Channel inhibitor |
| Pirfenidone | Stem Cells | TGF-beta/Smad inhibitor |
| Pomalidomide | Apoptosis | TNF |
| Panobinostat | Chromatin/Epigenetic; DNA Damage/DNA Repair | HDAC |
| Vemurafenib | Angiogenesis; MAPK; Neuroscience; Tyrosine Kinase/Adaptors | ACK; MAPK; Raf; Tyrosine Kinases |
| Abemaciclib | Cell Cycle/Checkpoint | CDK inhibitor |

| | | |
|---|---|---|
| Ponatinib | Angiogenesis; Tyrosine Kinase/Adaptors; Cytoskeletal Signaling | Bcr-Abl; c-Kit; FGFR; PDGFR; Src; VEGFR |
| Rociletinib | Angiogenesis; JAK/STAT signaling; Tyrosine Kinase/Adaptors | EGFR inhibitor |
| Tipiracil hydrochloride | DNA Damage/DNA Repair | Nucleoside Antimetabolite/Analog inhibitor |
| Enasidenib | Metabolism | Dehydrogenase inhibitor |
| Apalutamide | Membrane transporter/Ion channel; Endocrinology/Hormones; Neuroscience | Androgen Receptor inhibitor; GABA Receptor inhibitor |
| Menatetrenone | Metabolism | Vitamin inhibitor |
| Radotinib | Angiogenesis; Cytoskeletal Signaling | Bcr-Abl inhibitor |
| Neratinib | Angiogenesis; JAK/STAT signaling; Tyrosine Kinase/Adaptors | EGFR; HER |
| Afatinib | Angiogenesis; JAK/STAT signaling; Tyrosine Kinase/Adaptors | EGFR; HER |
| 2-Methoxyestradiol | Angiogenesis; Cytoskeletal Signaling; Chromatin/Epigenetic | HIF inhibitor; Microtubule Associated inhibitor |
| Cephalomannine | Cytoskeletal Signaling | Microtubule Associated |
| 6-Mercaptopurine monohydrate | DNA Damage/DNA Repair; Others | DNA/RNA Synthesis inhibitor; Others inhibitor |
| Apigenin | Metabolism | P450 inhibitor |
| Fulvestrant | Endocrinology/Hormones | Estrogen/progestogen Receptor |
| Trametinib | MAPK | MEK |
| Ixazomib | Proteases/Proteasome; Ubiquitination | Proteasome inhibitor |
| ABT199 | Apoptosis | BCL |
| Pexidartinib | Angiogenesis; Tyrosine Kinase/Adaptors | c-Kit inhibitor; CSF-1R inhibitor; FLT inhibitor |
| Nimorazole | Others | Others |
| Chidamide | Chromatin/Epigenetic; DNA Damage/DNA Repair; NF-Kb | HDAC inhibitor |
| MLN9708 | Proteases/Proteasome; Ubiquitination | Proteasome inhibitor |
| Fedratinib | Angiogenesis; Apoptosis; Chromatin/Epigenetic; Tyrosine Kinase/Adaptors; JAK/STAT signaling; Stem Cells | JAK; c-RET; FLT |
| Duvelisib | PI3K/Akt/mTOR signaling | PI3K inhibitor |

| | | |
|---|---|---|
| Alectinib | Angiogenesis; Tyrosine Kinase/Adaptors; Tyrosine Kinase/Adaptors | ALK inhibitor; Tyrosine Kinases inhibitor; VEGFR inhibitor |
| NVP-LDE225 | GPCR/G Protein; Stem Cells | Hedgehog/Smoothened antagonist |
| Alpelisib | PI3K/Akt/mTOR signaling | PI3K |
| Dinaciclib | Cell Cycle/Checkpoint | CDK |
| Dabrafenib | MAPK | Raf |
| Idelalisib | PI3K/Akt/mTOR signaling | PI3K |
| Pracinostat | Chromatin/Epigenetic; DNA Damage/DNA Repair; NF-Kb | HDAC inhibitor |
| Belinostat | Chromatin/Epigenetic; DNA Damage/DNA Repair; NF-Kb | HDAC inhibitor |
| Ibrutinib | Angiogenesis; Tyrosine Kinase/Adaptors | BTK; Src; Tyrosine Kinases |
| Ruxolitinib | Angiogenesis; Chromatin/Epigenetic; JAK/STAT signaling; Stem Cells | JAK |
| Cabozantinib Malate | Angiogenesis; Tyrosine Kinase/Adaptors | c-Kit; c-Met/HGFR; TAM Receptor; VEGFR |
| Carfilzomib | Proteases/Proteasome; Ubiquitination | Proteasome inhibitor |
| Regorafenib | Angiogenesis; Apoptosis; MAPK; Tyrosine Kinase/Adaptors | c-Kit; c-RET; Raf; VEGFR |
| LDK378 | Angiogenesis; Tyrosine Kinase/Adaptors | ALK; IGF-1R |
| EPZ6438 | Chromatin/Epigenetic | Histone Methyltransferase inhibitor |
| Palbociclib | Cell Cycle/Checkpoint | CDK inhibitor |
| Everolimus | PI3K/Akt/mTOR signaling | mTOR |
| Nintedanib | Angiogenesis; Tyrosine Kinase/Adaptors | FLT; Src; VEGFR; PDGFR; FGFR |
| Plerixafor | Autophagy; GPCR/G Protein; Immunology/Inflammation | CXCR antagonist |
| Afatinib Dimaleate | Angiogenesis; JAK/STAT signaling; Tyrosine Kinase/Adaptors | EGFR inhibitor; HER inhibitor |
| Torezolid | Metabolism | MAO inhibitor |
| Genistein | Angiogenesis; JAK/STAT signaling; Tyrosine Kinase/Adaptors | EGFR antagonist |
| Vinblastine sulfate | Neuroscience | AChR inhibitor |
| 5-Aminolevulinic acid hydrochloride | Autophagy | Autophagy; Mitophagy |
| Crizotinib | Angiogenesis; Tyrosine Kinase/Adaptors | ALK; c-Met/HGFR |

| | | |
|---|---|---|
| Silibinin | Autophagy | Autophagy |
| Melatonine | Endocrinology/Hormones; GPCR/G Protein; Metabolism; Neuroscience | CaMK agonist; Estrogen/progestogen Receptor antagonist; Melatonin Receptor agonist; MPO inhibitor; ROR agonist |
| Vandetanib | Angiogenesis; JAK/STAT signaling; Tyrosine Kinase/Adaptors | EGFR; VEGFR |
| Pentamidine isethionate | Microbiology&Virology | Antibacterial |
| Lenalidomide | Apoptosis | TNF |
| Beta-Carotene | Metabolism | Vitamin inhibitor |
| Rosiglitazone maleate | Metabolism | PPAR agonist |
| Imatinib Mesylate | Angiogenesis; Tyrosine Kinase/Adaptors; Cytoskeletal Signaling | Bcr-Abl; c-Kit; PDGFR |
| Hydralazine hydrochloride | Metabolism | MAO inhibitor |
| Isotretinoin | Metabolism | Retinoid Receptor inhibitor |
| Lomustine | DNA Damage/DNA Repair; Others | DNA Alkylation inhibitor; Others |
| Doxifluridine | DNA Damage/DNA Repair | DNA/RNA Synthesis antagonist |
| Ibandronate sodium | Microbiology&Virology; Others | HBV inhibitor; Others |
| Cyclocytidine hydrochloride | DNA Damage/DNA Repair | DNA/RNA Synthesis inhibitor |
| Letrozole | Endocrinology/Hormones | Aromatase inhibitor |
| Exemestane | Endocrinology/Hormones | Aromatase inhibitor |
| Vorinostat | Chromatin/Epigenetic; DNA Damage/DNA Repair | HDAC |
| Etoricoxib | Immunology/Inflammation; Neuroscience | COX inhibitor |
| Cisplatin | DNA Damage/DNA Repair | DNA/RNA Synthesis |
| Rebamipide | oxidation-reduction | Free radical scavengers inhibitor |
| Resveratrol | Chromatin/Epigenetic; Immunology/Inflammation; DNA Damage/DNA Repair; Metabolism; NF-Kb; Neuroscience | COX; DNA/RNA Synthesis; IκB/IKK; Lipoxygenase; NADPH; Sirtuin |
| Raloxifene hydrochloride | Endocrinology/Hormones; Metabolism; Others | Estrogen/progestogen Receptor antagonist; MAO antagonist; Others antagonist |
| 8-Methoxypsoralen | DNA Damage/DNA Repair | DNA Alkylation agonist |
| Rapamycin | PI3K/Akt/mTOR signaling; Autophagy | mTOR inhibitor; Autophagy activator |
| Sodium Phenylbutyrate | Chromatin/Epigenetic; DNA Damage/DNA Repair; NF-Kb | HDAC inhibitor |
| Nilotinib | Angiogenesis; Cytoskeletal Signaling | Bcr-Abl |

| | | |
|---|---|---|
| Teniposide | DNA Damage/DNA Repair | Topoisomerase inhibitor |
| Hydroxy Camptothecine | DNA Damage/DNA Repair | Topoisomerase inhibitor |
| Daunorubicin hydrochloride | DNA Damage/DNA Repair | DNA/RNA Synthesis inhibitor |
| Decitabine | Chromatin/Epigenetic | DNA Methyltransferase |
| Streptozocin | DNA Damage/DNA Repair | DNA Alkylation inducer |
| Vidarabine | DNA Damage/DNA Repair; Tyrosine Kinase/Adaptors | DNA/RNA Synthesis inhibitor; Tyrosine Kinases |
| Methotrexate | Metabolism | Dehydrogenase inhibitor |
| AICAR (Acadesine) | PI3K/Akt/mTOR signaling | AMPK activator |
| Trilostane | Metabolism | Dehydrogenase inhibitor |
| Toremifene citrate | Endocrinology/Hormones | Estrogen/progestogen Receptor modulator |
| Axitinib | Angiogenesis; Tyrosine Kinase/Adaptors | PDGFR; VEGFR; c-Kit |
| Dasatinib monohydrate | Angiogenesis; Tyrosine Kinase/Adaptors; Cytoskeletal Signaling | Bcr-Abl inhibitor; c-Kit inhibitor; Ephrin Receptor inhibitor; Src inhibitor |
| Dasatinib | Angiogenesis; Tyrosine Kinase/Adaptors; Cytoskeletal Signaling | Bcr-Abl; c-Kit; Src |
| Trifluridine | DNA Damage/DNA Repair | DNA/RNA Synthesis inhibitor |
| Mechlorethamine hydrochloride | DNA Damage/DNA Repair | DNA/RNA Synthesis inhibitor |
| Capecitabine | DNA Damage/DNA Repair | DNA/RNA Synthesis inhibitor |
| Chlorpromazine hydrochloride | GPCR/G Protein; Neuroscience | 5-HT Receptor antagonist; Dopamine Receptor antagonist |
| Tegafur | DNA Damage/DNA Repair | DNA/RNA Synthesis inhibitor |
| 5-Azacytidine | Chromatin/Epigenetic | DNA Methyltransferase inhibitor |
| Pamidronate disodium salt | Microbiology&Virology; Others | HBV; Others |
| Carmofur | DNA Damage/DNA Repair | DNA/RNA Synthesis inhibitor |
| Anethole trithione | Others | Others |
| Megestrol acetate | Endocrinology/Hormones; Metabolism | Estrogen/progestogen Receptor agonist; GR inhibitor |
| Cytarabine | DNA Damage/DNA Repair | DNA/RNA Synthesis inhibitor |
| Ubenimex | Immunology/Inflammation; Others | LTR inhibitor; Others inhibitor |
| Triethylenethiophosphoramide | DNA Damage/DNA Repair | DNA Alkylation |
| Altretamine | DNA Damage/DNA Repair | DNA Alkylation |
| Sodium etidronate | Proteases/Proteasome | Tyrosinase inhibitor |
| Amsacrine | DNA Damage/DNA Repair; Membrane transporter/Ion channel | Potassium Channel inhibitor; Topoisomerase inhibitor |
| Mitotan | Neuroscience | AChR |

| | | |
|---|---|---|
| Rofecoxib | Immunology/Inflammation; Neuroscience | COX inhibitor |
| Retinol | Vitamin | VA |
| Gefitinib | Angiogenesis; JAK/STAT signaling; Tyrosine Kinase/Adaptors | EGFR |
| Temozolomide | DNA Damage/DNA Repair | DNA/RNA Synthesis |
| Topotecan hydrochloride | DNA Damage/DNA Repair | Topoisomerase inhibitor |
| Cyproterone acetate | Endocrinology/Hormones | Androgen Receptor antagonist |
| Phenoxybenzamine hydrochloride | GPCR/G Protein; Neuroscience | Adrenergic Receptor antagonist; CaMK inhibitor |
| Mebendazole | Cytoskeletal Signaling | Microtubule Associated inhibitor |
| Nifedipine | Membrane transporter/Ion channel; Neuroscience | Calcium Channel inhibitor; CaMK inhibitor; Potassium Channel inhibitor |
| Dacarbazine | DNA Damage/DNA Repair | DNA/RNA Synthesis inhibitor |
| Ciclopirox ethanolamine | Membrane transporter/Ion channel; Others | ATPase inhibitor; Others |
| Aminoglutethimide | Endocrinology/Hormones | Aromatase inhibitor |
| Mifepristone | Endocrinology/Hormones; Metabolism | Estrogen/progestogen Receptor antagonist; GR antagonist |
| Formestane | Endocrinology/Hormones | Aromatase inhibitor |
| Tolnaftate | GPCR/G Protein; Stem Cells | Hedgehog/Smoothened inhibitor |
| Vitamin D2 | DNA Damage/DNA Repair; Metabolism | DNA/RNA Synthesis inhibitor; Vitamin inhibitor |
| Apramycin sulfate | Microbiology&Virology | Antibacterial inhibitor |
| Dexamethasone | Endocrinology/Hormones; GPCR/G Protein; Immunology/Inflammation; Metabolism | GR agonist; IL Receptor modulator; |
| Carboplatin | DNA Damage/DNA Repair | DNA/RNA Synthesis inhibitor |
| Ifosfamide | DNA Damage/DNA Repair | DNA/RNA Synthesis inhibitor |
| Tretinoin | Metabolism | Retinoid Receptor agonist |
| Granisetron hydrochloride | GPCR/G Protein; Neuroscience | 5-HT Receptor antagonist |
| Fludarabine | JAK/STAT signaling; Stem Cells; DNA Damage/DNA Repair | STAT; DNA/RNA Synthesis |
| Docetaxel | Cytoskeletal Signaling | Microtubule Associated |
| Tranylcypromine (2-PCPA) hydrochloride | Metabolism | MAO inhibitor |
| Roflumilast | Metabolism | PDE inhibitor |
| Doxorubicin hydrochloride | DNA Damage/DNA Repair | Topoisomerase |
| Itraconazole | Metabolism | P450 inhibitor |
| Gimeracil | Autophagy | Autophagy inhibitor |

| | | |
|---|---|---|
| Fluorocytosine | DNA Damage/DNA Repair | DNA/RNA Synthesis inhibitor |
| Fluorouracil | DNA Damage/DNA Repair | DNA/RNA Synthesis |
| d-penicillamin | Others | Others |
| Noscapine hydrochloride | Endocrinology/Hormones; GPCR/G Protein; Neuroscience | Opioid Receptor agonist |
| Chloroambucil | DNA Damage/DNA Repair | DNA Alkylation inhibitor |
| Paclitaxel | Cytoskeletal Signaling | Microtubule |
| Floxuridine | DNA Damage/DNA Repair | DNA/RNA Synthesis inhibitor |
| Dexamethason acetate | GPCR/G Protein; Immunology/Inflammation; Metabolism | Annexin A; GR; IL Receptor modulator; NOS modulator |
| Nicotinamide | Chromatin/Epigenetic; DNA Damage/DNA Repair | Sirtuin inhibitor |
| Busulfan | DNA Damage/DNA Repair | DNA Alkylation |
| Clioquinol | Microbiology&Virology | Antibiotic |
| Tamoxifen Z-isomer citrate | Endocrinology/Hormones | Estrogen/progestogen Receptor agonist |
| Propranolol hydrochloride | GPCR/G Protein | Adrenergic Receptor antagonist |
| Rutin | GPCR/G Protein | Prostaglandin Receptor inhibitor |
| Isoliquiritigenin | Endocrinology/Hormones; Enzyme | Reductase inhibitor |
| Brivudine | Microbiology&Virology | HSV |
| L-Cycloserine | Membrane Transporter/Ion Channel; Neuroscience | GABA Receptor |
| Cyclophosphamide monohydrate | DNA Damage/DNA Repair; Immunology/Inflammation | DNA Alkylation inducer; MRP inhibitor |
| Gabapentin | Membrane transporter/Ion channel | GABA Receptor |
| Rifampicin | DNA Damage/DNA Repair; Microbiology&Virology | Antifection inhibitor; DNA/RNA Synthesis inhibitor |
| Hydroxyurea | DNA Damage/DNA Repair | DNA/RNA Synthesis inhibitor |
| Piceatannol | Apoptosis; Cytoskeletal Signaling; GPCR/G Protein | PKA inhibitor; PKC inhibitor; Serine/threonin kinase inhibitor |
| Naringin | Metabolism | P450 antagonist |
| Lenvatinib | Angiogenesis; Tyrosine Kinase/Adaptors | FGFR inhibitor; PDGFR inhibitor; VEGFR inhibitor |
| Flutamide | Endocrinology/Hormones | Androgen Receptor antagonist |
| Irinotecan hydrochloride trihydrate | DNA Damage/DNA Repair | Topoisomerase inhibitor |
| Celecoxib | Immunology/Inflammation; Neuroscience | COX inhibitor |
| Sulindac | Immunology/Inflammation; Neuroscience | COX inhibitor |
| Artesunate | JAK/STAT signaling; Stem Cells | STAT inhibitor |

| | | |
|---|---|---|
| Bicalutamide | Endocrinology/Hormones | Androgen Receptor antagonist |
| Sunitinib Malate | Angiogenesis; Tyrosine Kinase/Adaptors | c-Kit; FLT; PDGFR; VEGFR |
| Erlotinib | Angiogenesis; JAK/STAT signaling; Tyrosine Kinase/Adaptors | EGFR |
| Etidronate | Microbiology&Virology; Others | HBV antagonist; Others inhibitor |
| Clofarabine | DNA Damage/DNA Repair | DNA/RNA Synthesis inhibitor |
| Meticrane | Membrane Transporter/Ion Channel | Chloride Channel; Sodium Channel |
| Nilutamide | Endocrinology/Hormones | Androgen Receptor antagonist |
| Hydrocortisone butyrate | GPCR/G Protein; Metabolism | Annexin A; GR |
| Chloropyramine hydrochloride | GPCR/G Protein; Angiogenesis; Cytoskeletal Signaling; Immunology/Inflammation; Neuroscience | FAK; Histamine Receptor antagonist |
| Tilorone dihydrochloride | Others | Others |
| Lonidamine | Others | Others |
| Thalidomide | Apoptosis | TNF inhibitor |
| Pemetrexed disodium | DNA Damage/DNA Repair; Metabolism | DHFR inhibitor; DNA/RNA Synthesis inhibitor |
| Pemetrexed acid | DNA Damage/DNA Repair; Metabolism | DHFR inhibitor; DNA/RNA Synthesis inhibitor |
| Docetaxel trihydrate | Apoptosis | BCL antagonist; Microtubule Associated inhibitor |
| Oxaliplatin | DNA Damage/DNA Repair | DNA/RNA Synthesis inhibitor |
| Bosutinib | Angiogenesis; Cytoskeletal Signaling | Bcr-Abl; Src |
| Toremifene | Endocrinology/Hormones | Estrogen/progestogen Receptor modulator |
| Imiquimod | Immunology/Inflammation | TLR agonist |
| Etoposide | DNA Damage/DNA Repair | Topoisomerase |
| Cepharanthine | Apoptosis | TNF |
| Epirubicin hydrochloride | DNA Damage/DNA Repair | Topoisomerase inhibitor |
| Pazopanib | Angiogenesis; Tyrosine Kinase/Adaptors | c-Kit inhibitor; PDGFR inhibitor; VEGFR inhibitor |
| Bendamustine hydrochloride | DNA Damage/DNA Repair | DNA/RNA Synthesis inhibitor |
| Sorafenib | Angiogenesis; MAPK; Tyrosine Kinase/Adaptors | c-Kit; FLT; PDGFR; Raf; VEGFR |
| Sorafenib tosylate | Angiogenesis; MAPK; Tyrosine Kinase/Adaptors | c-Kit; FLT; PDGFR; Raf; VEGFR |
| Lapatinib | Angiogenesis; MAPK; JAK/STAT signaling; Tyrosine Kinase/Adaptors | EGFR |

| | | |
|---|---|---|
| Uracil | Others | Others |
| Fluphenazine hydrochloride | GPCR/G Protein; Neuroscience | Dopamine Receptor antagonist |
| Disulfiram | Metabolism | Dehydrogenase inhibitor |
| Urethane | Membrane transporter/Ion channel; Neuroscience | AChR inhibitor; Chloride channel inhibitor; GABA Receptor inhibitor; GluR antagonist; NMDAR inhibitor |
| Quinestrol | Endocrinology/Hormones | Estrogen/progestogen Receptor agonist |
| Mercaptopurine | Metabolism; Others | Dehydrogenase inhibitor; Others |

**Supplementary Information Table 2 | Table of the 756 non-redundant image-based quantitative features (and mean value) for morphological profiling in the drug screen.**

| Feature | Average |
|---|---|
| MedianCytoplasmIntensityMeanIntensityW5 | 0.035 |
| CountCells | 404.226 |
| MedianCellsNeighborsSecondClosestDistance1 | 60.092 |
| MedianCellsNeighborsAngleBetweenNeighbors1 | 92.78 |
| MedianCellsNeighborsFirstClosestDistance1 | 42.726 |
| MedianCytoplasmCorrelationCostesW5W4 | 1 |
| MedianCytoplasmTextureContrastW5301256 | 3.985 |
| MedianCytoplasmIntensityMeanIntensityW4 | 0.053 |
| MedianCytoplasmIntensityUpperQuartileIntensityW5 | 0.043 |
| MedianCytoplasmIntensityMedianIntensityW5 | 0.031 |
| MedianCytoplasmIntensityMeanIntensityW2 | 0.063 |
| MedianCytoplasmTextureSumAverageW2300256 | 32.532 |
| MedianCytoplasmTextureSumVarianceW5300256 | 55.361 |
| MedianCytoplasmIntensityMaxIntensityEdgeW4 | 0.115 |
| MedianCytoplasmIntensityMaxIntensityW4 | 0.121 |
| MedianCellsAreaShapeMajorAxisLength | 77.751 |
| MedianCellsAreaShapeMaxFeretDiameter | 83.921 |
| MedianCytoplasmTextureDifferenceEntropyW2300256 | 2.461 |
| MedianCytoplasmTextureDifferenceEntropyW2301256 | 2.757 |
| MedianCellsAreaShapeArea | 2295.887 |
| MedianCellsAreaShapeEquivalentDiameter | 53.611 |
| MedianCytoplasmTextureEntropyW5300256 | 5.304 |
| MedianCytoplasmTextureSumEntropyW5300256 | 4.15 |
| MedianCytoplasmTextureDifferenceEntropyW2303256 | 2.748 |
| MedianCytoplasmTextureDifferenceEntropyW2302256 | 2.481 |

| | |
|---|---|
| MedianCytoplasmTextureInfoMeas2W2300256 | 0.96 |
| MedianCytoplasmTextureInfoMeas2W2301256 | 0.938 |
| MedianCytoplasmTextureDifferenceEntropyW4301256 | 2.491 |
| MedianCytoplasmTextureDifferenceEntropyW4303256 | 2.484 |
| MedianCytoplasmTextureInverseDifferenceMomentW5301 | 0.543 |
| MedianCytoplasmTextureInverseDifferenceMomentW5302 | 0.604 |
| MedianCytoplasmTextureAngularSecondMomentW5300256 | 0.054 |
| MedianCytoplasmTextureDifferenceVarianceW5300256 | 0.016 |
| MedianCellsAreaShapeMinFeretDiameter | 47.831 |
| MedianCellsAreaShapeMinorAxisLength | 43.709 |
| MedianCytoplasmTextureDifferenceEntropyW5301256 | 1.991 |
| MedianCytoplasmTextureDifferenceEntropyW5302256 | 1.761 |
| MedianCytoplasmIntensityStdIntensityEdgeW5 | 0.015 |
| MedianCytoplasmIntensityStdIntensityW5 | 0.015 |
| MedianCytoplasmIntensityMedianIntensityW4 | 0.048 |
| MedianCytoplasmTextureDifferenceEntropyW5300256 | 1.741 |
| MedianCytoplasmTextureDifferenceEntropyW5303256 | 1.984 |
| MedianCytoplasmTextureSumVarianceW2300256 | 126.342 |
| MedianCytoplasmTextureSumVarianceW2303256 | 120.039 |
| MedianCytoplasmTextureEntropyW2300256 | 6.742 |
| MedianCytoplasmTextureSumEntropyW2300256 | 4.888 |
| MedianCytoplasmRadialDistributionZernikeMagnitudeW79 | 0.001 |
| MedianCytoplasmRadialDistributionZernikeMagnitudeW89 | 0.001 |
| MedianCellsGranularity16W4 | 0.327 |
| MedianCytoplasmTextureInverseDifferenceMomentW5303 | 0.544 |
| MedianCytoplasmRadialDistributionZernikeMagnitudeW71 | 0.001 |
| MedianCellsAreaShapeConvexArea | 3075.382 |
| MedianCytoplasmTextureInverseDifferenceMomentW5300 | 0.612 |
| MedianCytoplasmIntensityMedianIntensityW2 | 0.059 |
| MedianCytoplasmTextureInverseDifferenceMomentW2300 | 0.434 |
| MedianCytoplasmTextureInverseDifferenceMomentW2303 | 0.37 |
| MedianCytoplasmIntensityMeanIntensityEdgeW5 | 0.035 |
| MedianCytoplasmTextureContrastW2301256 | 11.986 |
| MedianCytoplasmTextureContrastW2302256 | 7.465 |
| MedianCytoplasmTextureContrastW4301256 | 8.667 |
| MedianCytoplasmTextureContrastW4303256 | 8.55 |
| MedianCellsGranularity15W4 | 0.323 |
| MedianCytoplasmRadialDistributionMeanFracW24of5 | 0.798 |
| MedianCytoplasmRadialDistributionMeanFracW44of5 | 0.751 |

| | |
|---|---|
| MedianCytoplasmTextureInverseDifferenceMomentW2302 | 0.429 |
| MedianCytoplasmIntensityLowerQuartileIntensityW5 | 0.022 |
| MedianCytoplasmTextureContrastW4300256 | 5.264 |
| MedianCytoplasmTextureDifferenceEntropyW4302256 | 2.3 |
| MedianCytoplasmTextureInverseDifferenceMomentW2301 | 0.368 |
| MedianCytoplasmTextureEntropyW4300256 | 5.937 |
| MedianCytoplasmTextureSumEntropyW4300256 | 4.242 |
| MedianCytoplasmTextureDifferenceEntropyW4300256 | 2.222 |
| MedianCellsGranularity16W3 | 0.331 |
| MedianCytoplasmRadialDistributionZernikeMagnitudeW78 | 0.001 |
| MedianCytoplasmRadialDistributionZernikeMagnitudeW88 | 0.001 |
| MedianCytoplasmIntensityUpperQuartileIntensityW2 | 0.078 |
| MedianCytoplasmTextureAngularSecondMomentW2300256 | 0.021 |
| MedianCytoplasmTextureDifferenceVarianceW2300256 | 0.007 |
| MedianCytoplasmTextureInverseDifferenceMomentW4301 | 0.431 |
| MedianCytoplasmTextureInverseDifferenceMomentW4302 | 0.476 |
| MedianCytoplasmTextureInverseDifferenceMomentW4300 | 0.493 |
| MedianCytoplasmTextureInverseDifferenceMomentW4303 | 0.432 |
| MedianCytoplasmRadialDistributionZernikeMagnitudeW82 | 0.001 |
| MedianCytoplasmRadialDistributionZernikeMagnitudeW87 | 0.001 |
| MedianCytoplasmRadialDistributionZernikeMagnitudeW38 | 0.004 |
| MedianCytoplasmRadialDistributionZernikeMagnitudeW45 | 0.002 |
| MedianCytoplasmTextureContrastW4302256 | 6.109 |
| MedianCellsGranularity15W3 | 0.333 |
| MedianCytoplasmRadialDistributionZernikeMagnitudeW54 | 0.001 |
| MedianCytoplasmRadialDistributionZernikeMagnitudeW77 | 0.002 |
| MedianCytoplasmRadialDistributionZernikeMagnitudeW86 | 0.001 |
| MedianCellsAreaShapePerimeter | 277.214 |
| MedianCellsGranularity7W1 | 4.885 |
| MedianCytoplasmRadialDistributionZernikeMagnitudeW74 | 0.002 |
| MedianCytoplasmRadialDistributionZernikeMagnitudeW83 | 0.001 |
| MedianCytoplasmRadialDistributionZernikeMagnitudeW63 | 0.006 |
| MedianCytoplasmRadialDistributionZernikeMagnitudeW33 | 0.009 |
| MedianCytoplasmRadialDistributionRadialCVW24of5 | 0.127 |
| MedianCytoplasmRadialDistributionRadialCVW54of5 | 0.152 |
| MedianCytoplasmIntensityLowerQuartileIntensityW4 | 0.039 |
| MedianCytoplasmIntensityMeanIntensityEdgeW4 | 0.055 |
| MedianCytoplasmRadialDistributionZernikeMagnitudeW76 | 0.002 |
| MedianCytoplasmRadialDistributionZernikeMagnitudeW85 | 0.001 |

| | |
|---|---|
| MedianCytoplasmRadialDistributionZernikeMagnitudeW68 | 0.002 |
| MedianCytoplasmRadialDistributionZernikeMagnitudeW67 | 0.004 |
| MedianCytoplasmRadialDistributionZernikeMagnitudeW69 | 0.003 |
| MedianCytoplasmRadialDistributionRadialCVW25of5 | 0.218 |
| MedianCytoplasmRadialDistributionRadialCVW55of5 | 0.273 |
| MedianCytoplasmRadialDistributionZernikeMagnitudeW73 | 0.002 |
| MedianCellsAreaShapeMaximumRadius | 17.501 |
| MedianCellsAreaShapeMeanRadius | 6.078 |
| MedianCytoplasmRadialDistributionZernikeMagnitudeW65 | 0.002 |
| MedianCytoplasmIntensityStdIntensityEdgeW2 | 0.022 |
| MedianCytoplasmIntensityStdIntensityW2 | 0.023 |
| MedianCytoplasmRadialDistributionZernikeMagnitudeW70 | 0.002 |
| MedianCytoplasmRadialDistributionZernikeMagnitudeW84 | 0.001 |
| MedianCellsGranularity5W1 | 14.443 |
| MedianCellsGranularity6W1 | 10.111 |
| MedianCytoplasmRadialDistributionZernikeMagnitudeW61 | 0.006 |
| MedianCytoplasmRadialDistributionZernikeMagnitudeW64 | 0.005 |
| MedianCytoplasmTextureInfoMeas1W2300256 | -0.329 |
| MedianCytoplasmTextureInfoMeas1W2302256 | -0.325 |
| MedianCytoplasmTextureContrastW2300256 | 7.219 |
| MedianCytoplasmTextureContrastW2303256 | 11.803 |
| MedianCytoplasmRadialDistributionZernikeMagnitudeW81 | 0.002 |
| MedianCellsGranularity8W1 | 2.267 |
| MedianCellsGranularity7W3 | 2.331 |
| MedianCytoplasmRadialDistributionRadialCVW44of5 | 0.09 |
| MedianCellsGranularity1W3 | 25.718 |
| MedianCytoplasmIntensityStdIntensityW4 | 0.016 |
| MedianCytoplasmTextureSumVarianceW4300256 | 58.985 |
| MedianCellsGranularity1W1 | 13.632 |
| MedianCytoplasmRadialDistributionRadialCVW45of5 | 0.165 |
| MedianCytoplasmTextureCorrelationW5300256 | 0.891 |
| MedianCytoplasmTextureCorrelationW5302256 | 0.891 |
| MedianCellsGranularity7W4 | 1.995 |
| MedianCellsGranularity4W1 | 10.7 |
| MedianCellsGranularity1W4 | 34.317 |
| MedianCellsGranularity6W3 | 2.725 |
| MedianCytoplasmCorrelationMandersW2W5 | 0.962 |
| MedianCytoplasmCorrelationMandersW4W5 | 0.954 |
| MedianCellsGranularity8W3 | 1.809 |

| | |
|---|---|
| MedianCellsGranularity9W1 | 1.11 |
| MedianCytoplasmIntensityStdIntensityEdgeW4 | 0.018 |
| MedianCytoplasmRadialDistributionZernikeMagnitudeW41 | 0.002 |
| MedianCytoplasmRadialDistributionZernikeMagnitudeW49 | 0.001 |
| MedianCellsGranularity6W4 | 2.401 |
| MedianCytoplasmTextureInfoMeas1W2301256 | -0.272 |
| MedianCytoplasmTextureInfoMeas1W2303256 | -0.274 |
| MedianCytoplasmTextureInfoMeas1W5300256 | -0.389 |
| MedianCytoplasmTextureInfoMeas1W5302256 | -0.385 |
| MedianCellsGranularity10W1 | 0.58 |
| MedianCytoplasmRadialDistributionZernikeMagnitudeW59 | 0.001 |
| MedianCellsGranularity16W1 | 0.052 |
| MedianCytoplasmCorrelationKW5W4 | 1.508 |
| MedianCytoplasmTextureInfoMeas2W4302256 | 0.883 |
| MedianCytoplasmTextureInfoMeas2W4303256 | 0.845 |
| MedianCytoplasmRadialDistributionZernikeMagnitudeW72 | 0.002 |
| MedianCytoplasmIntensityMeanIntensityEdgeW2 | 0.061 |
| MedianCytoplasmCorrelationKW2W5 | 0.586 |
| MedianCytoplasmRadialDistributionZernikeMagnitudeW80 | 0.001 |
| MedianCytoplasmTextureInfoMeas1W5303256 | -0.326 |
| MedianCytoplasmTextureInfoMeas2W5300256 | 0.953 |
| MedianCytoplasmCorrelationRWCW2W4 | 0.879 |
| MedianCytoplasmCorrelationRWCW4W2 | 0.876 |
| MedianCytoplasmTextureInfoMeas2W4300256 | 0.89 |
| MedianCytoplasmRadialDistributionFracAtDW23of5 | 0 |
| MedianCytoplasmRadialDistributionMeanFracW23of5 | 0.001 |
| MedianCytoplasmIntensityLowerQuartileIntensityW2 | 0.045 |
| MedianCytoplasmIntensityMADIntensityW2 | 0.014 |
| MedianCytoplasmTextureInfoMeas1W5301256 | -0.324 |
| MedianCytoplasmRadialDistributionZernikeMagnitudeW66 | 0.004 |
| MedianCytoplasmTextureCorrelationW2300256 | 0.876 |
| MedianCytoplasmTextureCorrelationW2303256 | 0.8 |
| MedianCytoplasmIntensityMADIntensityW5 | 0.008 |
| MedianCytoplasmCorrelationKW5W2 | 1.789 |
| MedianCellsGranularity5W3 | 2.76 |
| MedianCytoplasmRadialDistributionZernikeMagnitudeW35 | 0.003 |
| MedianCytoplasmRadialDistributionZernikeMagnitudeW44 | 0.003 |
| MedianCytoplasmRadialDistributionZernikeMagnitudeW53 | 0.002 |
| MedianCytoplasmTextureInfoMeas2W4301256 | 0.843 |

| | |
|---|---|
| MedianCytoplasmTextureCorrelationW5301256 | 0.829 |
| MedianCellsGranularity5W4 | 2.622 |
| MedianCytoplasmTextureCorrelationW2302256 | 0.874 |
| MedianCellsGranularity15W1 | 0.062 |
| MedianCytoplasmRadialDistributionZernikeMagnitudeW15 | 0.003 |
| MedianCytoplasmRadialDistributionZernikeMagnitudeW8 | 0.005 |
| MedianCytoplasmRadialDistributionZernikeMagnitudeW31 | 0.008 |
| MedianCytoplasmRadialDistributionZernikeMagnitudeW34 | 0.007 |
| MedianCytoplasmRadialDistributionZernikeMagnitudeW40 | 0.003 |
| MedianCytoplasmTextureCorrelationW5303256 | 0.832 |
| MedianCytoplasmRadialDistributionZernikeMagnitudeW48 | 0.002 |
| MedianCytoplasmRadialDistributionZernikeMagnitudeW58 | 0.001 |
| MedianCytoplasmIntensityMaxIntensityEdgeW5 | 0.082 |
| MedianCytoplasmRadialDistributionFracAtDW24of5 | 0.104 |
| MedianCytoplasmRadialDistributionZernikeMagnitudeW62 | 0.009 |
| MedianCellsGranularity4W4 | 2.583 |
| MedianCellsGranularity11W1 | 0.313 |
| MedianCytoplasmRadialDistributionZernikeMagnitudeW24 | 0.002 |
| MedianCytoplasmTextureContrastW5300256 | 2.347 |
| MedianCytoplasmRadialDistributionZernikeMagnitudeW37 | 0.006 |
| MedianCytoplasmRadialDistributionZernikeMagnitudeW3 | 0.011 |
| MedianCellsGranularity4W3 | 2.392 |
| MedianCellsAreaShapeCompactness | 2.68 |
| MedianCellsAreaShapeFormFactor | 0.395 |
| MedianCytoplasmIntensityMinIntensityEdgeW5 | 0.01 |
| MedianCytoplasmRadialDistributionZernikeMagnitudeW39 | 0.004 |
| MedianCytoplasmRadialDistributionZernikeMagnitudeW46 | 0.003 |
| MedianCytoplasmTextureCorrelationW2301256 | 0.796 |
| MedianCytoplasmRadialDistributionZernikeMagnitudeW19 | 0.001 |
| MedianCytoplasmRadialDistributionZernikeMagnitudeW29 | 0.001 |
| MedianCytoplasmRadialDistributionZernikeMagnitudeW43 | 0.003 |
| MedianCytoplasmRadialDistributionZernikeMagnitudeW47 | 0.002 |
| MedianCytoplasmRadialDistributionZernikeMagnitudeW56 | 0.002 |
| MedianCytoplasmTextureInfoMeas1W4302256 | -0.24 |
| MedianCytoplasmTextureInfoMeas1W4303256 | -0.197 |
| MedianCytoplasmIntensityMADIntensityW4 | 0.009 |
| MedianCytoplasmTextureInfoMeas1W4300256 | -0.253 |
| MedianCytoplasmRadialDistributionZernikeMagnitudeW55 | 0.002 |
| MedianCytoplasmIntensityMaxIntensityEdgeW2 | 0.128 |

| | |
|---|---|
| MedianCellsGranularity3W1 | 5.575 |
| MedianCytoplasmRadialDistributionZernikeMagnitudeW57 | 0.002 |
| MedianCytoplasmRadialDistributionZernikeMagnitudeW51 | 0.002 |
| MedianCellsAreaShapeSolidity | 0.77 |
| MedianCytoplasmTextureCorrelationW4300256 | 0.785 |
| MedianCytoplasmTextureCorrelationW4303256 | 0.68 |
| MedianCytoplasmRadialDistributionZernikeMagnitudeW52 | 0.002 |
| MedianCytoplasmTextureCorrelationW4302256 | 0.765 |
| MedianCellsGranularity12W4 | 0.444 |
| MedianCytoplasmRadialDistributionZernikeMagnitudeW1 | 0.011 |
| MedianCytoplasmRadialDistributionZernikeMagnitudeW4 | 0.009 |
| MedianCytoplasmRadialDistributionZernikeMagnitudeW11 | 0.002 |
| MedianCellsAreaShapeExtent | 0.508 |
| MedianCellsGranularity3W4 | 2.331 |
| MedianCytoplasmRadialDistributionZernikeMagnitudeW5 | 0.004 |
| MedianCytoplasmIntensityIntegratedIntensityW2 | 93.223 |
| MedianCytoplasmIntensityIntegratedIntensityW4 | 86.942 |
| MedianCellsAreaShapeZernike00 | 0.428 |
| MedianCytoplasmTextureCorrelationW4301256 | 0.674 |
| MedianCytoplasmIntensityIntegratedIntensityEdgeW5 | 11.588 |
| MedianCytoplasmRadialDistributionZernikeMagnitudeW14 | 0.004 |
| MedianCytoplasmRadialDistributionZernikeMagnitudeW23 | 0.002 |
| MedianCytoplasmIntensityMinIntensityEdgeW2 | 0.026 |
| MedianCytoplasmRadialDistributionZernikeMagnitudeW17 | 0.003 |
| MedianCytoplasmRadialDistributionZernikeMagnitudeW26 | 0.002 |
| MedianCellsAreaShapeMedianRadius | 5.183 |
| MedianCytoplasmRadialDistributionZernikeMagnitudeW10 | 0.004 |
| MedianCytoplasmCorrelationKW4W5 | 0.72 |
| MedianCytoplasmCorrelationKW2W4 | 0.828 |
| MedianCytoplasmCorrelationKW4W2 | 1.278 |
| MedianCytoplasmRadialDistributionZernikeMagnitudeW18 | 0.003 |
| MedianCytoplasmRadialDistributionZernikeMagnitudeW28 | 0.002 |
| MedianCytoplasmIntensityMinIntensityEdgeW4 | 0.027 |
| MedianCytoplasmTextureSumVarianceW2301256 | 119.661 |
| MedianCellsGranularity2W1 | 2.742 |
| MedianCytoplasmRadialDistributionZernikeMagnitudeW36 | 0.005 |
| MedianCytoplasmRadialDistributionZernikeMagnitudeW16 | 0.003 |
| MedianCytoplasmRadialDistributionZernikeMagnitudeW25 | 0.002 |
| MedianCytoplasmRadialDistributionZernikeMagnitudeW7 | 0.007 |

| | |
|---|---|
| MedianCytoplasmRadialDistributionZernikeMagnitudeW9 | 0.005 |
| MedianCytoplasmTextureAngularSecondMomentW4300256 | 0.048 |
| MedianCellsGranularity8W4 | 1.523 |
| MedianCytoplasmRadialDistributionZernikeMagnitudeW22 | 0.002 |
| MedianCytoplasmRadialDistributionZernikeMagnitudeW13 | 0.004 |
| MedianCytoplasmRadialDistributionZernikeMagnitudeW21 | 0.003 |
| MedianCytoplasmRadialDistributionZernikeMagnitudeW27 | 0.002 |
| MedianCellsGranularity3W3 | 1.92 |
| MedianCytoplasmRadialDistributionZernikeMagnitudeW42 | 0.003 |
| MedianCytoplasmIntensityIntegratedIntensityEdgeW4 | 19.343 |
| MedianCytoplasmIntensityIntegratedIntensityW5 | 52.504 |
| MedianCytoplasmRadialDistributionZernikeMagnitudeW50 | 0.002 |
| MedianCellsGranularity12W1 | 0.181 |
| MedianCellsGranularity13W1 | 0.113 |
| MedianCytoplasmIntensityIntegratedIntensityEdgeW2 | 20.214 |
| MedianCytoplasmIntensityMassDisplacementW2 | 3.242 |
| MedianCytoplasmIntensityMassDisplacementW5 | 4.519 |
| MedianCellsGranularity2W4 | 2.029 |
| MedianCytoplasmCorrelationCostesW2W4 | 1 |
| MedianCellsGranularity2W3 | 1.708 |
| MedianCellsAreaShapeEccentricity | 0.799 |
| MedianCytoplasmRadialDistributionZernikeMagnitudeW32 | 0.014 |
| MedianCytoplasmRadialDistributionZernikeMagnitudeW12 | 0.004 |
| MedianCytoplasmTextureInfoMeas1W4301256 | -0.195 |
| MedianCellsGranularity10W3 | 0.951 |
| MedianCytoplasmTextureContrastW5303256 | 3.859 |
| MedianCytoplasmRadialDistributionZernikeMagnitudeW20 | 0.002 |
| MedianCytoplasmCorrelationRWCW2W5 | 0.827 |
| MedianCytoplasmCorrelationRWCW5W2 | 0.891 |
| MedianCytoplasmRadialDistributionMeanFracW25of5 | 0.996 |
| MedianCytoplasmRadialDistributionMeanFracW55of5 | 0.995 |
| MedianCytoplasmRadialDistributionZernikeMagnitudeW2 | 0.017 |
| MedianCytoplasmRadialDistributionZernikeMagnitudeW6 | 0.007 |
| MedianCytoplasmIntensityMassDisplacementW4 | 2.324 |
| MedianCellsGranularity9W3 | 1.312 |
| MedianCellsGranularity10W4 | 0.778 |
| MedianCytoplasmCorrelationCostesW4W2 | 1 |
| MedianCytoplasmCorrelationRWCW5W4 | 0.906 |
| MedianCytoplasmCorrelationCorrelationW4W5 | 0.65 |

| | |
|---|---|
| MedianCytoplasmCorrelationCorrelationW2W4 | 0.533 |
| MedianCellsNeighborsNumberOfNeighbors1 | 3.286 |
| MedianCellsNeighborsPercentTouching1 | 62.91 |
| MedianCytoplasmCorrelationCostesW5W2 | 1 |
| MedianCellsGranularity9W4 | 1.088 |
| MedianCytoplasmCorrelationRWCW4W5 | 0.835 |
| MedianCytoplasmCorrelationCostesW2W5 | 1 |
| MedianCytoplasmCorrelationCorrelationW2W5 | 0.748 |
| MedianCellsAreaShapeZernike77 | 0.005 |
| MedianCellsAreaShapeZernike99 | 0.003 |
| MedianCellsAreaShapeZernike42 | 0.03 |
| MedianCytoplasmCorrelationOverlapW2W5 | 0.97 |
| MedianCellsAreaShapeZernike22 | 0.053 |
| MedianCytoplasmCorrelationOverlapW2W4 | 0.954 |
| MedianCytoplasmCorrelationOverlapW4W5 | 0.958 |
| MedianCellsGranularity11W3 | 0.689 |
| MedianCellsGranularity12W3 | 0.519 |
| MedianCytoplasmRadialDistributionRadialCVW23of5 | 0 |
| MedianCellsAreaShapeZernike31 | 0.031 |
| MedianCellsGranularity13W4 | 0.376 |
| MedianCellsGranularity11W4 | 0.573 |
| MedianCellsAreaShapeZernike44 | 0.02 |
| MedianCellsAreaShapeZernike66 | 0.011 |
| MedianCellsAreaShapeZernike64 | 0.014 |
| MedianCytoplasmCorrelationMandersW4W2 | 0.998 |
| MedianCellsAreaShapeZernike86 | 0.008 |
| MedianCellsAreaShapeZernike55 | 0.008 |
| MedianCellsAreaShapeZernike88 | 0.007 |
| MedianCellsAreaShapeOrientation | -0.699 |
| MedianCellsGranularity14W4 | 0.345 |
| MedianCellsAreaShapeZernike33 | 0.017 |
| MedianCellsAreaShapeZernike20 | 0.149 |
| MedianCytoplasmRadialDistributionMeanFracW45of5 | 1 |
| MedianCytoplasmCorrelationMandersW2W4 | 1 |
| MedianCellsAreaShapeZernike11 | 0.053 |
| MedianCellsAreaShapeZernike80 | 0.006 |
| MedianCellsGranularity14W3 | 0.367 |
| MedianCellsAreaShapeZernike97 | 0.006 |
| MedianCytoplasmCorrelationMandersW5W2 | 1 |

| | |
|---|---|
| MedianCellsGranularity13W3 | 0.425 |
| MedianCellsAreaShapeZernike51 | 0.015 |
| MedianCellsAreaShapeZernike95 | 0.006 |
| MedianCellsAreaShapeZernike82 | 0.007 |
| MedianCellsAreaShapeZernike53 | 0.015 |
| MedianCellsAreaShapeZernike75 | 0.009 |
| MedianCellsAreaShapeZernike84 | 0.007 |
| MedianCellsAreaShapeZernike71 | 0.009 |
| MedianCellsAreaShapeZernike73 | 0.009 |
| MedianCellsAreaShapeZernike93 | 0.006 |
| MedianCellsAreaShapeZernike60 | 0.01 |
| MedianCytoplasmCorrelationCostesW4W5 | 1 |
| MedianCellsAreaShapeZernike91 | 0.006 |
| MedianCellsAreaShapeZernike40 | 0.021 |
| MedianCellsGranularity14W1 | 0.081 |
| MedianCellsAreaShapeZernike62 | 0.012 |
| MedianNucleiCorrelationRWCW2W4 | 0.852 |
| MedianCellsNeighborsFirstClosestObjectNumber1 | 203.479 |
| MedianNucleiAreaShapeMajorAxisLength | 40.161 |
| MedianNucleiAreaShapeMaxFeretDiameter | 41.012 |
| MedianNucleiTextureDifferenceEntropyW1301256 | 2.336 |
| MedianNucleiTextureDifferenceEntropyW1302256 | 2.096 |
| MedianNucleiTextureInverseDifferenceMomentW1301256 | 0.43 |
| MedianNucleiTextureInverseDifferenceMomentW1302256 | 0.494 |
| MedianNucleiTextureDifferenceEntropyW3300256 | 2.055 |
| MedianNucleiTextureDifferenceEntropyW3302256 | 2.043 |
| MedianNucleiAreaShapeArea | 870.762 |
| MedianNucleiAreaShapeEquivalentDiameter | 33.101 |
| MedianNucleiTextureEntropyW3300256 | 5.768 |
| MedianNucleiTextureSumEntropyW3300256 | 4.171 |
| MedianNucleiTextureInverseDifferenceMomentW1300256 | 0.492 |
| MedianNucleiTextureInverseDifferenceMomentW1303256 | 0.43 |
| MedianNucleiTextureDifferenceEntropyW1303256 | 2.333 |
| MedianNucleiTextureDifferenceEntropyW1300256 | 2.099 |
| MedianNucleiRadialDistributionZernikeMagnitudeW544 | 0.002 |
| MedianNucleiRadialDistributionZernikeMagnitudeW588 | 0 |
| MedianNucleiRadialDistributionZernikeMagnitudeW577 | 0 |
| MedianNucleiRadialDistributionZernikeMagnitudeW599 | 0 |
| MedianNucleiIntensityMaxIntensityEdgeW1 | 0.056 |

| | |
|---|---|
| MedianNucleiIntensityMaxIntensityW1 | 0.076 |
| MedianNucleiTextureInverseDifferenceMomentW3300256 | 0.506 |
| MedianNucleiTextureInverseDifferenceMomentW3301256 | 0.437 |
| MedianNucleiTextureInverseDifferenceMomentW3303256 | 0.44 |
| MedianNucleiTextureEntropyW1300256 | 5.663 |
| MedianNucleiTextureSumEntropyW1300256 | 3.997 |
| MedianNucleiTextureInverseDifferenceMomentW3302256 | 0.508 |
| MedianNucleiCorrelationCostesW4W5 | 1 |
| MedianNucleiIntensityLowerQuartileIntensityW1 | 0.045 |
| MedianNucleiAreaShapeMaximumRadius | 13.125 |
| MedianNucleiAreaShapeMeanRadius | 5.08 |
| MedianNucleiRadialDistributionZernikeMagnitudeW560 | 0.001 |
| MedianNucleiRadialDistributionZernikeMagnitudeW586 | 0 |
| MedianNucleiRadialDistributionMeanFracW53of5 | 1.02 |
| MedianNucleiRadialDistributionMeanFracW55of5 | 0.978 |
| MedianNucleiGranularity16W4 | 0.286 |
| MedianNucleiTextureInfoMeas2W1300256 | 0.841 |
| MedianNucleiTextureInfoMeas2W1302256 | 0.842 |
| MedianNucleiRadialDistributionZernikeMagnitudeW593 | 0 |
| MedianNucleiRadialDistributionZernikeMagnitudeW595 | 0 |
| MedianNucleiRadialDistributionMeanFracW43of5 | 1.039 |
| MedianNucleiRadialDistributionMeanFracW45of5 | 0.951 |
| MedianNucleiIntensityMaxIntensityEdgeW3 | 0.092 |
| MedianNucleiIntensityMaxIntensityW3 | 0.106 |
| MedianNucleiIntensityLowerQuartileIntensityW3 | 0.065 |
| MedianNucleiRadialDistributionZernikeMagnitudeW282 | 0.001 |
| MedianNucleiRadialDistributionZernikeMagnitudeW286 | 0.001 |
| MedianNucleiGranularity15W4 | 0.281 |
| MedianNucleiTextureContrastW3300256 | 4.011 |
| MedianNucleiTextureSumVarianceW3300256 | 35.087 |
| MedianNucleiIntensityMinIntensityEdgeW1 | 0.027 |
| MedianNucleiRadialDistributionMeanFracW32of4 | 1.09 |
| MedianNucleiRadialDistributionMeanFracW34of4 | 0.935 |
| MedianNucleiRadialDistributionZernikeMagnitudeW264 | 0.001 |
| MedianNucleiRadialDistributionZernikeMagnitudeW573 | 0.001 |
| MedianNucleiTextureInfoMeas2W3300256 | 0.888 |
| MedianNucleiTextureInfoMeas2W3302256 | 0.891 |
| MedianNucleiGranularity16W3 | 0.286 |
| MedianNucleiRadialDistributionMeanFracW23of5 | 0.973 |

| | |
|---|---|
| MedianNucleiRadialDistributionMeanFracW25of5 | 1.034 |
| MedianNucleiRadialDistributionZernikeMagnitudeW260 | 0.001 |
| MedianNucleiRadialDistributionZernikeMagnitudeW540 | 0.002 |
| MedianNucleiCorrelationRWCW2W5 | 0.919 |
| MedianNucleiCorrelationRWCW5W2 | 0.922 |
| MedianNucleiAreaShapePerimeter | 120.217 |
| MedianNucleiRadialDistributionFracAtDW12of4 | 0.184 |
| MedianNucleiRadialDistributionFracAtDW14of4 | 0.414 |
| MedianNucleiRadialDistributionMeanFracW21of5 | 0.963 |
| MedianNucleiRadialDistributionMeanFracW22of5 | 0.968 |
| MedianNucleiGranularity15W3 | 0.289 |
| MedianNucleiRadialDistributionZernikeMagnitudeW553 | 0.001 |
| MedianNucleiRadialDistributionZernikeMagnitudeW575 | 0.001 |
| MedianNucleiTextureInfoMeas2W1301256 | 0.753 |
| MedianNucleiTextureInfoMeas2W1303256 | 0.754 |
| MedianNucleiTextureSumVarianceW1302256 | 24.834 |
| MedianNucleiTextureSumVarianceW1303256 | 22.673 |
| MedianNucleiIntensityMADIntensityW3 | 0.007 |
| MedianNucleiIntensityStdIntensityW3 | 0.011 |
| MedianNucleiGranularity7W1 | 4.349 |
| MedianNucleiRadialDistributionZernikeMagnitudeW531 | 0.002 |
| MedianNucleiRadialDistributionMeanFracW31of4 | 1.107 |
| MedianNucleiRadialDistributionZernikeMagnitudeW522 | 0.005 |
| MedianNucleiRadialDistributionZernikeMagnitudeW542 | 0.002 |
| MedianNucleiRadialDistributionZernikeMagnitudeW280 | 0.001 |
| MedianNucleiTextureInfoMeas2W3301256 | 0.826 |
| MedianNucleiTextureInfoMeas2W3303256 | 0.828 |
| MedianNucleiRadialDistributionZernikeMagnitudeW511 | 0.004 |
| MedianNucleiRadialDistributionZernikeMagnitudeW555 | 0.001 |
| MedianNucleiRadialDistributionMeanFracW41of5 | 1.043 |
| MedianNucleiRadialDistributionMeanFracW42of5 | 1.042 |
| MedianNucleiRadialDistributionZernikeMagnitudeW533 | 0.001 |
| MedianNucleiIntensityMinIntensityEdgeW3 | 0.044 |
| MedianNucleiRadialDistributionMeanFracW51of5 | 1.03 |
| MedianNucleiRadialDistributionMeanFracW52of5 | 1.028 |
| MedianNucleiRadialDistributionFracAtDW43of5 | 0.196 |
| MedianNucleiRadialDistributionFracAtDW45of5 | 0.365 |
| MedianNucleiRadialDistributionZernikeMagnitudeW551 | 0.001 |
| MedianNucleiRadialDistributionZernikeMagnitudeW571 | 0.001 |

| | |
|---|---|
| MedianNucleiRadialDistributionFracAtDW32of4 | 0.174 |
| MedianNucleiRadialDistributionFracAtDW34of4 | 0.442 |
| MedianNucleiAreaShapeMinFeretDiameter | 27.805 |
| MedianNucleiRadialDistributionMeanFracW11of4 | 1.179 |
| MedianNucleiRadialDistributionMeanFracW12of4 | 1.154 |
| MedianNucleiGranularity5W1 | 15.961 |
| MedianNucleiRadialDistributionZernikeMagnitudeW597 | 0 |
| MedianNucleiGranularity6W1 | 10.115 |
| MedianNucleiTextureSumVarianceW1300256 | 24.594 |
| MedianNucleiGranularity8W1 | 1.819 |
| MedianNucleiAreaShapeCompactness | 1.299 |
| MedianNucleiAreaShapeFormFactor | 0.772 |
| MedianNucleiGranularity7W3 | 2.927 |
| MedianNucleiRadialDistributionZernikeMagnitudeW591 | 0 |
| MedianNucleiGranularity1W3 | 21.124 |
| MedianNucleiGranularity1W1 | 11.012 |
| MedianNucleiRadialDistributionZernikeMagnitudeW284 | 0.001 |
| MedianNucleiRadialDistributionZernikeMagnitudeW520 | 0.012 |
| MedianNucleiRadialDistributionRadialCVW53of5 | 0.081 |
| MedianNucleiRadialDistributionRadialCVW54of5 | 0.124 |
| MedianNucleiGranularity7W4 | 2.472 |
| MedianNucleiGranularity4W1 | 12.614 |
| MedianNucleiRadialDistributionZernikeMagnitudeW244 | 0.002 |
| MedianNucleiRadialDistributionZernikeMagnitudeW266 | 0.001 |
| MedianNucleiRadialDistributionZernikeMagnitudeW288 | 0.001 |
| MedianNucleiGranularity1W4 | 29.299 |
| MedianNucleiRadialDistributionZernikeMagnitudeW500 | 0.044 |
| MedianNucleiRadialDistributionMeanFracW14of4 | 0.876 |
| MedianNucleiGranularity6W3 | 3.765 |
| MedianNucleiCorrelationCostesW2W4 | 1 |
| MedianNucleiCorrelationCostesW4W2 | 1 |
| MedianNucleiRadialDistributionZernikeMagnitudeW262 | 0.001 |
| MedianNucleiGranularity8W3 | 2.056 |
| MedianNucleiGranularity9W1 | 0.806 |
| MedianNucleiIntensityStdIntensityEdgeW3 | 0.011 |
| MedianNucleiTextureInfoMeas1W1300256 | -0.2 |
| MedianNucleiTextureInfoMeas1W1302256 | -0.201 |
| MedianNucleiGranularity6W4 | 3.332 |
| MedianNucleiGranularity10W1 | 0.386 |

| | |
|---|---|
| MedianNucleiGranularity16W1 | 0.031 |
| MedianNucleiCorrelationKW5W4 | 1.845 |
| MedianNucleiRadialDistributionRadialCVW24of5 | 0.132 |
| MedianNucleiRadialDistributionRadialCVW25of5 | 0.181 |
| MedianNucleiCorrelationKW2W5 | 0.681 |
| MedianNucleiRadialDistributionFracAtDW53of5 | 0.192 |
| MedianNucleiRadialDistributionFracAtDW55of5 | 0.376 |
| MedianNucleiTextureAngularSecondMomentW3300256 | 0.031 |
| MedianNucleiTextureDifferenceVarianceW3300256 | 0.01 |
| MedianNucleiRadialDistributionFracAtDW11of4 | 0.052 |
| MedianNucleiRadialDistributionFracAtDW31of4 | 0.049 |
| MedianNucleiRadialDistributionFracAtDW23of5 | 0.182 |
| MedianNucleiRadialDistributionFracAtDW25of5 | 0.4 |
| MedianNucleiRadialDistributionRadialCVW55of5 | 0.194 |
| MedianNucleiCorrelationKW5W2 | 1.611 |
| MedianNucleiGranularity5W3 | 4.145 |
| MedianNucleiTextureContrastW1301256 | 6.221 |
| MedianNucleiRadialDistributionZernikeMagnitudeW277 | 0.001 |
| MedianNucleiRadialDistributionZernikeMagnitudeW299 | 0 |
| MedianNucleiRadialDistributionZernikeMagnitudeW444 | 0.003 |
| MedianNucleiRadialDistributionZernikeMagnitudeW466 | 0.001 |
| MedianNucleiGranularity5W4 | 3.958 |
| MedianNucleiTextureInfoMeas1W1301256 | -0.137 |
| MedianNucleiTextureInfoMeas1W1303256 | -0.138 |
| MedianNucleiTextureCorrelationW3301256 | 0.615 |
| MedianNucleiTextureCorrelationW3302256 | 0.76 |
| MedianNucleiRadialDistributionZernikeMagnitudeW488 | 0.001 |
| MedianNucleiGranularity15W1 | 0.037 |
| MedianNucleiIntensityIntegratedIntensityEdgeW3 | 6.391 |
| MedianNucleiTextureCorrelationW3300256 | 0.755 |
| MedianNucleiRadialDistributionZernikeMagnitudeW240 | 0.003 |
| MedianNucleiTextureContrastW1302256 | 4.014 |
| MedianNucleiTextureContrastW1303256 | 6.01 |
| MedianNucleiTextureCorrelationW3303256 | 0.624 |
| MedianNucleiRadialDistributionZernikeMagnitudeW477 | 0.001 |
| MedianNucleiRadialDistributionZernikeMagnitudeW499 | 0 |
| MedianNucleiIntensityStdIntensityEdgeW1 | 0.007 |
| MedianNucleiIntensityIntegratedIntensityEdgeW1 | 3.837 |
| MedianNucleiGranularity4W4 | 4.034 |

| | |
|---|---|
| MedianNucleiTextureInfoMeas1W3301256 | -0.18 |
| MedianNucleiTextureInfoMeas1W3303256 | -0.182 |
| MedianNucleiGranularity11W1 | 0.194 |
| MedianNucleiRadialDistributionZernikeMagnitudeW222 | 0.007 |
| MedianNucleiRadialDistributionRadialCVW52of5 | 0.051 |
| MedianNucleiRadialDistributionRadialCVW23of5 | 0.098 |
| MedianNucleiRadialDistributionFracAtDW42of5 | 0.098 |
| MedianNucleiRadialDistributionZernikeMagnitudeW400 | 0.075 |
| MedianNucleiRadialDistributionZernikeMagnitudeW420 | 0.021 |
| MedianNucleiGranularity4W3 | 3.782 |
| MedianNucleiRadialDistributionZernikeMagnitudeW273 | 0.001 |
| MedianNucleiRadialDistributionZernikeMagnitudeW275 | 0.001 |
| MedianNucleiRadialDistributionZernikeMagnitudeW475 | 0.001 |
| MedianNucleiRadialDistributionZernikeMagnitudeW497 | 0.001 |
| MedianNucleiTextureCorrelationW1300256 | 0.673 |
| MedianNucleiTextureCorrelationW1302256 | 0.675 |
| MedianNucleiIntensityStdIntensityW1 | 0.01 |
| MedianNucleiRadialDistributionFracAtDW13of4 | 0.348 |
| MedianNucleiRadialDistributionZernikeMagnitudeW482 | 0.001 |
| MedianNucleiRadialDistributionZernikeMagnitudeW486 | 0.001 |
| MedianNucleiRadialDistributionZernikeMagnitudeW464 | 0.001 |
| MedianNucleiRadialDistributionZernikeMagnitudeW200 | 0.066 |
| MedianNucleiRadialDistributionZernikeMagnitudeW220 | 0.016 |
| MedianNucleiRadialDistributionZernikeMagnitudeW455 | 0.001 |
| MedianNucleiRadialDistributionZernikeMagnitudeW271 | 0.001 |
| MedianNucleiRadialDistributionZernikeMagnitudeW233 | 0.002 |
| MedianNucleiRadialDistributionZernikeMagnitudeW255 | 0.001 |
| MedianNucleiRadialDistributionZernikeMagnitudeW422 | 0.008 |
| MedianNucleiRadialDistributionZernikeMagnitudeW291 | 0.001 |
| MedianNucleiRadialDistributionZernikeMagnitudeW293 | 0.001 |
| MedianNucleiRadialDistributionZernikeMagnitudeW297 | 0.001 |
| MedianNucleiRadialDistributionZernikeMagnitudeW295 | 0.001 |
| MedianNucleiRadialDistributionZernikeMagnitudeW495 | 0.001 |
| MedianNucleiTextureInfoMeas1W3300256 | -0.243 |
| MedianNucleiTextureInfoMeas1W3302256 | -0.246 |
| MedianNucleiRadialDistributionZernikeMagnitudeW493 | 0.001 |
| MedianNucleiTextureCorrelationW1301256 | 0.512 |
| MedianNucleiTextureCorrelationW1303256 | 0.514 |
| MedianNucleiRadialDistributionZernikeMagnitudeW253 | 0.001 |

| | |
|---|---|
| MedianNucleiGranularity3W1 | 6.938 |
| MedianNucleiRadialDistributionZernikeMagnitudeW473 | 0.001 |
| MedianNucleiRadialDistributionZernikeMagnitudeW471 | 0.001 |
| MedianNucleiRadialDistributionZernikeMagnitudeW491 | 0.001 |
| MedianNucleiRadialDistributionZernikeMagnitudeW433 | 0.002 |
| MedianNucleiRadialDistributionZernikeMagnitudeW484 | 0.001 |
| MedianNucleiRadialDistributionZernikeMagnitudeW242 | 0.002 |
| MedianNucleiRadialDistributionZernikeMagnitudeW411 | 0.007 |
| MedianNucleiRadialDistributionZernikeMagnitudeW451 | 0.002 |
| MedianNucleiRadialDistributionZernikeMagnitudeW453 | 0.001 |
| MedianNucleiRadialDistributionFracAtDW33of4 | 0.334 |
| MedianNucleiRadialDistributionZernikeMagnitudeW211 | 0.006 |
| MedianNucleiRadialDistributionZernikeMagnitudeW231 | 0.002 |
| MedianNucleiGranularity12W4 | 0.388 |
| MedianNucleiCorrelationCostesW2W5 | 1 |
| MedianNucleiCorrelationCostesW5W2 | 1 |
| MedianNucleiRadialDistributionFracAtDW52of5 | 0.097 |
| MedianNucleiRadialDistributionZernikeMagnitudeW251 | 0.002 |
| MedianNucleiRadialDistributionMeanFracW33of4 | 1.034 |
| MedianNucleiAreaShapeMedianRadius | 4.539 |
| MedianNucleiRadialDistributionRadialCVW22of5 | 0.068 |
| MedianNucleiGranularity3W4 | 3.327 |
| MedianNucleiRadialDistributionZernikeMagnitudeW462 | 0.001 |
| MedianNucleiRadialDistributionZernikeMagnitudeW480 | 0.001 |
| MedianNucleiRadialDistributionRadialCVW43of5 | 0.071 |
| MedianNucleiRadialDistributionRadialCVW44of5 | 0.085 |
| MedianNucleiRadialDistributionRadialCVW31of4 | 0.038 |
| MedianNucleiRadialDistributionRadialCVW32of4 | 0.067 |
| MedianNucleiRadialDistributionRadialCVW42of5 | 0.058 |
| MedianNucleiCorrelationKW4W5 | 0.656 |
| MedianNucleiGranularity2W1 | 3.479 |
| MedianNucleiIntensityIntegratedIntensityW3 | 66.591 |
| MedianNucleiRadialDistributionZernikeMagnitudeW431 | 0.002 |
| MedianNucleiGranularity8W4 | 1.665 |
| MedianNucleiRadialDistributionZernikeMagnitudeW442 | 0.003 |
| MedianNucleiRadialDistributionZernikeMagnitudeW460 | 0.001 |
| MedianNucleiRadialDistributionRadialCVW45of5 | 0.122 |
| MedianNucleiCorrelationKW2W4 | 1.153 |
| MedianNucleiCorrelationKW4W2 | 1.06 |

| | |
|---|---|
| MedianNucleiRadialDistributionFracAtDW21of5 | 0.027 |
| MedianNucleiGranularity3W3 | 2.898 |
| MedianNucleiAreaShapeExtent | 0.668 |
| MedianNucleiAreaShapeZernike00 | 0.657 |
| MedianNucleiRadialDistributionRadialCVW21of5 | 0.038 |
| MedianNucleiRadialDistributionFracAtDW22of5 | 0.091 |
| MedianNucleiRadialDistributionRadialCVW12of4 | 0.069 |
| MedianNucleiRadialDistributionRadialCVW13of4 | 0.087 |
| MedianNucleiRadialDistributionFracAtDW41of5 | 0.029 |
| MedianNucleiRadialDistributionFracAtDW51of5 | 0.029 |
| MedianNucleiRadialDistributionRadialCVW41of5 | 0.036 |
| MedianNucleiRadialDistributionRadialCVW33of4 | 0.089 |
| MedianNucleiGranularity2W4 | 2.108 |
| MedianNucleiTextureAngularSecondMomentW1300256 | 0.031 |
| MedianNucleiTextureDifferenceVarianceW1300256 | 0.012 |
| MedianNucleiRadialDistributionRadialCVW34of4 | 0.125 |
| MedianNucleiRadialDistributionMeanFracW13of4 | 1.081 |
| MedianNucleiRadialDistributionRadialCVW11of4 | 0.045 |
| MedianNucleiGranularity12W1 | 0.107 |
| MedianNucleiGranularity13W1 | 0.066 |
| MedianNucleiRadialDistributionMeanFracW24of5 | 0.986 |
| MedianNucleiRadialDistributionZernikeMagnitudeW440 | 0.003 |
| MedianNucleiChildrenCellsCount | 1 |
| MedianNucleiIntensityIntegratedIntensityW1 | 50.964 |
| MedianNucleiAreaShapeSolidity | 0.942 |
| MedianNucleiGranularity2W3 | 2.035 |
| MedianNucleiIntensityMassDisplacementW1 | 0.54 |
| MedianNucleiRadialDistributionRadialCVW14of4 | 0.114 |
| MedianNucleiIntensityMassDisplacementW3 | 0.63 |
| MedianNucleiGranularity10W3 | 0.916 |
| MedianNucleiCorrelationMandersW2W5 | 1 |
| MedianNucleiIntensityMADIntensityW1 | 0.007 |
| MedianNucleiTextureContrastW1300256 | 4.044 |
| MedianNucleiCorrelationCorrelationW1W3 | 0.592 |
| MedianNucleiAreaShapeZernike42 | 0.02 |
| MedianNucleiAreaShapeZernike77 | 0.005 |
| MedianNucleiAreaShapeZernike99 | 0.004 |
| MedianNucleiRadialDistributionRadialCVW51of5 | 0.03 |
| MedianNucleiCorrelationKW3W1 | 0.791 |

| | |
|---|---|
| MedianNucleiGranularity10W4 | 0.715 |
| MedianNucleiRadialDistributionMeanFracW44of5 | 1.02 |
| MedianNucleiCorrelationKW1W3 | 1.495 |
| MedianNucleiCorrelationCorrelationW4W5 | 0.569 |
| MedianNucleiAreaShapeZernike66 | 0.011 |
| MedianNucleiAreaShapeZernike88 | 0.006 |
| MedianNucleiRadialDistributionFracAtDW24of5 | 0.298 |
| MedianNucleiRadialDistributionFracAtDW54of5 | 0.303 |
| MedianNucleiAreaShapeEccentricity | 0.698 |
| MedianNucleiGranularity14W1 | 0.049 |
| MedianNucleiCorrelationOverlapW4W5 | 0.989 |
| MedianNucleiRadialDistributionFracAtDW44of5 | 0.31 |
| MedianNucleiCorrelationCorrelationW2W5 | 0.624 |
| MedianNucleiGranularity9W3 | 1.355 |
| MedianNucleiGranularity9W4 | 1.065 |
| MedianNucleiCorrelationCorrelationW2W4 | 0.264 |
| MedianNucleiCorrelationOverlapW2W4 | 0.978 |
| MedianNucleiCorrelationRWCW4W5 | 0.864 |
| MedianNucleiCorrelationOverlapW2W5 | 0.987 |
| MedianNucleiAreaShapeZernike97 | 0.005 |
| MedianNucleiAreaShapeZernike55 | 0.009 |
| MedianNucleiAreaShapeZernike22 | 0.071 |
| MedianNucleiAreaShapeZernike44 | 0.024 |
| MedianNucleiRadialDistributionMeanFracW54of5 | 1.002 |
| MedianNucleiAreaShapeZernike11 | 0.056 |
| MedianNucleiAreaShapeZernike51 | 0.014 |
| MedianNucleiAreaShapeZernike64 | 0.009 |
| MedianNucleiAreaShapeZernike40 | 0.024 |
| MedianNucleiAreaShapeZernike33 | 0.018 |
| MedianNucleiAreaShapeZernike31 | 0.017 |
| MedianNucleiGranularity13W4 | 0.331 |
| MedianNucleiGranularity14W4 | 0.305 |
| MedianNucleiCorrelationRWCW1W3 | 0.903 |
| MedianNucleiGranularity14W3 | 0.322 |
| MedianNucleiGranularity11W3 | 0.632 |
| MedianNucleiAreaShapeZernike75 | 0.007 |
| MedianNucleiAreaShapeZernike86 | 0.006 |
| MedianNucleiAreaShapeZernike62 | 0.013 |
| MedianNucleiAreaShapeZernike20 | 0.16 |

| | |
|---|---|
| MedianNucleiAreaShapeOrientation | -0.904 |
| MedianNucleiAreaShapeZernike71 | 0.008 |
| MedianNucleiGranularity13W3 | 0.381 |
| MedianNucleiAreaShapeZernike80 | 0.006 |
| MedianNucleiAreaShapeZernike82 | 0.007 |
| MedianNucleiCorrelationOverlapW1W3 | 0.986 |
| MedianNucleiAreaShapeZernike53 | 0.012 |
| MedianNucleiAreaShapeZernike95 | 0.005 |
| MedianNucleiAreaShapeZernike73 | 0.008 |
| MedianNucleiGranularity11W4 | 0.511 |
| MedianNucleiAreaShapeZernike91 | 0.005 |
| MedianNucleiAreaShapeZernike93 | 0.005 |
| MedianNucleiCorrelationCostesW1W3 | 0.999 |
| MedianNucleiAreaShapeZernike84 | 0.007 |
| MedianNucleiCorrelationCostesW3W1 | 0.997 |
| MedianNucleiAreaShapeZernike60 | 0.01 |
| MedianNucleiGranularity12W3 | 0.47 |
| MedianNucleiLocationCenterMassIntensityYW1 | 1032.91 |
| MedianNucleiLocationCenterMassIntensityXW1 | 1022.556 |
| Granularity1W3 | 2.917 |
| Granularity1W4 | 3.967 |
| Granularity5W3 | 3.963 |
| Granularity6W3 | 4.19 |
| Granularity2W3 | 3.027 |
| Granularity2W4 | 3.908 |
| Granularity3W4 | 3.947 |
| Granularity5W4 | 4.108 |
| Granularity6W4 | 4.12 |
| Granularity4W3 | 3.511 |
| Granularity3W3 | 3.217 |
| Granularity7W3 | 4.036 |
| Granularity4W4 | 3.977 |
| Granularity7W4 | 3.894 |
| Granularity5W1 | 17.779 |
| Granularity8W3 | 3.652 |
| Granularity1W1 | 3.103 |
| Granularity4W1 | 13.669 |
| Granularity8W4 | 3.51 |
| Granularity6W1 | 15.28 |

| | |
|---|---|
| Granularity9W3 | 3.234 |
| Granularity2W1 | 5.331 |
| Granularity3W1 | 9.173 |
| Granularity9W4 | 3.091 |
| Granularity7W1 | 9.484 |
| Granularity10W1 | 1.972 |
| Granularity11W1 | 1.363 |
| Granularity12W1 | 1.018 |
| Granularity10W3 | 2.87 |
| Granularity8W1 | 5.265 |
| Granularity9W1 | 3.072 |
| Granularity13W1 | 0.784 |
| Granularity14W1 | 0.617 |
| Granularity10W4 | 2.715 |
| Granularity16W3 | 1.737 |
| Granularity15W3 | 1.823 |
| Granularity15W4 | 1.655 |
| Granularity12W3 | 2.313 |
| Granularity12W4 | 2.128 |
| Granularity13W4 | 1.93 |
| Granularity15W1 | 0.519 |
| Granularity16W1 | 0.445 |
| Granularity14W4 | 1.775 |
| Granularity16W4 | 1.569 |
| Granularity14W3 | 1.954 |
| Granularity11W3 | 2.566 |
| Granularity11W4 | 2.401 |
| Granularity13W3 | 2.113 |
| ImageQualityStdIntensityW2 | 0.029 |

**Supplementary Information Table 3 | Table summarising statistical accuracy, sensitivity, specificity and precision of Neural Network classification.**

| |
|---|
| Confusion Matrix and Statistics |
| Reference |
| Prediction NEGATIVE POSITIVE |
| NEGATIVE 1439 0 |
| POSITIVE 0 1564 |
| Accuracy : 1 |
| 95% CI : (0.9988, 1) |
| No Information Rate : 0.5208 |
| P-Value [Acc > NIR] : < 2.2e-16 |
| Kappa : 1 |
| Mcnemar's Test P-Value : NA |
| Sensitivity : 1.0000 |
| Specificity : 1.0000 |
| Pos Pred Value : 1.0000 |
| Neg Pred Value : 1.0000 |
| Prevalence : 0.4792 |
| Detection Rate : 0.4792 |
| Detection Prevalence : 0.4792 |
| Balanced Accuracy : 1.0000 |
| 'Positive' Class : NEGATIVE |

**Supplementary Information Table 4 | Table of classification scores (probPOSITIVE and probNEGATIVE) for untreated (NEGATIVE2), DMSO-vehicle treated (NEGATIVE) SORL1$^{-/-}$ NPCs and DMSO-vehicle treated wild-type controls (POSITIVE) classes.**

| ReagentClass | ReagentID | probPOSITIVE | probNEGATIVE | pvalue |
|---|---|---|---|---|
| NEGATIVE | DMSO | 0 | 1 | 0.58194103 |
| NEGATIVE | DMSO | 0 | 1 | 0.58194103 |
| NEGATIVE | DMSO | 0 | 1 | 0.58194103 |
| NEGATIVE | DMSO | 0 | 1 | 0.58194103 |
| NEGATIVE | DMSO | 0 | 1 | 0.58194103 |
| NEGATIVE | DMSO | 0 | 1 | 0.58194103 |
| NEGATIVE | DMSO | 0 | 1 | 0.58194103 |
| NEGATIVE | DMSO | 0 | 1 | 0.58194103 |
| NEGATIVE | DMSO | 0 | 1 | 0.58194103 |
| NEGATIVE | DMSO | 0 | 1 | 0.58194103 |
| NEGATIVE | DMSO | 0 | 1 | 0.58194103 |
| NEGATIVE | DMSO | 0 | 1 | 0.58194103 |
| NEGATIVE | DMSO | 0 | 1 | 0.58194103 |
| NEGATIVE | DMSO | 0 | 1 | 0.58194103 |
| NEGATIVE | DMSO | 0 | 1 | 0.58194103 |
| NEGATIVE | DMSO | 0 | 1 | 0.58194103 |
| NEGATIVE | DMSO | 0 | 1 | 0.58194103 |
| NEGATIVE | DMSO | 0 | 1 | 0.58194103 |
| NEGATIVE | DMSO | 0 | 1 | 0.58194103 |
| NEGATIVE | DMSO | 0 | 1 | 0.58194103 |
| NEGATIVE | DMSO | 0 | 1 | 0.58194103 |
| NEGATIVE | DMSO | 0 | 1 | 0.58194103 |
| NEGATIVE | DMSO | 0 | 1 | 0.58194103 |
| NEGATIVE | DMSO | 0 | 1 | 0.58194103 |
| NEGATIVE | DMSO | 0 | 1 | 0.58194103 |
| NEGATIVE | DMSO | 0 | 1 | 0.58194103 |
| NEGATIVE | DMSO | 0 | 1 | 0.58194103 |
| NEGATIVE | DMSO | 0 | 1 | 0.58194103 |
| NEGATIVE | DMSO | 0 | 1 | 0.58194103 |
| NEGATIVE | DMSO | 0 | 1 | 0.58194103 |
| NEGATIVE | DMSO | 0 | 1 | 0.58194103 |
| NEGATIVE | DMSO | 0 | 1 | 0.58194103 |
| NEGATIVE | DMSO | 0 | 1 | 0.58194103 |
| NEGATIVE | DMSO | 0 | 1 | 0.58194103 |
| NEGATIVE | DMSO | 0 | 1 | 0.58194103 |

| | | | | |
|---|---|---|---|---|
| NEGATIVE | DMSO | 0 | 1 | 0.58194103 |
| NEGATIVE | DMSO | 0 | 1 | 0.58194103 |
| NEGATIVE | DMSO | 8.22E-08 | 0.99999992 | 0.58194103 |
| NEGATIVE | DMSO | 8.04E-08 | 0.99999992 | 0.58194103 |
| NEGATIVE | DMSO | 7.69E-08 | 0.99999992 | 0.58194103 |
| NEGATIVE | DMSO | 1.02E-07 | 0.9999999 | 0.58194103 |
| NEGATIVE | DMSO | 1.20E-07 | 0.99999988 | 0.58194103 |
| NEGATIVE | DMSO | 1.17E-07 | 0.99999988 | 0.58194103 |
| NEGATIVE | DMSO | 1.15E-07 | 0.99999988 | 0.58194103 |
| NEGATIVE | DMSO | 1.26E-07 | 0.99999987 | 0.58194103 |
| NEGATIVE | DMSO | 1.79E-07 | 0.99999982 | 0.58194103 |
| NEGATIVE | DMSO | 1.91E-07 | 0.99999981 | 0.58194103 |
| NEGATIVE | DMSO | 2.14E-07 | 0.99999979 | 0.58194103 |
| NEGATIVE | DMSO | 2.29E-07 | 0.99999977 | 0.58194103 |
| NEGATIVE | DMSO | 2.88E-07 | 0.99999971 | 0.58194103 |
| NEGATIVE | DMSO | 3.14E-07 | 0.99999969 | 0.58194103 |
| NEGATIVE | DMSO | 3.24E-07 | 0.99999968 | 0.58194103 |
| NEGATIVE | DMSO | 3.25E-07 | 0.99999967 | 0.58194103 |
| NEGATIVE | DMSO | 3.34E-07 | 0.99999967 | 0.58194103 |
| NEGATIVE | DMSO | 4.33E-07 | 0.99999957 | 0.58194103 |
| NEGATIVE | DMSO | 4.50E-07 | 0.99999955 | 0.58194103 |
| NEGATIVE | DMSO | 4.73E-07 | 0.99999953 | 0.58194103 |
| NEGATIVE | DMSO | 4.94E-07 | 0.99999951 | 0.58194103 |
| NEGATIVE | DMSO | 5.11E-07 | 0.99999949 | 0.58194103 |
| NEGATIVE | DMSO | 5.56E-07 | 0.99999944 | 0.58194103 |
| NEGATIVE | DMSO | 6.24E-07 | 0.99999938 | 0.58194103 |
| NEGATIVE | DMSO | 6.56E-07 | 0.99999934 | 0.58194103 |
| NEGATIVE | DMSO | 6.69E-07 | 0.99999933 | 0.58194103 |
| NEGATIVE | DMSO | 7.93E-07 | 0.99999921 | 0.58194103 |
| NEGATIVE | DMSO | 8.50E-07 | 0.99999915 | 0.58194103 |
| NEGATIVE | DMSO | 8.64E-07 | 0.99999914 | 0.58194103 |
| NEGATIVE | DMSO | 8.75E-07 | 0.99999913 | 0.58194103 |
| NEGATIVE | DMSO | 9.54E-07 | 0.99999905 | 0.58194103 |
| NEGATIVE | DMSO | 1.25E-06 | 0.99999875 | 0.58194103 |
| NEGATIVE | DMSO | 1.32E-06 | 0.99999868 | 0.58194103 |
| NEGATIVE | DMSO | 1.52E-06 | 0.99999848 | 0.58194103 |
| NEGATIVE | DMSO | 1.53E-06 | 0.99999847 | 0.58194103 |
| NEGATIVE | DMSO | 1.68E-06 | 0.99999832 | 0.58194103 |
| NEGATIVE | DMSO | 1.70E-06 | 0.9999983 | 0.58194103 |

| | | | | |
|---|---|---|---|---|
| NEGATIVE | DMSO | 1.76E-06 | 0.99999824 | 0.58194103 |
| NEGATIVE | DMSO | 1.77E-06 | 0.99999823 | 0.58194103 |
| NEGATIVE | DMSO | 2.08E-06 | 0.99999792 | 0.58194103 |
| NEGATIVE | DMSO | 2.26E-06 | 0.99999774 | 0.58194103 |
| NEGATIVE | DMSO | 2.55E-06 | 0.99999745 | 0.58194103 |
| NEGATIVE | DMSO | 2.79E-06 | 0.99999721 | 0.58194103 |
| NEGATIVE | DMSO | 3.30E-06 | 0.9999967 | 0.58194103 |
| NEGATIVE | DMSO | 3.60E-06 | 0.9999964 | 0.58194103 |
| NEGATIVE | DMSO | 3.74E-06 | 0.99999626 | 0.58194103 |
| NEGATIVE | DMSO | 4.37E-06 | 0.99999563 | 0.58194103 |
| NEGATIVE | DMSO | 4.44E-06 | 0.99999556 | 0.58194103 |
| NEGATIVE | DMSO | 5.19E-06 | 0.99999481 | 0.58194103 |
| NEGATIVE | DMSO | 5.86E-06 | 0.99999414 | 0.58194103 |
| NEGATIVE | DMSO | 6.88E-06 | 0.99999312 | 0.58194103 |
| NEGATIVE | DMSO | 7.54E-06 | 0.99999246 | 0.58194103 |
| NEGATIVE | DMSO | 7.68E-06 | 0.99999232 | 0.58194103 |
| NEGATIVE | DMSO | 8.06E-06 | 0.99999194 | 0.58194103 |
| NEGATIVE | DMSO | 9.05E-06 | 0.99999095 | 0.58194103 |
| NEGATIVE | DMSO | 9.22E-06 | 0.99999078 | 0.58194103 |
| NEGATIVE | DMSO | 9.68E-06 | 0.99999032 | 0.58194103 |
| NEGATIVE | DMSO | 1.02E-05 | 0.9999898 | 0.58194103 |
| NEGATIVE | DMSO | 1.23E-05 | 0.99998765 | 0.58194103 |
| NEGATIVE | DMSO | 1.26E-05 | 0.99998738 | 0.58194103 |
| NEGATIVE | DMSO | 1.41E-05 | 0.9999859 | 0.58194103 |
| NEGATIVE | DMSO | 1.46E-05 | 0.99998537 | 0.58194103 |
| NEGATIVE | DMSO | 1.50E-05 | 0.99998496 | 0.58194103 |
| NEGATIVE | DMSO | 1.71E-05 | 0.99998294 | 0.58194103 |
| NEGATIVE | DMSO | 1.72E-05 | 0.99998279 | 0.58194103 |
| NEGATIVE | DMSO | 1.81E-05 | 0.9999819 | 0.58194103 |
| NEGATIVE | DMSO | 1.91E-05 | 0.99998093 | 0.58194103 |
| NEGATIVE | DMSO | 2.04E-05 | 0.99997956 | 0.58194103 |
| NEGATIVE | DMSO | 2.35E-05 | 0.99997654 | 0.58194103 |
| NEGATIVE | DMSO | 2.76E-05 | 0.99997237 | 0.58194103 |
| NEGATIVE | DMSO | 2.92E-05 | 0.99997082 | 0.58194103 |
| NEGATIVE | DMSO | 3.11E-05 | 0.9999689 | 0.58194103 |
| NEGATIVE | DMSO | 3.12E-05 | 0.9999688 | 0.58194103 |
| NEGATIVE | DMSO | 3.20E-05 | 0.99996802 | 0.58194103 |
| NEGATIVE | DMSO | 3.20E-05 | 0.99996799 | 0.58194103 |
| NEGATIVE | DMSO | 3.22E-05 | 0.99996784 | 0.58194103 |

| | | | | |
|---|---|---|---|---|
| NEGATIVE | DMSO | 3.32E-05 | 0.9999668 | 0.58194103 |
| NEGATIVE | DMSO | 4.20E-05 | 0.99995803 | 0.58194103 |
| NEGATIVE | DMSO | 4.55E-05 | 0.99995446 | 0.58194103 |
| NEGATIVE | DMSO | 4.67E-05 | 0.99995334 | 0.58194103 |
| NEGATIVE | DMSO | 5.19E-05 | 0.99994815 | 0.58194103 |
| NEGATIVE | DMSO | 5.73E-05 | 0.99994271 | 0.58194103 |
| NEGATIVE | DMSO | 6.25E-05 | 0.99993751 | 0.58194103 |
| NEGATIVE | DMSO | 6.35E-05 | 0.99993652 | 0.58194103 |
| NEGATIVE | DMSO | 7.28E-05 | 0.99992723 | 0.58194103 |
| NEGATIVE | DMSO | 7.76E-05 | 0.99992238 | 0.58194103 |
| NEGATIVE | DMSO | 8.52E-05 | 0.99991484 | 0.58194103 |
| NEGATIVE | DMSO | 9.61E-05 | 0.99990391 | 0.58194103 |
| NEGATIVE | DMSO | 0.00011278 | 0.99988722 | 0.58194103 |
| NEGATIVE | DMSO | 0.00011643 | 0.99988357 | 0.58194103 |
| NEGATIVE | DMSO | 0.0001255 | 0.9998745 | 0.58194103 |
| NEGATIVE | DMSO | 0.000129 | 0.999871 | 0.58194103 |
| NEGATIVE | DMSO | 0.00013832 | 0.99986168 | 0.58194103 |
| NEGATIVE | DMSO | 0.0001454 | 0.9998546 | 0.58194103 |
| NEGATIVE | DMSO | 0.0001462 | 0.9998538 | 0.58194103 |
| NEGATIVE | DMSO | 0.00015305 | 0.99984695 | 0.58194103 |
| NEGATIVE | DMSO | 0.0002115 | 0.9997885 | 0.58194103 |
| NEGATIVE | DMSO | 0.00021473 | 0.99978527 | 0.58194103 |
| NEGATIVE | DMSO | 0.00021586 | 0.99978414 | 0.58194103 |
| NEGATIVE | DMSO | 0.00022974 | 0.99977026 | 0.58194103 |
| NEGATIVE | DMSO | 0.00029811 | 0.99970189 | 0.47075057 |
| NEGATIVE | DMSO | 0.00043339 | 0.99956661 | 0.26547938 |
| NEGATIVE | DMSO | 0.00091896 | 0.99908104 | 0.00970182 |
| NEGATIVE | DMSO | 0.00113924 | 0.99886076 | 0.00111048 |
| NEGATIVE | DMSO | 0.00357971 | 0.99642029 | 5.49E-27 |
| POSITIVE | DMSO | 0.99685605 | 0.00314395 | 0 |
| POSITIVE | DMSO | 0.9984143 | 0.0015857 | 0 |
| POSITIVE | DMSO | 0.99909456 | 0.00090544 | 0 |
| POSITIVE | DMSO | 0.99917818 | 0.00082182 | 0 |
| POSITIVE | DMSO | 0.99942156 | 0.00057844 | 0 |
| POSITIVE | DMSO | 0.99948062 | 0.00051938 | 0 |
| POSITIVE | DMSO | 0.99968201 | 0.00031799 | 0 |
| POSITIVE | DMSO | 0.99970265 | 0.00029735 | 0 |
| POSITIVE | DMSO | 0.99982333 | 0.00017667 | 0 |
| POSITIVE | DMSO | 0.99987839 | 0.00012161 | 0 |

| | | | | |
|---|---|---|---|---|
| POSITIVE | DMSO | 0.99989424 | 0.00010576 | 0 |
| POSITIVE | DMSO | 0.99990674 | 9.33E-05 | 0 |
| POSITIVE | DMSO | 0.99992721 | 7.28E-05 | 0 |
| POSITIVE | DMSO | 0.99992922 | 7.08E-05 | 0 |
| POSITIVE | DMSO | 0.99993056 | 6.94E-05 | 0 |
| POSITIVE | DMSO | 0.99994738 | 5.26E-05 | 0 |
| POSITIVE | DMSO | 0.99995368 | 4.63E-05 | 0 |
| POSITIVE | DMSO | 0.99996442 | 3.56E-05 | 0 |
| POSITIVE | DMSO | 0.99997305 | 2.69E-05 | 0 |
| POSITIVE | DMSO | 0.99998077 | 1.92E-05 | 0 |
| POSITIVE | DMSO | 0.9999841 | 1.59E-05 | 0 |
| POSITIVE | DMSO | 0.99998408 | 1.59E-05 | 0 |
| POSITIVE | DMSO | 0.99998711 | 1.29E-05 | 0 |
| POSITIVE | DMSO | 0.9999875 | 1.25E-05 | 0 |
| POSITIVE | DMSO | 0.99998832 | 1.17E-05 | 0 |
| POSITIVE | DMSO | 0.99998923 | 1.08E-05 | 0 |
| POSITIVE | DMSO | 0.99999248 | 7.52E-06 | 0 |
| POSITIVE | DMSO | 0.99999345 | 6.55E-06 | 0 |
| POSITIVE | DMSO | 0.99999345 | 6.55E-06 | 0 |
| POSITIVE | DMSO | 0.99999396 | 6.04E-06 | 0 |
| POSITIVE | DMSO | 0.99999549 | 4.51E-06 | 0 |
| POSITIVE | DMSO | 0.99999557 | 4.43E-06 | 0 |
| POSITIVE | DMSO | 0.9999964 | 3.60E-06 | 0 |
| POSITIVE | DMSO | 0.99999649 | 3.51E-06 | 0 |
| POSITIVE | DMSO | 0.99999674 | 3.26E-06 | 0 |
| POSITIVE | DMSO | 0.99999682 | 3.18E-06 | 0 |
| POSITIVE | DMSO | 0.99999709 | 2.91E-06 | 0 |
| POSITIVE | DMSO | 0.99999738 | 2.62E-06 | 0 |
| POSITIVE | DMSO | 0.99999747 | 2.53E-06 | 0 |
| POSITIVE | DMSO | 0.99999755 | 2.45E-06 | 0 |
| POSITIVE | DMSO | 0.9999976 | 2.40E-06 | 0 |
| POSITIVE | DMSO | 0.99999777 | 2.23E-06 | 0 |
| POSITIVE | DMSO | 0.99999786 | 2.14E-06 | 0 |
| POSITIVE | DMSO | 0.99999793 | 2.07E-06 | 0 |
| POSITIVE | DMSO | 0.99999796 | 2.04E-06 | 0 |
| POSITIVE | DMSO | 0.99999817 | 1.83E-06 | 0 |
| POSITIVE | DMSO | 0.99999831 | 1.69E-06 | 0 |
| POSITIVE | DMSO | 0.99999845 | 1.55E-06 | 0 |
| POSITIVE | DMSO | 0.99999853 | 1.47E-06 | 0 |

| | | | | |
|---|---|---|---|---|
| POSITIVE | DMSO | 0.99999855 | 1.45E-06 | 0 |
| POSITIVE | DMSO | 0.99999858 | 1.42E-06 | 0 |
| POSITIVE | DMSO | 0.99999869 | 1.31E-06 | 0 |
| POSITIVE | DMSO | 0.99999875 | 1.25E-06 | 0 |
| POSITIVE | DMSO | 0.99999886 | 1.14E-06 | 0 |
| POSITIVE | DMSO | 0.99999886 | 1.14E-06 | 0 |
| POSITIVE | DMSO | 0.99999887 | 1.13E-06 | 0 |
| POSITIVE | DMSO | 0.99999888 | 1.12E-06 | 0 |
| POSITIVE | DMSO | 0.99999888 | 1.12E-06 | 0 |
| POSITIVE | DMSO | 0.99999892 | 1.08E-06 | 0 |
| POSITIVE | DMSO | 0.99999896 | 1.04E-06 | 0 |
| POSITIVE | DMSO | 0.99999902 | 9.79E-07 | 0 |
| POSITIVE | DMSO | 0.99999908 | 9.19E-07 | 0 |
| POSITIVE | DMSO | 0.99999912 | 8.81E-07 | 0 |
| POSITIVE | DMSO | 0.99999915 | 8.47E-07 | 0 |
| POSITIVE | DMSO | 0.9999992 | 8.00E-07 | 0 |
| POSITIVE | DMSO | 0.9999993 | 7.05E-07 | 0 |
| POSITIVE | DMSO | 0.99999935 | 6.46E-07 | 0 |
| POSITIVE | DMSO | 0.99999942 | 5.82E-07 | 0 |
| POSITIVE | DMSO | 0.99999944 | 5.64E-07 | 0 |
| POSITIVE | DMSO | 0.99999945 | 5.46E-07 | 0 |
| POSITIVE | DMSO | 0.99999947 | 5.31E-07 | 0 |
| POSITIVE | DMSO | 0.99999947 | 5.27E-07 | 0 |
| POSITIVE | DMSO | 0.99999948 | 5.19E-07 | 0 |
| POSITIVE | DMSO | 0.99999951 | 4.93E-07 | 0 |
| POSITIVE | DMSO | 0.99999951 | 4.85E-07 | 0 |
| POSITIVE | DMSO | 0.99999952 | 4.83E-07 | 0 |
| POSITIVE | DMSO | 0.99999952 | 4.80E-07 | 0 |
| POSITIVE | DMSO | 0.99999952 | 4.79E-07 | 0 |
| POSITIVE | DMSO | 0.99999952 | 4.78E-07 | 0 |
| POSITIVE | DMSO | 0.99999953 | 4.65E-07 | 0 |
| POSITIVE | DMSO | 0.99999955 | 4.51E-07 | 0 |
| POSITIVE | DMSO | 0.99999956 | 4.40E-07 | 0 |
| POSITIVE | DMSO | 0.99999956 | 4.39E-07 | 0 |
| POSITIVE | DMSO | 0.99999957 | 4.33E-07 | 0 |
| POSITIVE | DMSO | 0.99999957 | 4.32E-07 | 0 |
| POSITIVE | DMSO | 0.99999958 | 4.20E-07 | 0 |
| POSITIVE | DMSO | 0.99999959 | 4.09E-07 | 0 |
| POSITIVE | DMSO | 0.99999962 | 3.77E-07 | 0 |

| | | | | |
|---|---|---|---|---|
| POSITIVE | DMSO | 0.99999965 | 3.54E-07 | 0 |
| POSITIVE | DMSO | 0.99999965 | 3.53E-07 | 0 |
| POSITIVE | DMSO | 0.99999965 | 3.52E-07 | 0 |
| POSITIVE | DMSO | 0.99999965 | 3.48E-07 | 0 |
| POSITIVE | DMSO | 0.99999966 | 3.44E-07 | 0 |
| POSITIVE | DMSO | 0.99999966 | 3.39E-07 | 0 |
| POSITIVE | DMSO | 0.99999968 | 3.25E-07 | 0 |
| POSITIVE | DMSO | 0.99999968 | 3.24E-07 | 0 |
| POSITIVE | DMSO | 0.9999997 | 3.00E-07 | 0 |
| POSITIVE | DMSO | 0.9999997 | 2.99E-07 | 0 |
| POSITIVE | DMSO | 0.99999973 | 2.74E-07 | 0 |
| POSITIVE | DMSO | 0.99999974 | 2.56E-07 | 0 |
| POSITIVE | DMSO | 0.99999975 | 2.48E-07 | 0 |
| POSITIVE | DMSO | 0.99999976 | 2.44E-07 | 0 |
| POSITIVE | DMSO | 0.99999976 | 2.36E-07 | 0 |
| POSITIVE | DMSO | 0.99999977 | 2.30E-07 | 0 |
| POSITIVE | DMSO | 0.99999977 | 2.25E-07 | 0 |
| POSITIVE | DMSO | 0.99999979 | 2.15E-07 | 0 |
| POSITIVE | DMSO | 0.99999979 | 2.08E-07 | 0 |
| POSITIVE | DMSO | 0.99999979 | 2.07E-07 | 0 |
| POSITIVE | DMSO | 0.9999998 | 2.03E-07 | 0 |
| POSITIVE | DMSO | 0.99999982 | 1.85E-07 | 0 |
| POSITIVE | DMSO | 0.99999982 | 1.77E-07 | 0 |
| POSITIVE | DMSO | 0.99999983 | 1.73E-07 | 0 |
| POSITIVE | DMSO | 0.99999983 | 1.66E-07 | 0 |
| POSITIVE | DMSO | 0.99999984 | 1.64E-07 | 0 |
| POSITIVE | DMSO | 0.99999984 | 1.62E-07 | 0 |
| POSITIVE | DMSO | 0.99999984 | 1.60E-07 | 0 |
| POSITIVE | DMSO | 0.99999985 | 1.55E-07 | 0 |
| POSITIVE | DMSO | 0.99999985 | 1.48E-07 | 0 |
| POSITIVE | DMSO | 0.99999986 | 1.44E-07 | 0 |
| POSITIVE | DMSO | 0.99999986 | 1.39E-07 | 0 |
| POSITIVE | DMSO | 0.99999986 | 1.37E-07 | 0 |
| POSITIVE | DMSO | 0.99999986 | 1.37E-07 | 0 |
| POSITIVE | DMSO | 0.99999987 | 1.35E-07 | 0 |
| POSITIVE | DMSO | 0.99999988 | 1.24E-07 | 0 |
| POSITIVE | DMSO | 0.99999988 | 1.23E-07 | 0 |
| POSITIVE | DMSO | 0.99999988 | 1.19E-07 | 0 |
| POSITIVE | DMSO | 0.99999988 | 1.18E-07 | 0 |

| | | | | |
|---|---|---|---|---|
| POSITIVE | DMSO | 0.99999988 | 1.17E-07 | 0 |
| POSITIVE | DMSO | 0.99999992 | 8.19E-08 | 0 |
| POSITIVE | DMSO | 0.99999992 | 8.18E-08 | 0 |
| POSITIVE | DMSO | 1 | 0 | 0 |
| POSITIVE | DMSO | 1 | 0 | 0 |
| POSITIVE | DMSO | 1 | 0 | 0 |
| POSITIVE | DMSO | 1 | 0 | 0 |
| POSITIVE | DMSO | 1 | 0 | 0 |
| POSITIVE | DMSO | 1 | 0 | 0 |
| POSITIVE | DMSO | 1 | 0 | 0 |
| POSITIVE | DMSO | 1 | 0 | 0 |
| POSITIVE | DMSO | 1 | 0 | 0 |
| POSITIVE | DMSO | 1 | 0 | 0 |
| POSITIVE | DMSO | 1 | 0 | 0 |
| POSITIVE | DMSO | 1 | 0 | 0 |
| POSITIVE | DMSO | 1 | 0 | 0 |
| POSITIVE | DMSO | 1 | 0 | 0 |
| POSITIVE | DMSO | 1 | 0 | 0 |
| POSITIVE | DMSO | 1 | 0 | 0 |
| POSITIVE | DMSO | 1 | 0 | 0 |
| POSITIVE | DMSO | 1 | 0 | 0 |
| POSITIVE | DMSO | 1 | 0 | 0 |
| POSITIVE | DMSO | 1 | 0 | 0 |
| POSITIVE | DMSO | 1 | 0 | 0 |
| POSITIVE | DMSO | 1 | 0 | 0 |
| POSITIVE | DMSO | 1 | 0 | 0 |
| POSITIVE | DMSO | 1 | 0 | 0 |
| POSITIVE | DMSO | 1 | 0 | 0 |
| POSITIVE | DMSO | 1 | 0 | 0 |
| POSITIVE | DMSO | 1 | 0 | 0 |
| POSITIVE | DMSO | 1 | 0 | 0 |
| POSITIVE | DMSO | 1 | 0 | 0 |
| POSITIVE | DMSO | 1 | 0 | 0 |
| POSITIVE | DMSO | 1 | 0 | 0 |
| POSITIVE | DMSO | 1 | 0 | 0 |
| POSITIVE | DMSO | 1 | 0 | 0 |
| POSITIVE | DMSO | 1 | 0 | 0 |
| POSITIVE | DMSO | 1 | 0 | 0 |
| POSITIVE | DMSO | 1 | 0 | 0 |

| | | | | |
|---|---|---|---|---|
| POSITIVE | DMSO | 1 | 0 | 0 |
| POSITIVE | DMSO | 1 | 0 | 0 |
| POSITIVE | DMSO | 1 | 0 | 0 |
| POSITIVE | DMSO | 1 | 0 | 0 |
| POSITIVE | DMSO | 1 | 0 | 0 |
| POSITIVE | DMSO | 1 | 0 | 0 |
| POSITIVE | DMSO | 1 | 0 | 0 |
| POSITIVE | DMSO | 1 | 0 | 0 |
| POSITIVE | DMSO | 1 | 0 | 0 |
| POSITIVE | DMSO | 1 | 0 | 0 |
| POSITIVE | DMSO | 1 | 0 | 0 |
| POSITIVE | DMSO | 1 | 0 | 0 |
| POSITIVE | DMSO | 1 | 0 | 0 |
| POSITIVE | DMSO | 1 | 0 | 0 |
| POSITIVE | DMSO | 1 | 0 | 0 |
| POSITIVE | DMSO | 1 | 0 | 0 |
| POSITIVE | DMSO | 1 | 0 | 0 |
| POSITIVE | DMSO | 1 | 0 | 0 |
| POSITIVE | DMSO | 1 | 0 | 0 |
| POSITIVE | DMSO | 1 | 0 | 0 |
| POSITIVE | DMSO | 1 | 0 | 0 |
| POSITIVE | DMSO | 1 | 0 | 0 |
| POSITIVE | DMSO | 1 | 0 | 0 |
| POSITIVE | DMSO | 1 | 0 | 0 |
| POSITIVE | DMSO | 1 | 0 | 0 |
| POSITIVE | DMSO | 1 | 0 | 0 |
| POSITIVE | DMSO | 1 | 0 | 0 |
| POSITIVE | DMSO | 1 | 0 | 0 |
| POSITIVE | DMSO | 1 | 0 | 0 |
| POSITIVE | DMSO | 1 | 0 | 0 |
| POSITIVE | DMSO | 1 | 0 | 0 |
| POSITIVE | DMSO | 1 | 0 | 0 |
| POSITIVE | DMSO | 1 | 0 | 0 |
| POSITIVE | DMSO | 1 | 0 | 0 |
| POSITIVE | DMSO | 1 | 0 | 0 |
| POSITIVE | DMSO | 1 | 0 | 0 |
| POSITIVE | DMSO | 1 | 0 | 0 |
| POSITIVE | DMSO | 1 | 0 | 0 |
| POSITIVE | DMSO | 1 | 0 | 0 |

| | | | | |
|---|---|---|---|---|
| POSITIVE | DMSO | 1 | 0 | 0 |
| POSITIVE | DMSO | 1 | 0 | 0 |
| POSITIVE | DMSO | 1 | 0 | 0 |
| POSITIVE | DMSO | 1 | 0 | 0 |
| POSITIVE | DMSO | 1 | 0 | 0 |
| POSITIVE | DMSO | 1 | 0 | 0 |
| POSITIVE | DMSO | 1 | 0 | 0 |

**Supplementary Information Table 5 | Ranking PCs with highest power in the neural network classification model for selecting hits from the 330 compounds screened**

| Importance | Features |
|---|---|
| 100 | PCA12 |
| 65.1376629 | PCA07 |
| 60.54747033 | PCA02 |
| 60.18198402 | PCA05 |
| 58.23427369 | PCA01 |
| 57.79192763 | PCA11 |
| 53.95356952 | PCA08 |
| 43.44318034 | PCA48 |
| 43.43448467 | PCA34 |
| 43.39162756 | PCA31 |
| 41.37813617 | PCA10 |
| 41.02176346 | PCA15 |
| 40.77832799 | PCA41 |
| 40.47157613 | PCA03 |
| 39.11842555 | PCA16 |
| 37.72982538 | PCA42 |
| 37.5399662 | PCA04 |
| 34.42199848 | PCA18 |
| 33.1968855 | PCA09 |
| 32.04067388 | PCA21 |
| 27.68372093 | PCA20 |
| 27.68365951 | PCA37 |
| 25.44489243 | PCA40 |
| 25.13802912 | PCA49 |
| 23.93581621 | PCA06 |
| 22.89048908 | PCA17 |
| 22.76816728 | PCA13 |
| 22.60880899 | PCA23 |
| 21.22950563 | PCA29 |
| 19.95312176 | PCA35 |
| 18.85598276 | PCA28 |
| 18.02362965 | PCA19 |
| 17.92094649 | PCA25 |
| 17.6081184 | PCA24 |
| 17.42245276 | PCA47 |
| 14.9189612 | PCA38 |
| 13.87203804 | PCA30 |

| | |
|---|---|
| 13.60772636 | PCA46 |
| 12.77642976 | PCA14 |
| 12.76357655 | PCA33 |
| 12.73489226 | PCA45 |
| 11.55466193 | PCA44 |
| 10.74774516 | PCA39 |
| 10.1442005 | PCA36 |
| 9.281071306 | PCA22 |
| 6.954209204 | PCA26 |
| 6.336888663 | PCA50 |
| 5.552087165 | PCA27 |
| 2.808355951 | PCA32 |
| 0 | PCA43 |
